# Targeting Radiation-Induced Glioma-Initiating Cells in Patient-Derived Glioblastoma

**DOI:** 10.1101/2025.10.08.681196

**Authors:** Anjelica Cardenas, Sabrina Sutlief, Frank Pajonk

## Abstract

**Background:** Glioblastoma (GB) is a highly aggressive and treatment-resistant brain cancer with poor prognosis. Surgical resection followed by radiotherapy (RT) with the chemotherapeutic, temozolomide (TMZ), is the standard GB treatment; yet recurrence often occurs. GB is organized hierarchically with a small population of radiation-resistant glioma-initiating cells (GICs) that self-renew and drive tumor growth. Importantly, RT can induce a subset of cells from non-tumor-initiating into glioma-initiating cells (iGICs). Both GICs and iGICs contribute to tumor recurrence and therapy resistance. Thus, without effective elimination of non-tumorigenic GB and prevention or targeting of GICs, a cure is unlikely. The objective of this study is to identify small molecules that block RT-induced phenotypic conversion to occur.

**Method:** We conducted a high-throughput screen of NCI’s Cancer Therapy Evaluation Program (CTEP) compounds with evidence for crossing the blood-brain-barrier. To identify “stemness” or reprogramming of cells, we transduced GB cell lines representing each TCGA subtype to express a fluorescent reporter for proteasomal activity that distinguishes non–tumor-initiating cells from GICs. We tested CTEP agents at 10 different concentrations in combination with radiation.

**Results:** Our results identified selumetinib as a candidate compound that effectively prevents radiation-induced phenotype conversion. Furthermore, in combination with radiation, selumetinib decreased stem cell maintenance in GICs with differential effects on viability in non-tumorigenic cells.

**Conclusion:** Taken together, these findings suggest that repurposing FDA-approved compounds alongside current therapies may effectively target the cellular and molecular heterogeneity of GB—and because these agents are already clinically approved, this approach can be rapidly implemented in the clinic.

## INTRODUCTION

Glioblastoma (GB) is an aggressive brain cancer with a poor prognosis and median survival of 15 months ^1^. Standard treatment for GB is surgical resection followed by radiotherapy and temozolomide; however, improved therapies to extend survival and prevent progression remain elusive. GB is especially challenging because they are organized hierarchically, where cancer stem cells or glioma-initiating cells (GICs) self-renew and give rise to all lineages of differentiated cells ^2,3^. Furthermore, the ‘clonal evolution model’ suggests that non-initiating cells in a tumor can stochastically acquire a cancer stem cell phenotype as well. This conversion from non-tumorigenic to stem-like cells has been demonstrated under various stressors ^4^ including radiation in GB ^5,6^ and breast cancer ^7,8^. Therefore, data and the cancer stem cell hypothesis suggest that targeting GICs alone is not sufficient to cure GB ^9^. Instead, a combined effort of targeting GICs and tumor bulk as well as preventing phenotype conversion may provide improved therapeutic outcomes.

Radiation resistant tumor-initiating cells have been identified in breast cancer, cancer of the head and neck, and GB ^5,7,10-13^. It has been further reported that spontaneous and radiation-induced phenotype conversion of nontumorigenic cells into radioresistant GICs occurs via reprogramming through epigenetic changes. This phenomenon is not restricted to a specific TCGA subtype of GB, however, it is radiation dose-dependent and independent of p53, EGFRvIII, and MGMT status ^5^. While spontaneous conversion does take place, these events are more pronounced after irradiation ^5^. The induction of GICs by radiotherapy (RT) underscores the clinical relevance in targeting phenotype conversion with existing GICs to control the tumor.

Tumor-initiating cells (TICs) have been shown to have low proteasome activity. A fluorescent reporter system has been developed where cells with low proteasome activity are identified with the accumulation of the fluorescent protein, ZsGreen (ZsG+), fused to a degron that is normally recognized and eliminated by the 26S proteasome in solid tumors ^7,14-17^ including GB ^5,18^. From the National Cancer Institute (NCI) cancer therapy evaluation program (CTEP) portfolio, we selected compounds with evidence for crossing the blood-brain-barrier to test whether they prevent RT-induced conversion of non-GICs from proneural, classical, and mesenchymal GBM\ cell lines into iGICs. To account for off-target effects at higher drug concentrations, we conduced dose-response curves in conjunction with 3 radiation (Gy) doses. Once we identified effective compounds, we subsequently assessed combination effects on intrinsic GICs using limiting dilution assays.

## MATERIALS AND METHODS

### Cell Culture

All patient-derived glioma cell lines were established as previously described ^2^ at the University of California, Los Angeles (UCLA) and have been characterized ^19^. They were kindly provided by Dr. Harley I. Kornblum (UCLA). Primary cell lines were confirmed with DNA fingerprinting (Laragen, Culver City, CA, USA). ZsGreen (ZsG)-C-terminal degron of murine ornithine carboxylase (cODC) expressing cells were transduced with a lenti-viral reporter system as previously described ^18^. ZsG-cODC expressing cells were cultured in log-growth phase in Dulbecco’s modified Eagle’s medium (DMEM, Gibco, Waltham, MA, USA) with 10% fetal bovine serum (FBS, Sigma-Aldrich, St. Louis, MO, USA), penicillin (100 units/mL), and streptomycin (100 μg/mL). Primary glioma cells were propagated as gliomaspheres as previously described ^2^ in ultra-low adhesion plates in serum-free DMEM/F12 1:1 (Gibco) supplemented with SM1 (StemCell Technologies, Vancouver, Canada), epidermal growth factor (EGF, Sigma, Burlington, MA, USA), fibroblast growth factor 2 (bFGF, Sigma), and 0.68 U/mL heparin. All cells were grown in a humidified incubator at 37°C with 5% CO_2_ and routinely tested for mycoplasma via PCR (Applied Biological Materials, Vancouver, Canada).

### High-throughput screening assay

For screening of selected NCI CTEP compounds, ZsG-cODC-negative primary glioma cells were sorted by high-speed fluorescence activated cell sorting (BDFACSAria III, Becton Dickinson, Milpitas, CA). Low-evaporation 96-well plates (Greiner Bio-One, Kremsmunster, Austria) were filled by a manifold liquid dispenser (Multidrop Combi, Thermo Fisher, Waltham, MA) with media in each well (DMEM, 5% FBS, 1% penicillin-streptomycin). Using a liquid handler (Biomek FX, Beckman Coulter, Brea, CA) with a custom pin tool (V&P Scientific, San Diego, CA), 500 nL of each unique compounds in neat dimethyl sulfoxide (DMSO) were transferred from a compound source serial dilution plate to each 96-well plate according to predefined plate maps. The other wells received an equal volume of DMSO alone, serving as negative controls. Optimized number of cells (2,000-4,000) were then seeded into each well of the 96-well plates and incubated at 37°C, 5% CO_2_.

Twenty-four hours after plating, cells were irradiated (0, 4, or 8 Gy). Five days post-irradiation, Hoechst 33342 solution (25 μg/mL) was added to the cells. Following 1 hr incubation at 37°C, 5% CO_2_, plates were imaged on ImageXpress Micro Confocal (Molecular Devices, San Jose, CA) at 10× magnification, with 16 fields of view acquired per well. Laser scanning with a 488-nm laser was used for detection of cells expressing the fusion protein, ZsG-cODC. Additionally, scanning with an ultraviolet laser at 405 nm allowed for detection of Hoechst-stained nuclei, giving a measure of total viable cells. Images were acquired and cell quantification was performed in MetaXpress (Molecular Devices, San Jose, CA) for each field of view using automated segmentation and counting algorithms, for each fluorescence channel corresponding to the excitation lasers. Means of the counts for 16 images/well were used for determining viability and ZsG positive (+) to total viable cell ratios for each treatment condition. Ratios were normalized to the DMSO control, and values were subsequently transformed into Z-scores across all treatments, where more negative Z-scores reflect a greater reduction in the proportion of ZsG+ cells (iGICs) relative to controls.

### Limiting Dilution Assays

Patient-derived GB cells were seeded under serum-free condition at clonal densities into non-treated tissue culture plates. The next day, cells were treated with compounds and irradiated 1h later using an experimental X-ray irradiator (Gulmay Medical Inc.). Cells were grown for 7-14 days and supplemented with 10 μl/well of media with growth factors every other day. Upon formation of visibly distinct spheres, spheres were counted in each well using a conventional microscope and recorded. For sphere formation capacity (SFC), sphere formation at each dose was normalized against the non-irradiated control. Data points were fitted using a linear quadratic model. Limiting dilutions were calculated and plotted using Extreme Limiting Dilution Analysis software ^20^.

### Drug Treatment

A full list of CTEP compounds and vendors can be found in **Supplementary Table 1**. We selected compounds on the current agreement at the start of this study (12/15/2021), however, the CTEP portfolio is updated every six months. Some of the evaluated compounds are no longer on the agreement and newer agents have not been evaluated.

### Irradiation

Cells were irradiated at room temperature with an x-ray irradiator at a dose rate of 5.519 Gy/min for the calculated time necessary to apply prescribed dose. The X-ray beam was operated at 300 kV and hardened using a 4-mm Be, a 3-mm Al, and a 1.5 mm Cu filter and calibrated using National Institute of Standards and Technology-traceable dosimetry. Corresponding controls were sham irradiated.

### Statistical Analysis

Graphpad Prism Software (Graphpad Software, San Diego, CA) was used to perform statistical analyses. For sphere forming assays, Extreme Limiting Dilution Analysis software was used to calculate limiting dilutions ^20^. Synergy was quantified using the ZIP model in SynergyFinder+ ^21^. A ZIP-score ≥ 10 is considered strongly synergistic and ≤ -10 strongly antagonistic.

## RESULTS

### High-throughput screen of CTEP portfolio compounds alone and in combination with radiation in patient-derived GB

To identify compounds that prevent RT-induced conversion of non-tumorigenic glioma cells into induced glioma-initiating cells (iGICs), we leveraged a fluorescent reporter system and performed a high-throughput (HT) dose-response screen (**Figure 1A**). We selected a panel of small molecule compounds from the NCI CTEP portfolio predicted to or with published evidence for crossing the blood-brain-barrier (BBB, **Table 1**). BBB permeability was determined by a LogBB permeability predictor server with the cutoff of logBB value >= -1 ^22^.

**Table 1.** Predicted and published BBB Permeability.

| Compound | LogBB Value | Predicted BBB Permeability | Published evidence |
| --- | --- | --- | --- |
| Alisertib | -1.1062 | No | 43,44 |
| AMG337 | -0.58253 | Yes |  |
| Azacitidine | -0.74865 | Yes |  |
| AZD 1775 | -0.83637 | Yes |  |
| AZD 8186 | -0.32544 | Yes |  |
| AZD 9291 | -1.02864 | No | 45,46 |
| Belinostat | -0.93999 | Yes |  |
| Birinapant | -0.9818 | Yes |  |
| Bortezomib | -0.93766 | Yes |  |
| Cabozantinib | -0.48749 | Yes |  |
| Copanlisib | -0.68312 | Yes |  |
| Dabrafenib mesylate | -0.91475 | Yes |  |
| Dasatinib | -1.00632 | No | 47 |
| Entinostat | -0.96386 | Yes |  |
| Eribulin | -0.33528 | Yes |  |
| Erlotinib | -0.1436 | Yes |  |
| Ibrutinib | -0.60372 | Yes |  |
| INK128 | -0.62164 | Yes |  |
| Lapatinib | -0.11286 | Yes |  |
| Lenalidomide | -0.67264 | Yes |  |
| Methoxyamine HCl | 0.03146 | Yes |  |
| Olaparib | -0.41314 | Yes |  |
| Onalespib lactate | -0.54827 | Yes |  |
| Pazopanib | -0.87894 | Yes |  |
| Pomalidomide | -0.57043 | Yes |  |
| Rociletinib | -0.54702 | Yes |  |
| Romidepsin | -0.68971 | Yes |  |
| Selumetinib | -0.54623 | Yes |  |
| Sorafenib tosylate | -0.91091 | Yes |  |
| Sunitinib malate | -0.73894 | Yes |  |
| Talazoparib | -0.55141 | Yes |  |
| Tazemetostat | -0.72856 | Yes |  |
| Temsirolimus | -0.53552 | Yes |  |
| Tivantinib | -0.55436 | Yes |  |
| Triapine | -0.45847 | Yes |  |
| Veliparib | -0.52999 | Yes |  |
| Vismodegib | -0.65919 | Yes |  |

**Figure 1.**
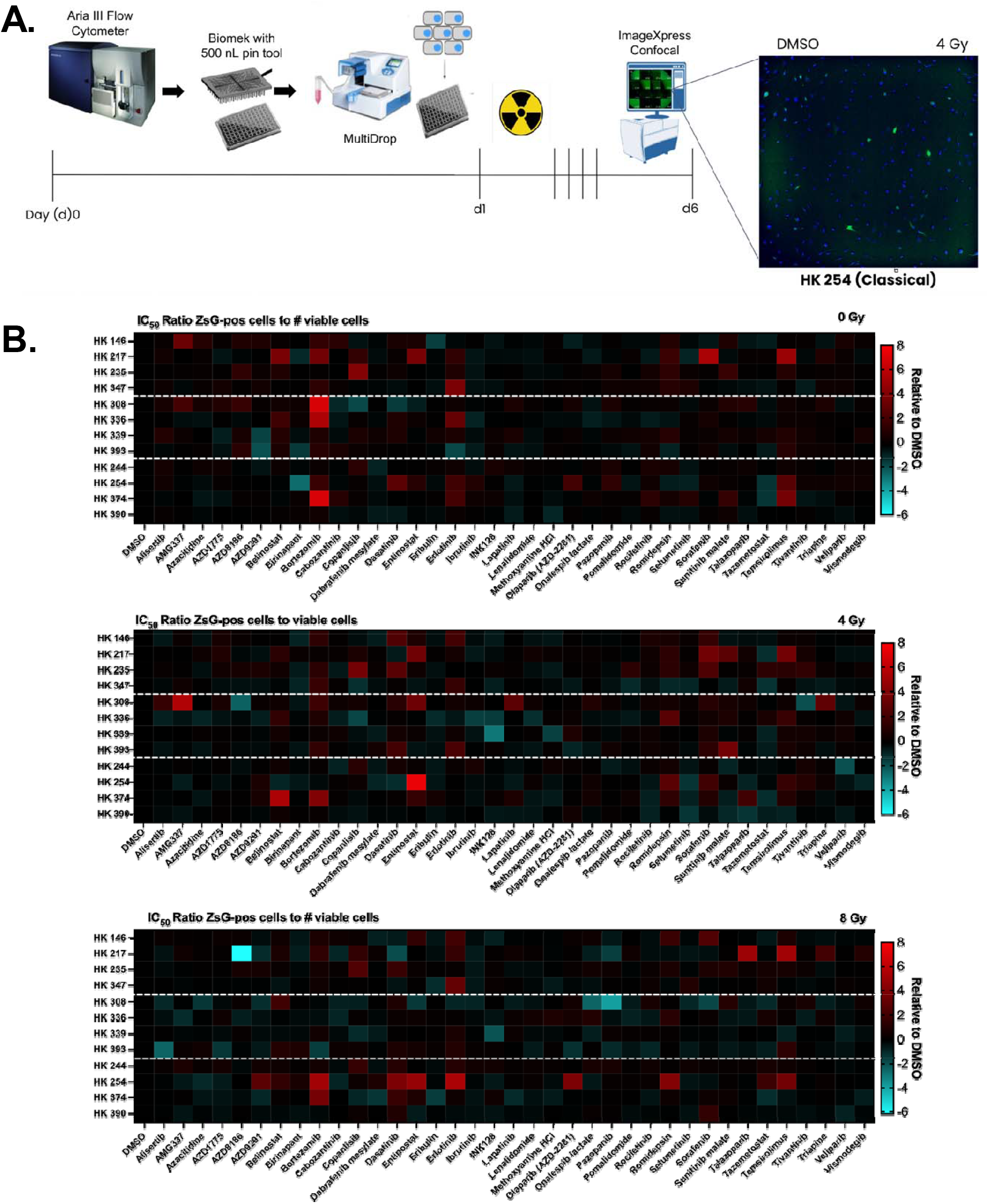
High-throughput screen of CTEP portfolio compounds alone and in combination with radiation in patient-derived GB. A. Schematic of high-throughput assay workflow. B. Heatmaps of ZsG+/total viable cell ratios at the approximate IC_50_ concentration calculated for each compound and cell line at 0 Gy (top), 4 Gy (middle), and 8 Gy (bottom). Each cell in the heatmap represents log2-transformed ratios from average triplicate data (compound vs. DMSO control), normalized as a z-score across compounds to highlight relative changes. Cell lines are grouped by TCGA subtype, top: proneural (HK 157, HK 217, HK 235, and HK 347), middle: mesenchymal (HK 308, HK 336, HK 339, HK 393), and bottom: classical (HK 244, HK 254, HK 374, HK 390).

**Figure 2.**
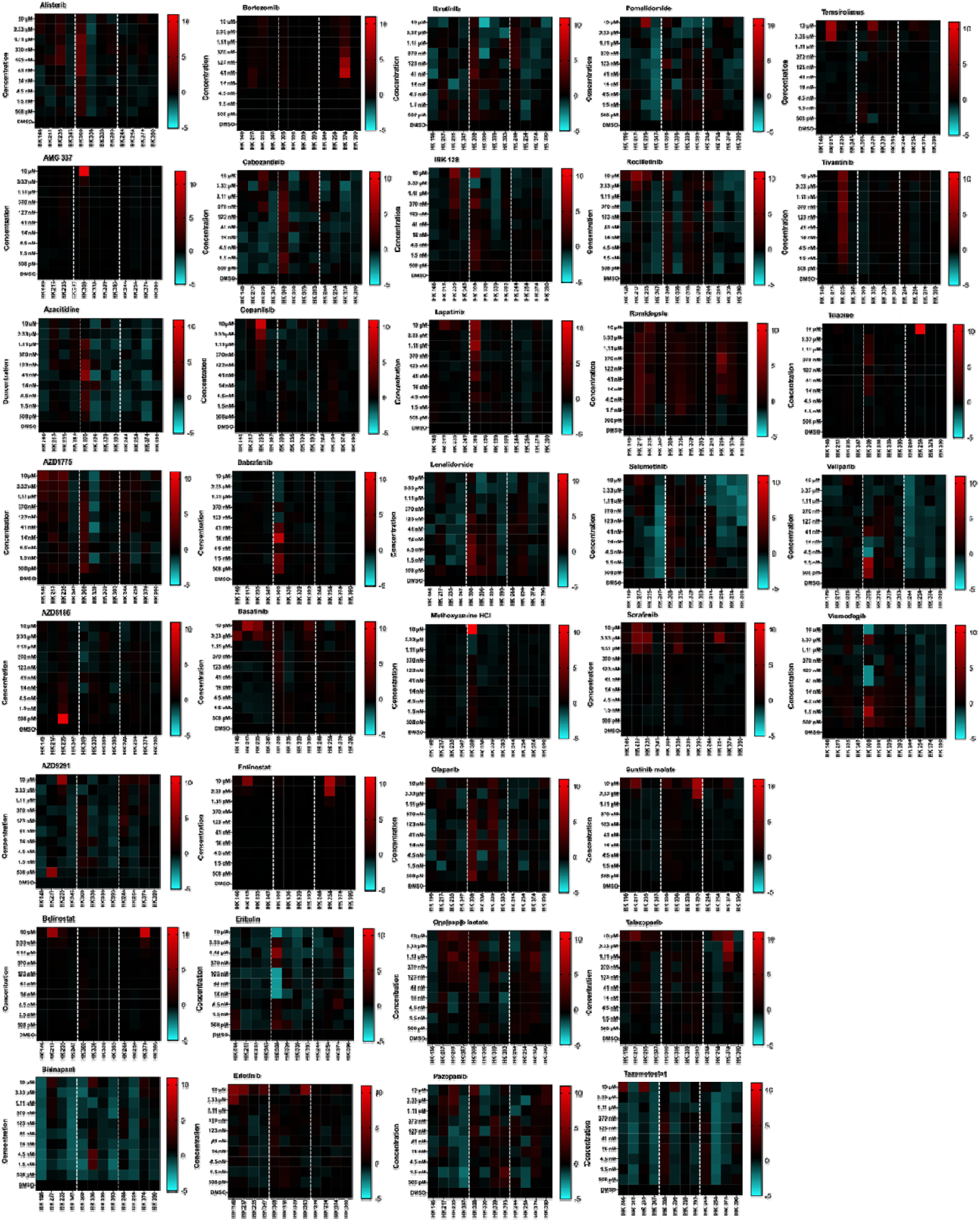
Hit identification: selumetinib prevents radiation-induced reprogramming at 4 Gy in classical GB cell lines. Heatmaps of ZsG+/total viable cell ratio across compounds were quantified for all compounds tested at 10 concentrations (508 pM–10 μM) across 12 cell lines following 4 Gy irradiation. Values represent the mean ZsG+/total viable cell ratio from three technical replicates per condition (compound vs. DMSO control), normalized as a z-score across cell lines to highlight relative changes. Heat maps display relative changes in GIC induction relative to DMSO control. Cell lines are grouped by TCGA subtype, left: proneural (HK 157, HK 217, HK 235, and HK 347), middle: mesenchymal (HK 308, HK 336, HK 339, HK 393), and right: classical (HK 244, HK 254, HK 374, HK 390).

Using a fluorescent reporter system to identify TICs based on lack of proteasome activity, we transduced 12 patient-derived GB cell lines, representing three TCGA subtypes (proneural, mesenchymal, and classical) with various genetic mutations (**Table 2**). For the HTS, cells were sorted for negative ZsG expression (non-tumorigenic), plated, and treated with CTEP drugs. We screened each drug at 10 concentrations ranging from 508 pM-10 μM. Cells were then irradiated (0, 4 and 8 Gy) the next day. Five days post-irradiation, Hoechst was added to each well and plates were imaged. Images were analyzed to determine cell viability and radiation-induced phenotype conversion (**Figure 1A**). Percent viability was determined relative to DMSO control for each plate. Viability heatmaps of compounds screened at each concentration and radiation dose for all cell lines can be found in **Supplementary Figures 1-3**. Further, dose response curves and IC_50_ values were generated for all experimental groups (**Supplementary Figures 4-40)**.

**Table 2.** Patient demographics and TCGA-classification of GB subtypes.

| Line | Origin | TCGA subtype | Culture P53 CN | EGFRvIII | PTEN | MGMT |
| --- | --- | --- | --- | --- | --- | --- |
| HK 146 | Recurrent GBM | p | Unknown | Unknown | Unknown | Unknown |
| HK 217 | Primary GBM | p | wt | Negative | Negative | Unknown |
| HK 235 | PNET Grade IV | p | Unknown | Negative | Positive | Unknown |
| HK 347 | Recurrent GBM | p | wt | Unknown | Unknown | Unknown |
| HK 308 | Recurrent GBM | m | Unknown | Positive | Positive | not methylated |
| HK 336 | Recurrent GBM | m | CN loss | Negative | Positive | methylated |
| HK 339 | Recurrent GBM | m | Unknown | Unknown | Unknown | Unknown |
| HK 393 | Recurrent GBM | m | Unknown | Unknown | Unknown | Unknown |
| HK 244 | Primary GBM | c | Unknown | Negative | Positive | not methylated |
| HK 254 | Recurrent GBM | c | Unknown | Unknown | Unknown | Unknown |
| HK 374 | Primary GBM | c | Loss "mosaic" | Positive | Positive | not methylated |
| HK 390 | Primary GBM | c | wt | Negative | Positive | not methylated |

**Abbreviations for TCGA subtypes: c: classical; m: mesenchymal; p: proneural.**

Radiation-induced cellular reprogramming was determined by ZsG+ to total viable cell ratio relative to DMSO control. To summarize HTS results, ZsG+/total viable cell ratio at the most biologically relevant concentration (approximate IC_50_) for each compound were plotted in a heatmap for each radiation dose (**Figure 1B**). If IC_50_ values were undetermined/greater than 10 μM or less that 508 pM, we used those doses. Two-way ANOVA showed a main effect of CTEP compound for all radiation doses (**Supplementary Table 2**). At 8 Gy, a main effect of TCGA subtype was identified, however, at 0 and 4 Gy, no main effect of TCGA subtype was observed (**Supplementary Table 2**). For both 4 and 8 Gy, no significant interaction between CTEP compounds and TCGA subtypes were detected (**Supplementary Table 2**). Of note, none of the tested compounds showed significant reduction in radiation-induced phenotype conversion (ZsG+/total viable cell ratio) compared to DMSO control.

### Hit Identification: Selumetinib prevents radiation-induced reprogramming

For spontaneous reprogramming (0 Gy), using a mixed-effects model (REML), AZD 1775, AZD 9291, belinostat, bortezomib, cabozantinib, and suntinib malate showed a main effect for concentration while entinostat and ibrutinib showed a main effect for TCGA subtype (**Supplementary Figure 41, Supplementary Table 2**). Post-hoc analysis only identified significance for mesenchymal GB for 123 nM belinostat treatment (Dunnett’s Adjusted P-value = 0.0358), 1.5 nM cabozantinib treatment in classical GB (Dunnett’s Adjusted P-value = 0.0131), and 14 nM entinostat treatment in classical GB (Dunnett’s Adjusted P-value = 0.0008). Of the CTEP compounds tested for radiation-induced reprogramming, dasatinib and erlotinib at 4 Gy and bortezomib, dasatinib, and suntinib malate at 8 Gy exhibited a main effect of concentration while romidepsin at 8 Gy showed a main effect for TCGA subtype (**Figures 3-4, Supplementary Tables 3-4**). No comparisons survived post-hoc analyses for dasatinib at 4 Gy or bortezomib at 8 Gy. Post-hoc analysis identified significance in proneural cell lines treated with 1.5 nM erlotinib at 4 Gy (Dunnett’s Adjusted P-value = 0.0183) and for classical cell lines treated with 10 μM dasatinib (Dunnett’s Adjusted P-value = 0.0411). All other compounds except for selumetinib were not statistically significant for any main or interactive effects.

**Figure 3.**
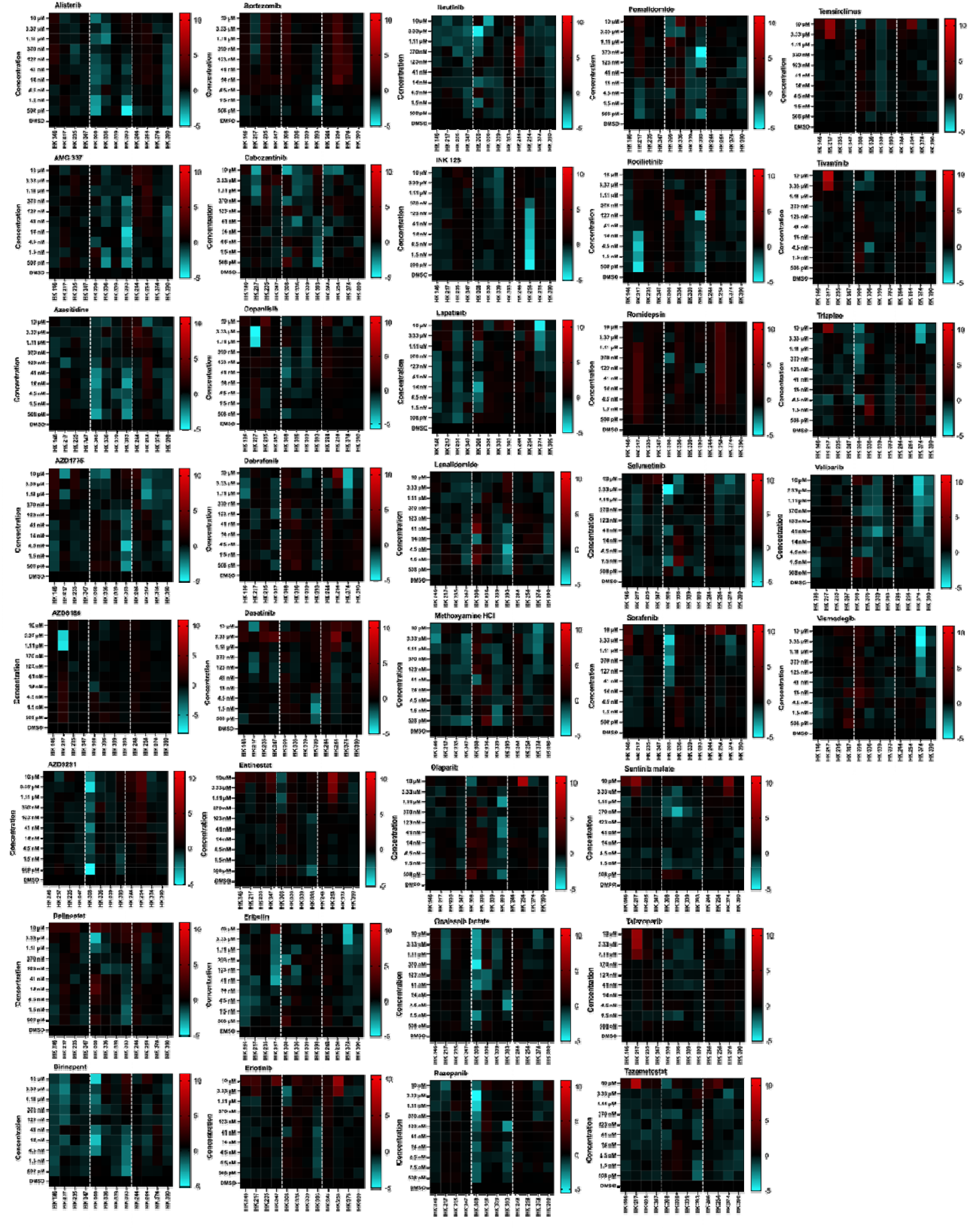
Hit identification: no compound was identified to prevent radiation-induced reprogramming at 8 Gy in patient-derived GB cell lines. Heatmaps of ZsG+/total viable cell ratio across compounds were quantified for all compounds tested at 10 concentrations (508 pM–10 μM) across 12 cell lines following 8 Gy irradiation. Values represent the mean ZsG+/total viable cell ratio from three technical replicates per condition (compound vs. DMSO control), normalized as a z-score across cell lines to highlight relative changes. Heat maps display relative changes in GIC induction relative to DMSO control. Cell lines are grouped by TCGA subtype, left: proneural, middle: mesenchymal, and right: classical.

Mixed-effects model of selumetinib, a MEK inhibitor, revealed an interactive effect for concentration x TCGA subtype with a trend for TCGA subtype at 4 Gy (**Figure 4, Supplementary Table 4**). No significant effects were observed at 0 Gy (spontaneous phenotype conversion) or 8 Gy (**Supplementary Tables 2 and 5)**. Post-hoc analysis identified significance for classical cell lines at 1.11 and 10 μM concentrations of selumetinib (**Supplementary Table 6**). To determine if observed effects of selumetinib with radiation was synergistic, we compared ZsG+/total viable cell ratio relative to DMSO with radiation and did not observe synergy in any cell lines tested **(Supplementary Figures 42)**.

**Figure 4.**
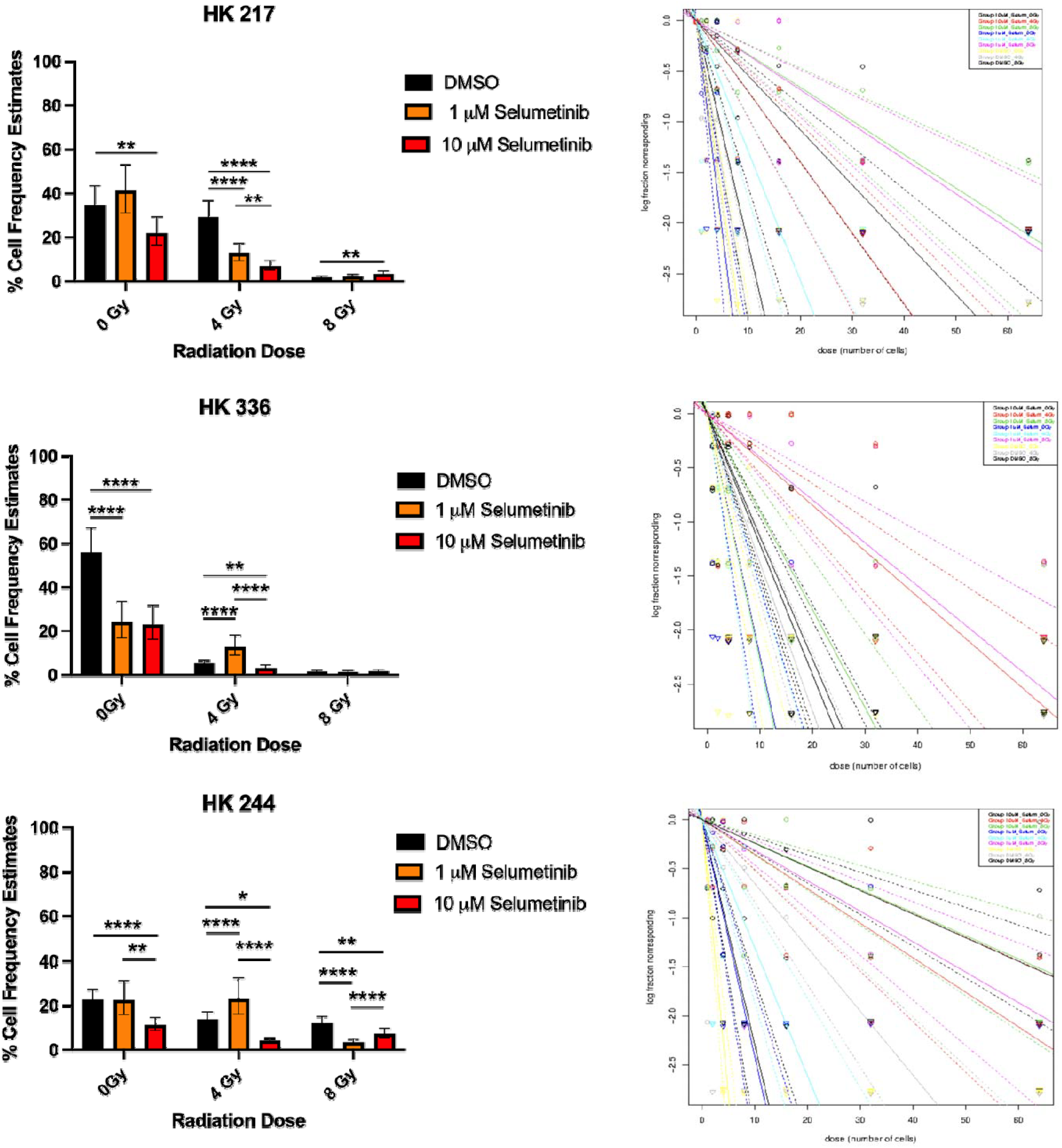
HIT validation: selumetinib reduces stem cell frequency in a dose- and radiation-dependent manner. Left: Bar graphs of percent (%) cell frequency estimates as calculated by extreme limiting dilution analysis (ELDA)^2^ for Proneural (HK 217, top), Mesenchymal (HK 336, middle), and Classical (HK 244, bottom) patient-derived gliomaspheres, n=3-8/group. Right: Log-fraction plots of the limiting dilution model fitted to the data. Slope of the line is the log-active cell fraction. Data are shown as mean ± 95% confidence intervals.

Two-way repeated measures ANOVA revealed a significant main effect of concentration at both 0 Gy (F (2.177, 6.531) = 14.86, p = 0.0034) and 4 Gy (F (2.346, 7.037) = 8.877, p = 0.0105), indicating dose-dependent reductions in viability with selumetinib. At 0 Gy, classical cell lines showed significantly reduced viability at the two highest concentrations relative to DMSO, whereas pairwise comparisons at 4 Gy did not remain significant after post-hoc correction. No main effect of TCGA subtype or concentration × subtype interaction was observed at either radiation dose, suggesting no subtype-specific sensitivity to selumetinib. At 8 Gy, no main effects or interactions were detected, indicating that selumetinib did not alter viability at higher radiation doses. Taken together, these findings suggest that reduction in radiation-induced reprogramming observed with selumetinib is not attributed to cell death at higher doses of radiation.

### Hit validation: Selumetinib reduces stem cell maintenance

To investigate whether the selected compound, selumetinib, impact GICs, we assessed their effects alone and in combination with radiation on self-renewal capacity *in vitro*. Using functional limited dilution assays, in three cell lines that represent each TCGA type (Classical HK244; Proneural HK 217; Mesencymal HK 336), combined selumetinib with radiation treatment significantly decreased stem cell frequencies in all cell lines (**Figure 4). Supplementary Tables 5-6** show confidence intervals for 1/(stem cell frequency and overall test for differences in stem cell frequencies between any of the groups calculated via ELDA webtool. Our findings suggest that in addition to preventing radiation-induced phenotype conversion, selumetinib combined with radiation treatment reduced stem cell maintenance in existing GIC populations.

## DISCUSSION

RT for GB has reached a therapeutic plateau, with no improvements observed in decades ^23^, therefore, effective treatments are urgently needed to extend survival and prevent tumor recurrence. To address cellular heterogeneity and plasticity in GB with the overall goal of improving RT outcome, we sought to identify compounds capable of preventing radiation-induced conversion of bulk tumor cells into iGICs. Our HTS of CTEP compounds that cross the BBB revealed selemetinib limits radiation-induced reprogramming in patient-derived GB for the classical TCGA subtype. Functional assays demonstrate reduced stem cell frequency, consistent with impaired self-renewal via combined radiation and selumetinib treatment. Importantly, the overall lack of efficacy across compounds tested at 10 concentrations, both with and without radiation, underscores the major challenge in identifying drugs that can work with RT to simultaneously target bulk tumor cells, radiation-induced reprogramming, and intrinsic GICs.

In this study, we utilized an imaging system developed to track the development of radiation-induced phenotype conversion of non-tumorigenic GB cells into GICs ^18^ based on lack of proteasome activity in TICs ^18^. While our reporter system relies on the 26S proteasome, observed effects of some compounds may be due to proteasome (bortezomib) or proteostasis inhibition (onalespib lactate) and not radiation-induced conversion. However, while bortezomib did exhibit main effects, there were no identified effects in post-hoc analysis or any significant effects for onalepib lactate. Bortezomib also served as a positive control because it is a proteasome inhibitor that has not been successful in the clinical setting for solid tumors ^24-28^. Our candidate compound, selumetinib, likely doesn’t impact the proteasome since it is a specific inhibitor of MEK. Nonetheless, we followed up HTS assay with limiting dilution assays, which do not rely on the proteasome, to validate our findings and show that selumetinib dose- and radiation-dependently reduces stem cell maintenance.

Selumetinib is approved for pediatric plexiform neurofibromas and low-grade gliomas. Few studies have investigated selumetinib in GB ^29-31^ and none could be located where selumetinib is combined with radiation. Although, MEK inhibition has been shown to be effective in BRAF mutant and neurofibromin type 1 (NF1)-deficient GB ^32^, it is not long-lasting ^33-35^. Furthermore these MEK inhibitors are not predicted to cross the BBB (LogBB = -1.018) ^22^ with evidence of limited penetration due to efflux pumps ^36^. Classical GB do not typically contain BRAF and NF1 alterations. Yet, in our study, classical cell lines are the most affected by selumetinib in preventing radiation-induced reprogramming. This suggest that selumetinib may be working through another mechanism. Furthermore, since we did not observe synergy with radiation, we can infer that selumetinib is working independently of ionizing radiation as well.

Targeted therapy combined with TMZ and radiotherapy in clinical trials against dysregulated GB targets (EGFR, mTOR, PI3K) have been unsuccessful in improving overall survival in GB patients, despite preclinical data support ^37,38^. Many clinical trials have included agents used in this study. Although preclinical data suggest dasatanib with radiation is efficacious, a clinical trial found that dasatinib combined with standard chemo-radiation treatment did not improve outcome in patients ^39^. Another study demonstrated that ibrutinib alone reduced stem cell maintenance but did not affect neural stem cells and in combination with radiation, ibrutinib increased overall survival *in vivo* ^40^. Phase I clinical trials for the use of ibrutinib in combination with TMZ and radiation in GB is currently underway ^41^. However, we did not observe significant reduction of radiation-induced reprogramming in our HTS for ibrutinib.

On the other hand, selumetinib has not been extensively studied in GB. Before selumetinib can be transitioned to the clinic for GB, additional studies are necessary to evaluate the effects of selumetinib alone and in combination with radiation in both orthotopic models and immunocompetent systems. Alternatively, a Human Organoid Tumor Transplantation (HOTT) translational *ex vivo* model for GB can be utilized ^42^.

Moreover, the mechanisms underlying our observed effects need to be investigated. Additional studies can evaluate the expression of “stem” markers such as SOX2 and OLIG2 to validate loss of stem cell maintenance and prevention of radiation-induced reprogramming. Bulk and/or single-cell RNA sequencing should be pursued comparing radiation vs combined radiation and selumetinib treatment to detect reprogramming signatures and cell-state shifts as well as identify altered pathways followed-up by mechanistic perturbations of targets of interest and determine whether effects are lost or gained. Furthermore, while we used a limited number of patient-derived cell lines to represent each TCGA subtype, it would be good to evaluate effects in additional classical GB cell line and limit generality. Proposed future experiments are required to determine whether selumetinib is a candidate compound to advance forward as a rational combination therapy with RT to treat GB and prevent recurrence in classical GB subtypes.

## Conclusion

Overall, our HTS revealed that few already FDA-approved compounds effectively prevent radiation-induced phenotype conversion in GB, underscoring the difficulty of identifying small molecules that cross the BBB and impact both non-tumorigenic and glioma-initiating cells. Importantly, our HTS findings with functional stemness assays suggests selumetinib specifically targets the tumor-initiating population, offering potential to reduce recurrence and improve therapeutic response that could be rapidly implemented into the clinic. While limitations remain regarding dosing and translation to *in vivo* models, these findings highlight how drug screening can inform clinical care and accelerate the repurposing of existing therapies for GB.

## Author Contributions

The concept was conceived by FP. FP and AC contributed to experimental design. AC transfected cell lines and performed most experiments. SS contributed to experiments. Analyses were done by AC. AC and FP wrote the manuscript.

## Acknowledgements

We would like to thank Jeremy Karkafi and Sophia Tate for data management as well as Carter Hoffman and Carlos Calderon for their laboratory assistance. We would also like to thank the UCLA Molecular Screening Shared Resource and Jonsson Comprehensive Cancer Center (JCCC) and Center for AIDS Research Flow Cytometry Core personnel. The flow cytometry core facility is supported by National Institutes of Health awards P30 CA016042 and 5P30 AI028697, and by the JCCC, the UCLA AIDS Institute, the David Geffen School of Medicine at UCLA, the UCLA Chancellor’s Office, and the UCLA Vice Chancellor’s Office of Research.

