## Supplementary Figure 1 for "Targeting Radiation-Induced Glioma-Initiating Cells in Patient-Derived Glioblastoma"

### Slide 1
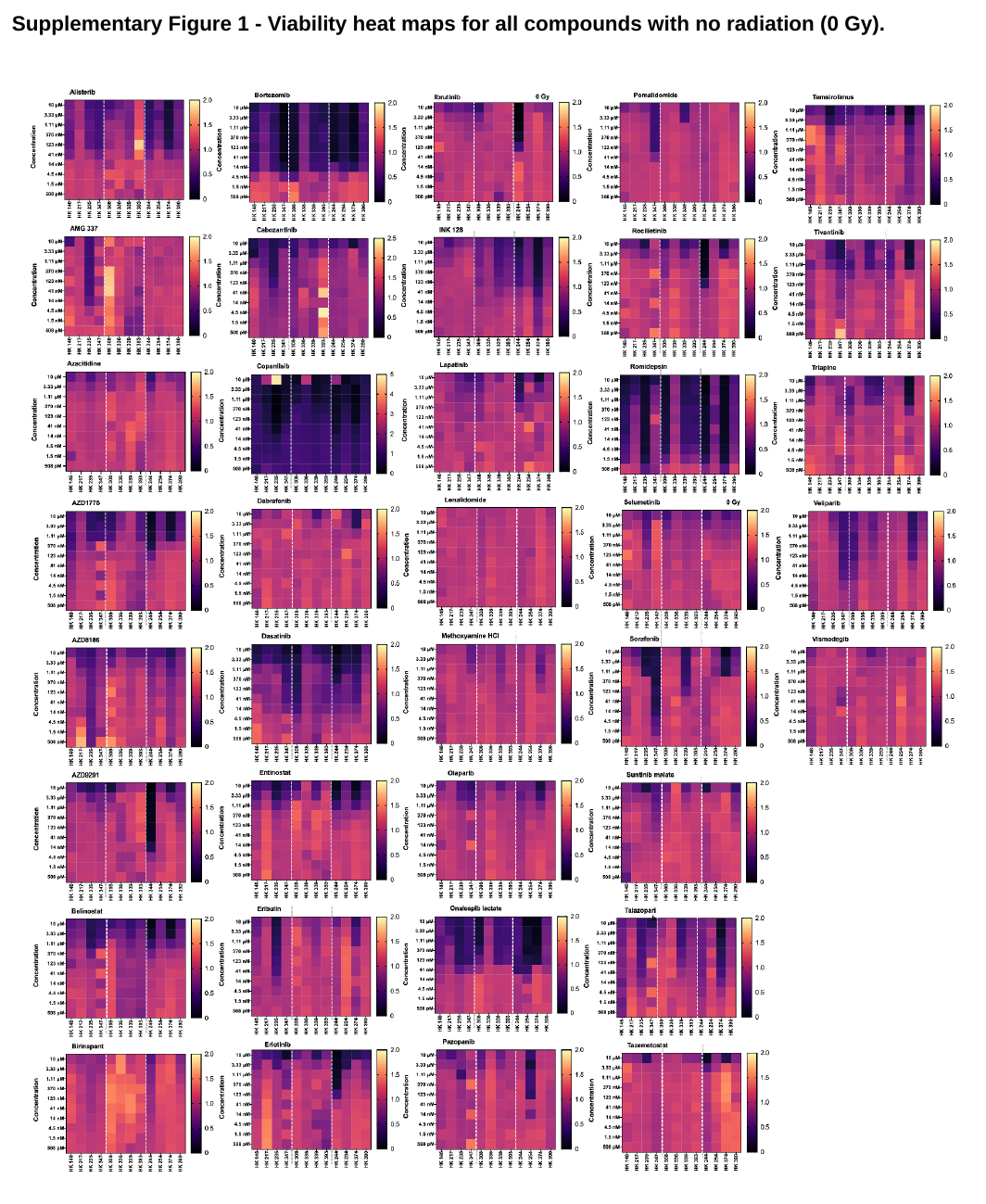

#### Supplementary Figure 1 - Viability heat maps for all compounds with no radiation (0 Gy).

### Slide 2
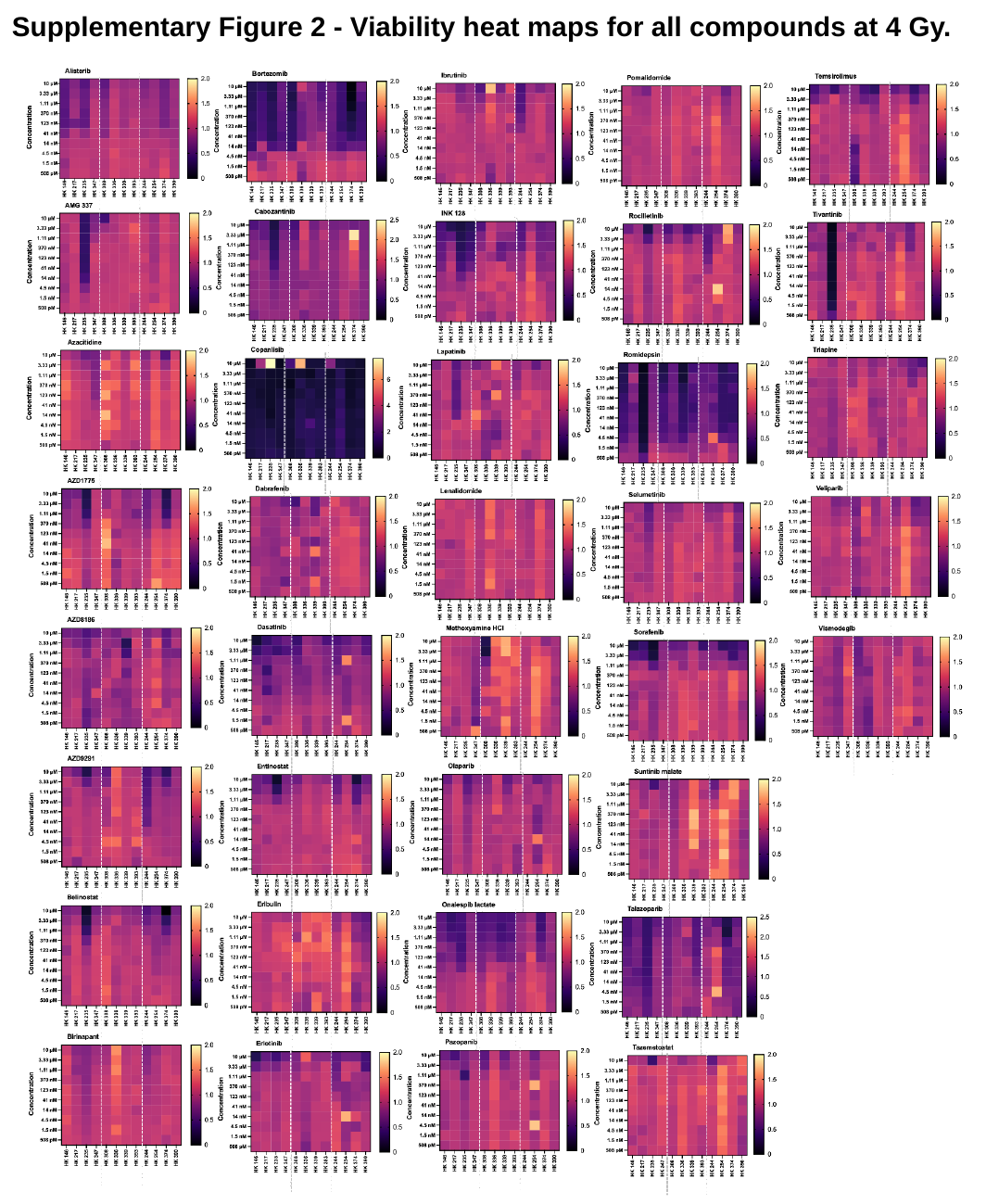

#### Supplementary Figure 2 - Viability heat maps for all compounds at 4 Gy.

### Slide 3
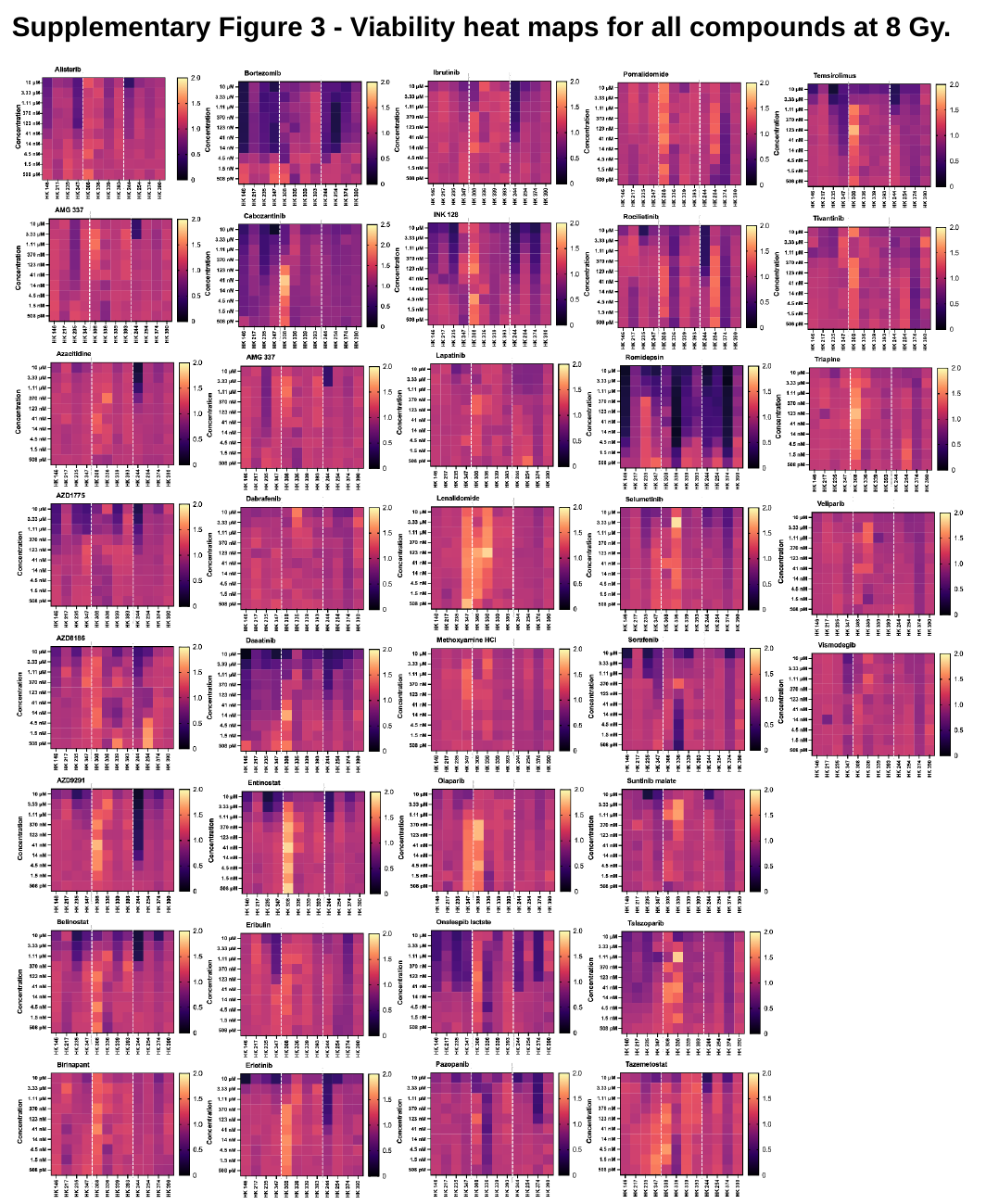

#### Supplementary Figure 3 - Viability heat maps for all compounds at 8 Gy.

### Slide 4
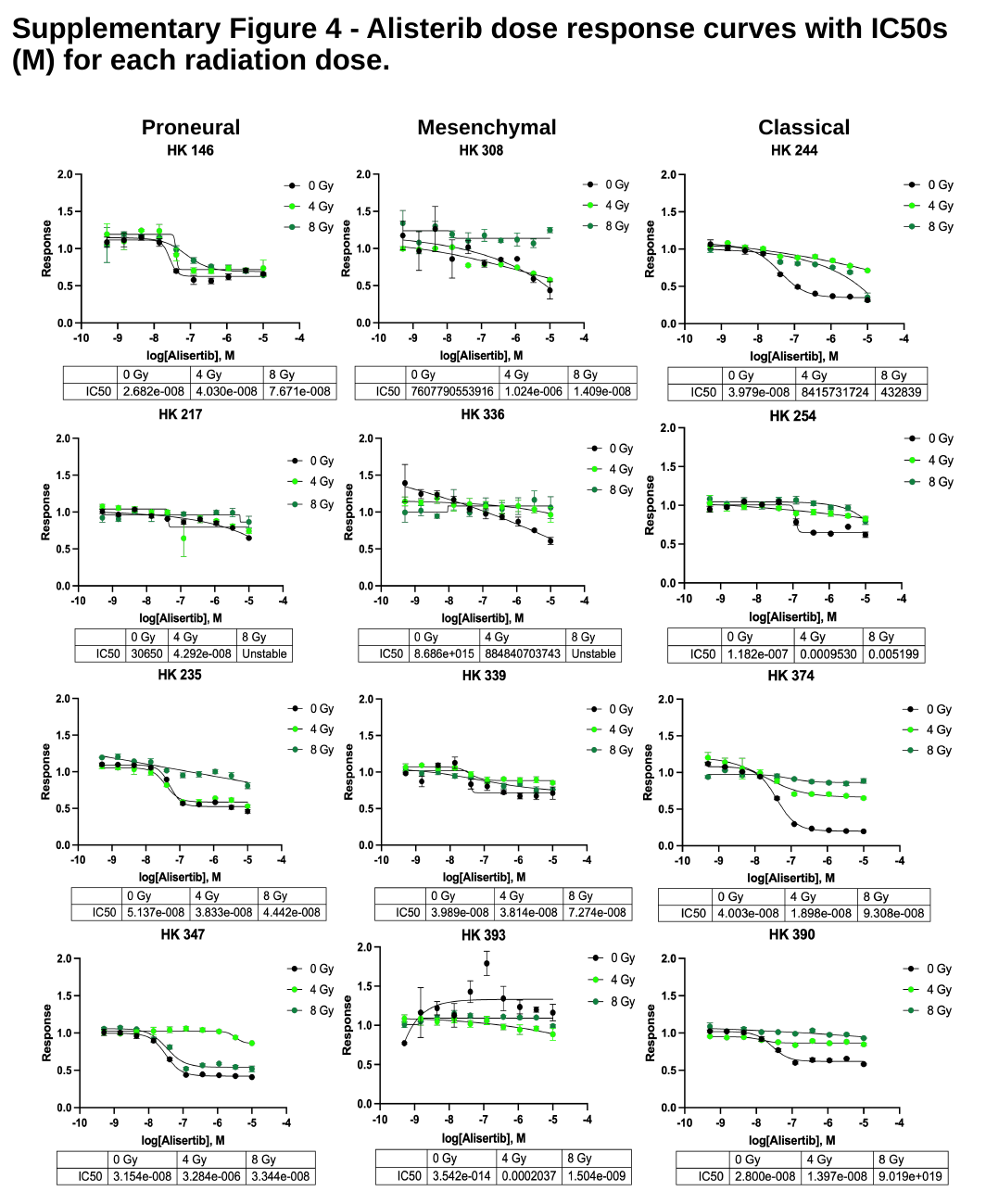

#### Supplementary Figure 4 - Alisterib dose response curves with IC50s (M) for each radiation dose.
Classical
Proneural
Mesenchymal

### Slide 5
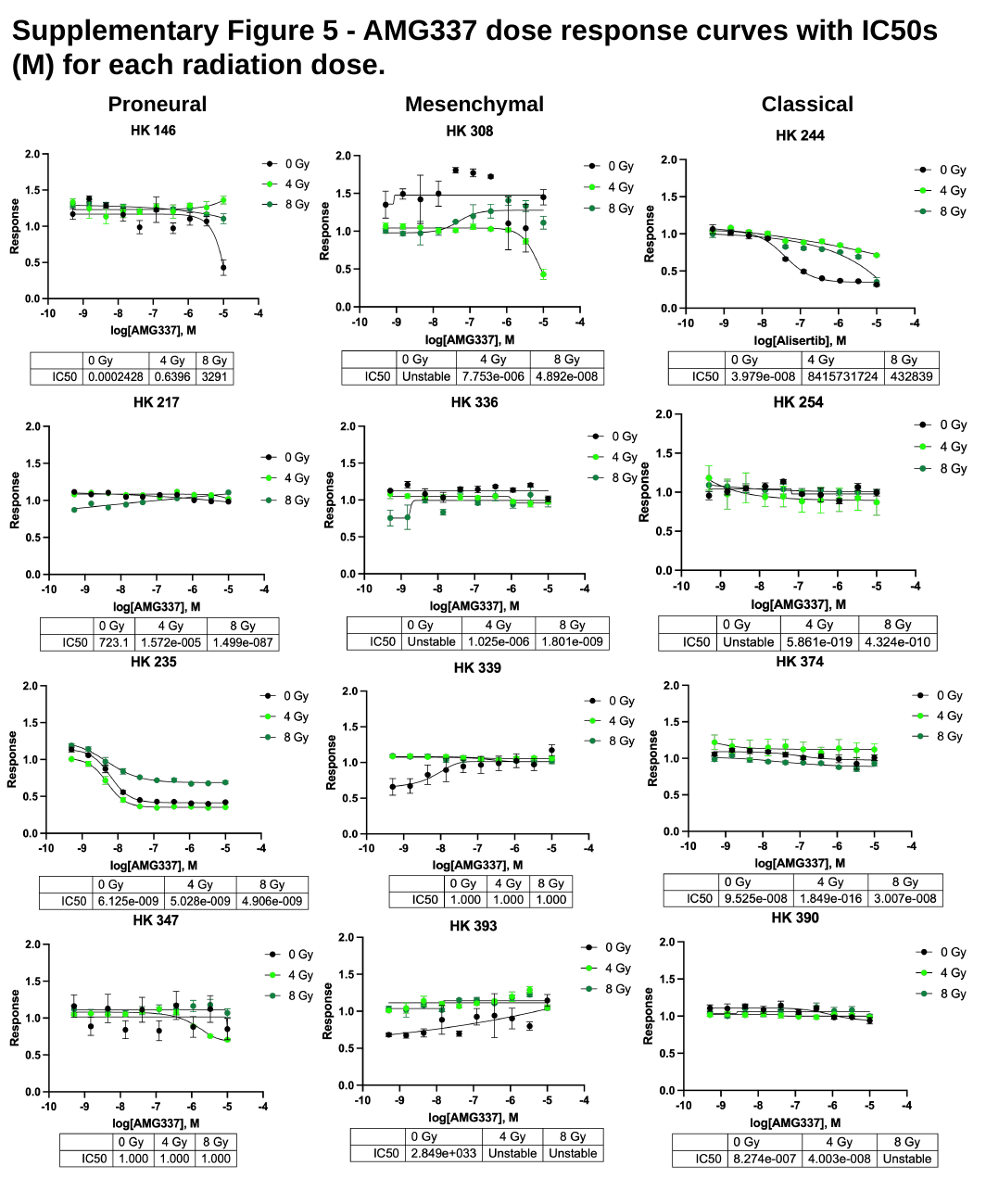

#### Supplementary Figure 5 - AMG337 dose response curves with IC50s (M) for each radiation dose.
Proneural
Classical
Mesenchymal

### Slide 6
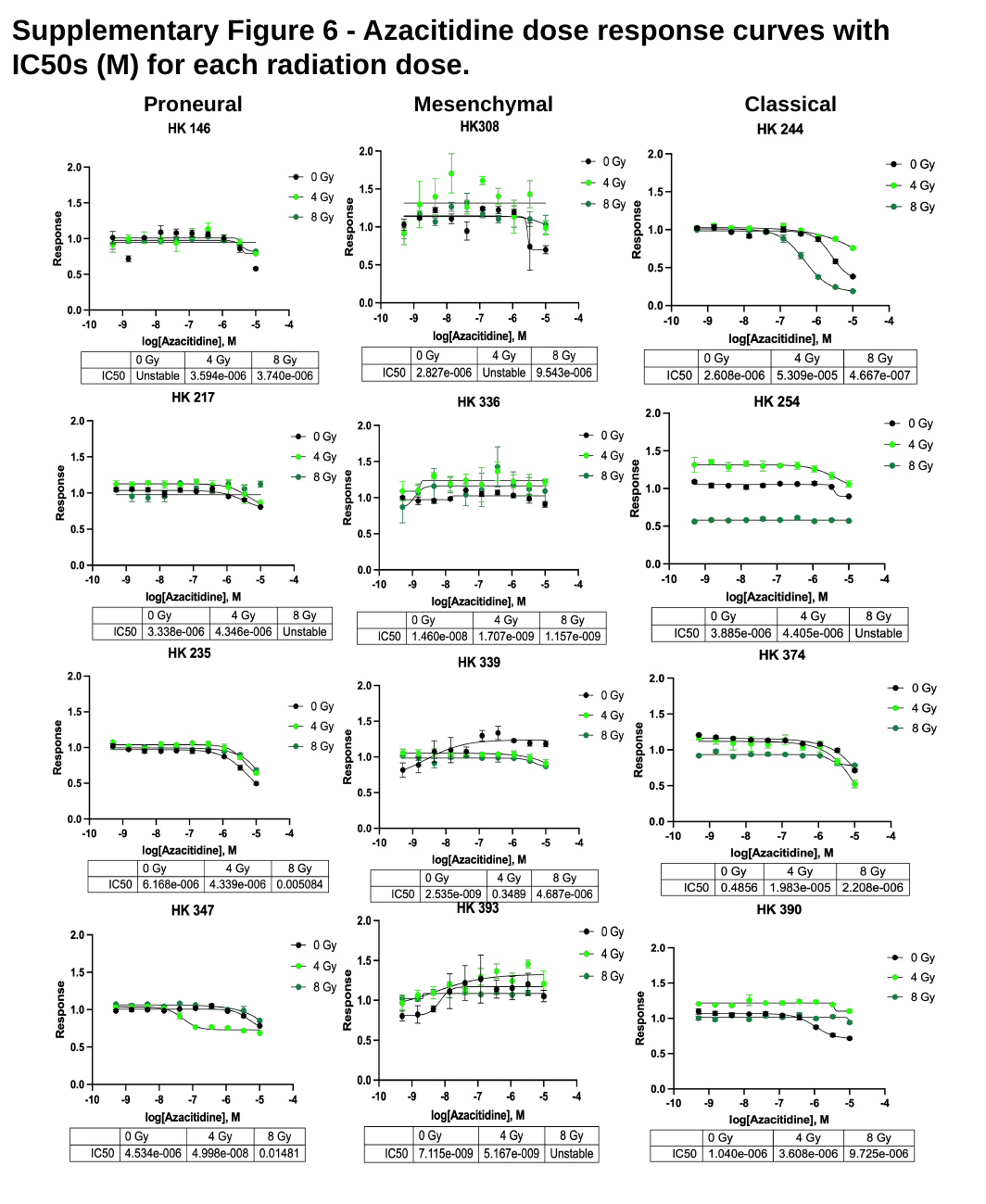

#### Supplementary Figure 6 - Azacitidine dose response curves with IC50s (M) for each radiation dose.
Mesenchymal
Classical
Proneural

### Slide 7
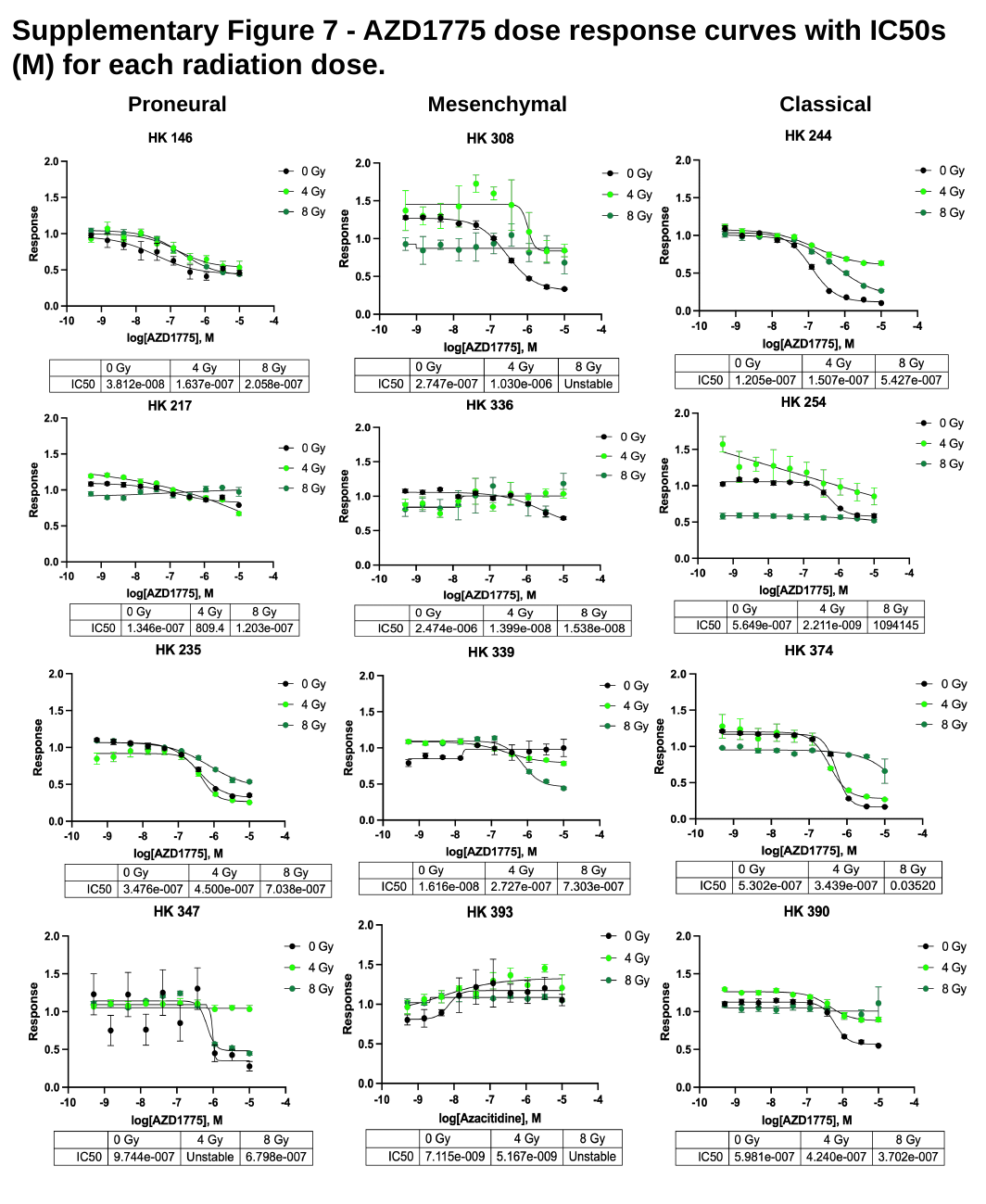

#### Supplementary Figure 7 - AZD1775 dose response curves with IC50s (M) for each radiation dose.
Mesenchymal
Proneural
Classical

### Slide 8
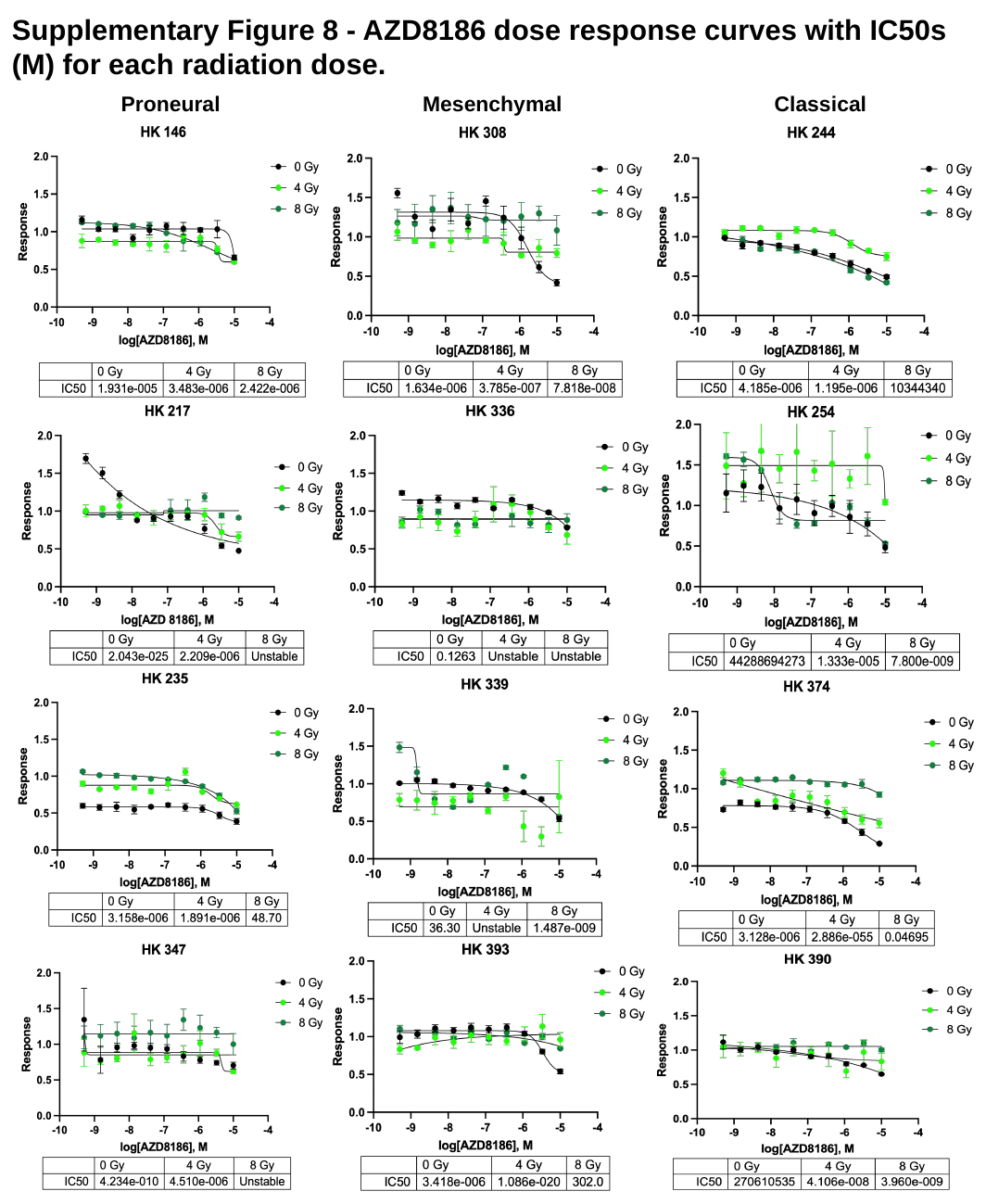

#### Supplementary Figure 8 - AZD8186 dose response curves with IC50s (M) for each radiation dose.
Mesenchymal
Proneural
Classical

### Slide 9
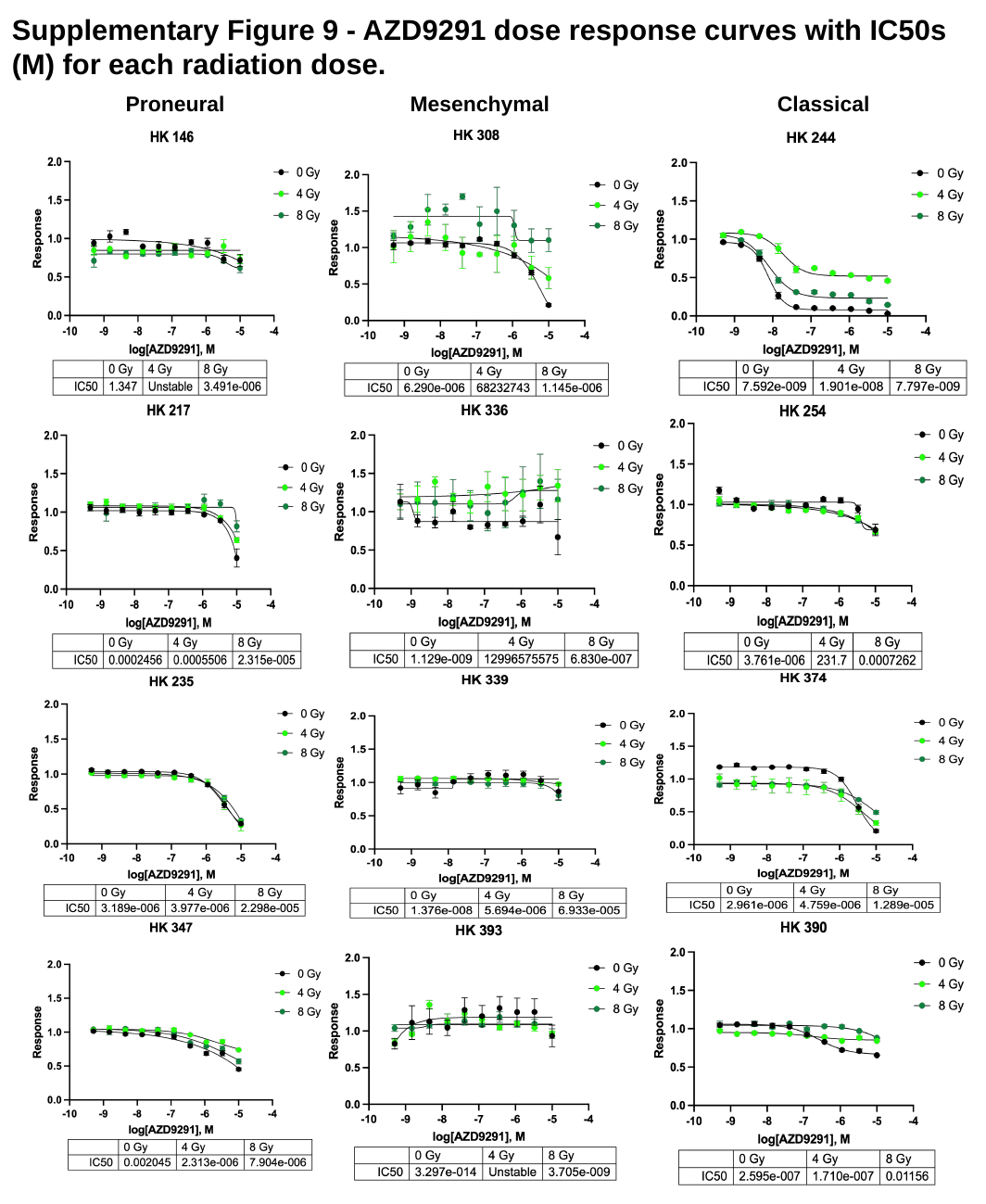

#### Supplementary Figure 9 - AZD9291 dose response curves with IC50s (M) for each radiation dose.
Mesenchymal
Proneural
Classical

### Slide 10
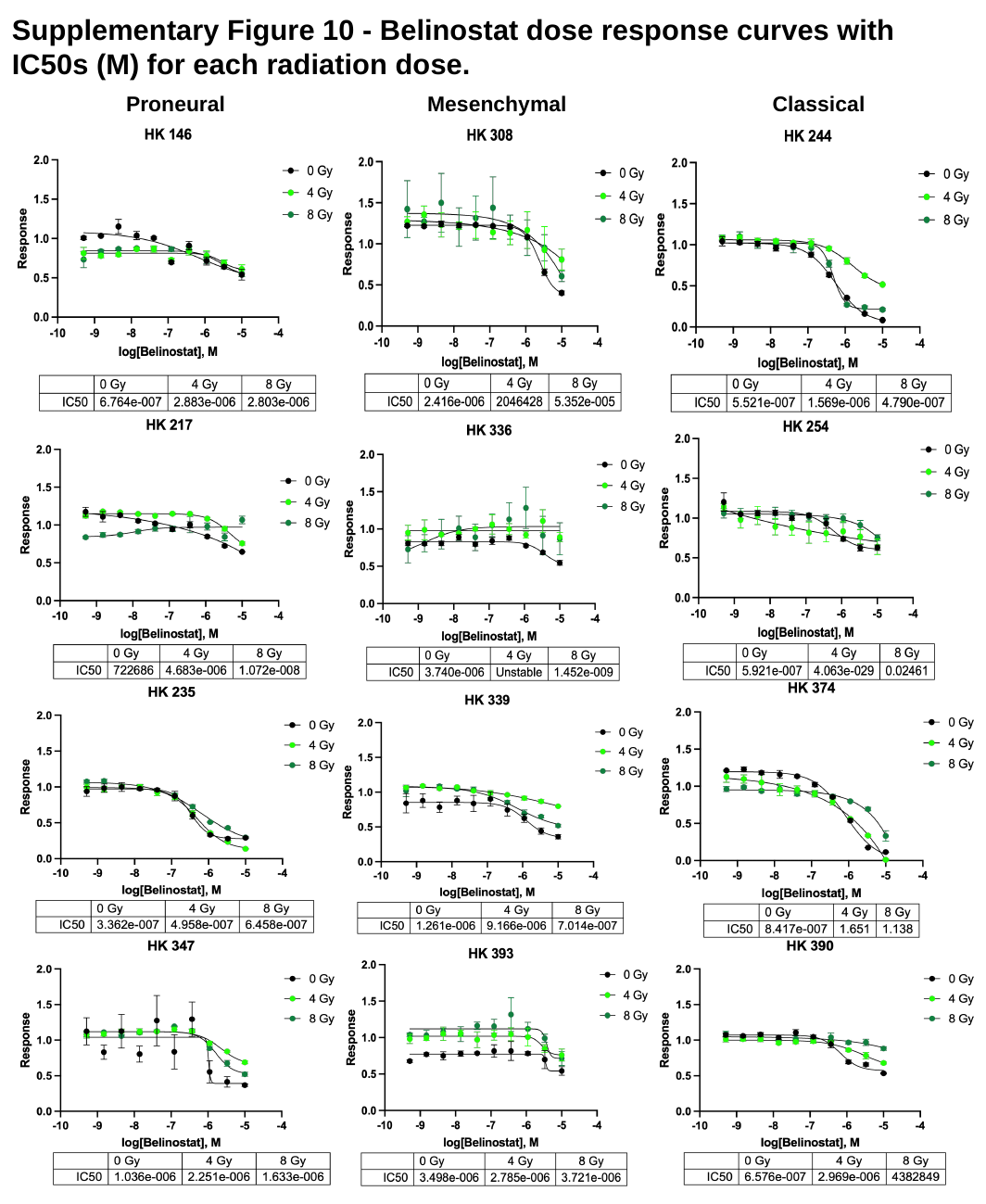

#### Supplementary Figure 10 - Belinostat dose response curves with IC50s (M) for each radiation dose.
Mesenchymal
Classical
Proneural

### Slide 11
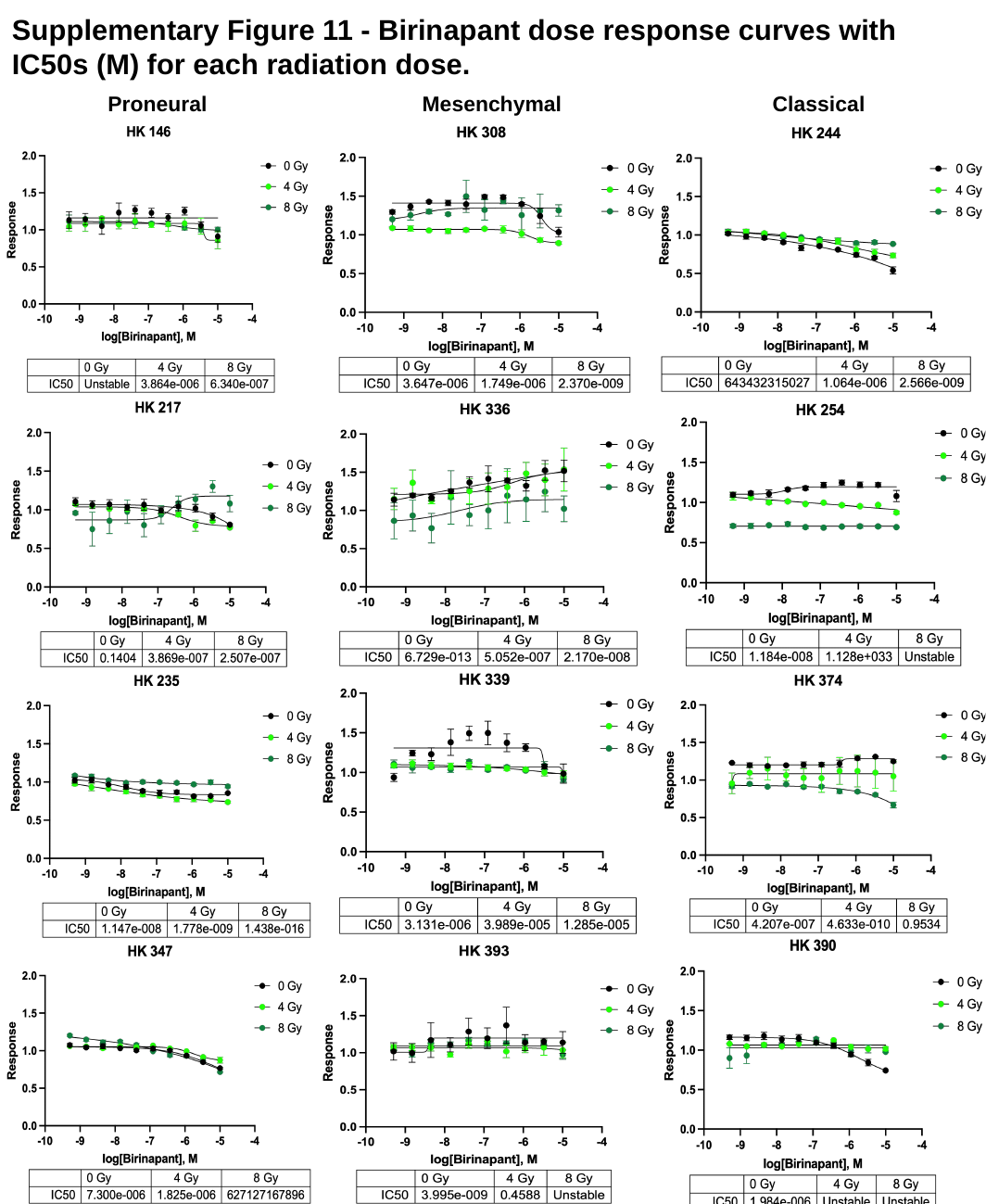

#### Supplementary Figure 11 - Birinapant dose response curves with IC50s (M) for each radiation dose.
Mesenchymal
Proneural
Classical

### Slide 12
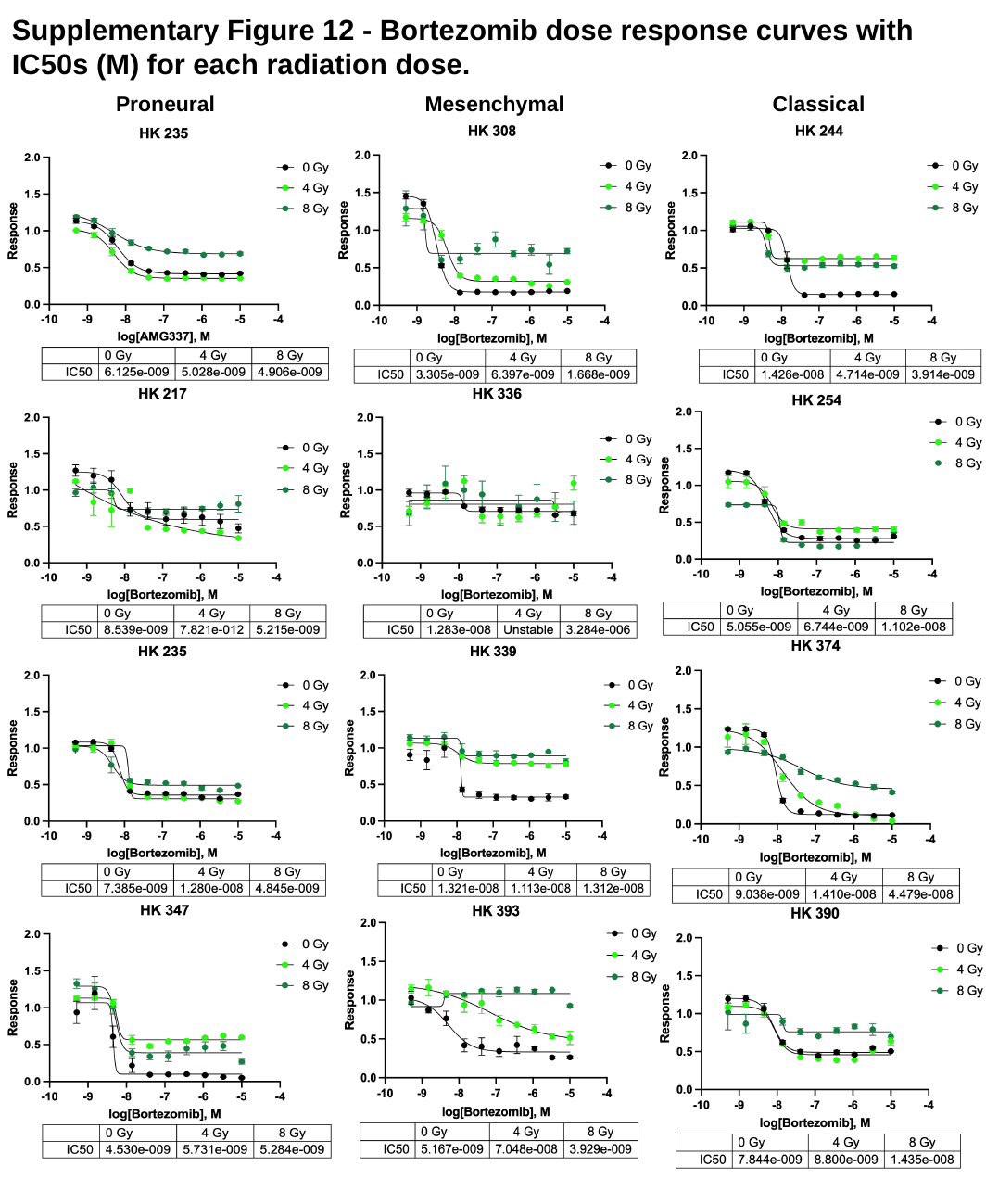

#### Supplementary Figure 12 - Bortezomib dose response curves with IC50s (M) for each radiation dose.
Mesenchymal
Proneural
Classical

### Slide 13
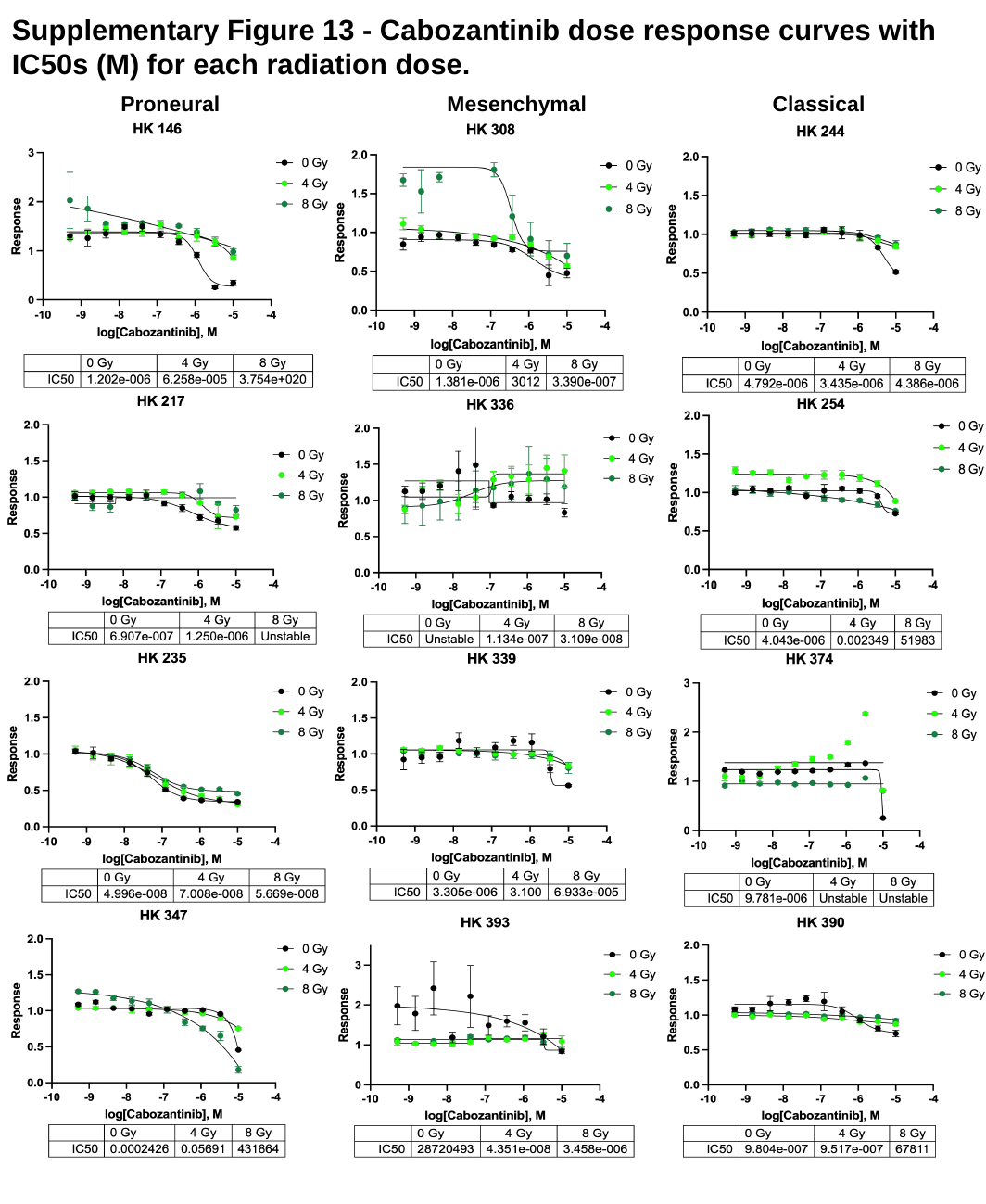

#### Supplementary Figure 13 - Cabozantinib dose response curves with IC50s (M) for each radiation dose.
Proneural
Mesenchymal
Classical

### Slide 14
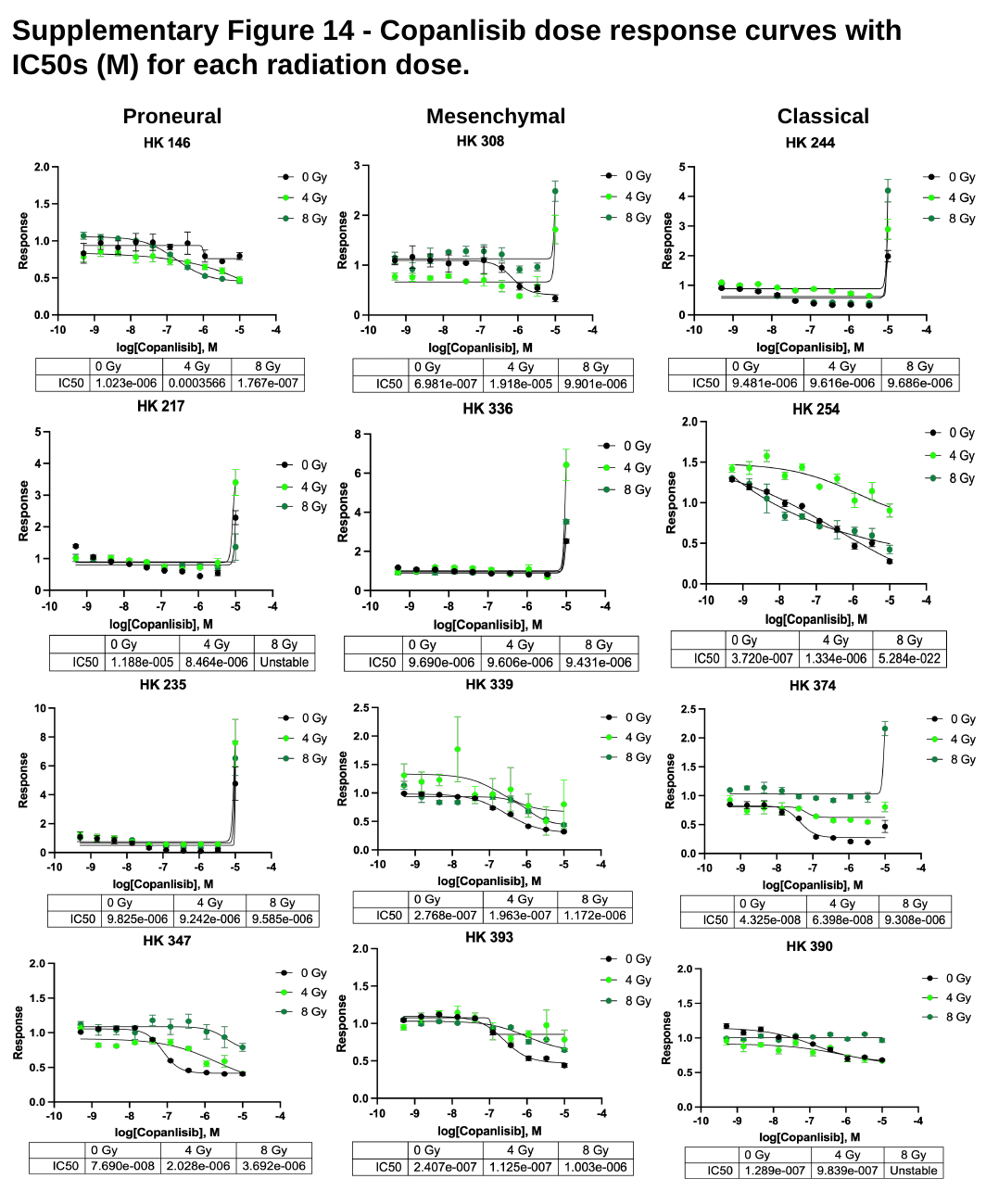

#### Supplementary Figure 14 - Copanlisib dose response curves with IC50s (M) for each radiation dose.
Mesenchymal
Proneural
Classical

### Slide 15
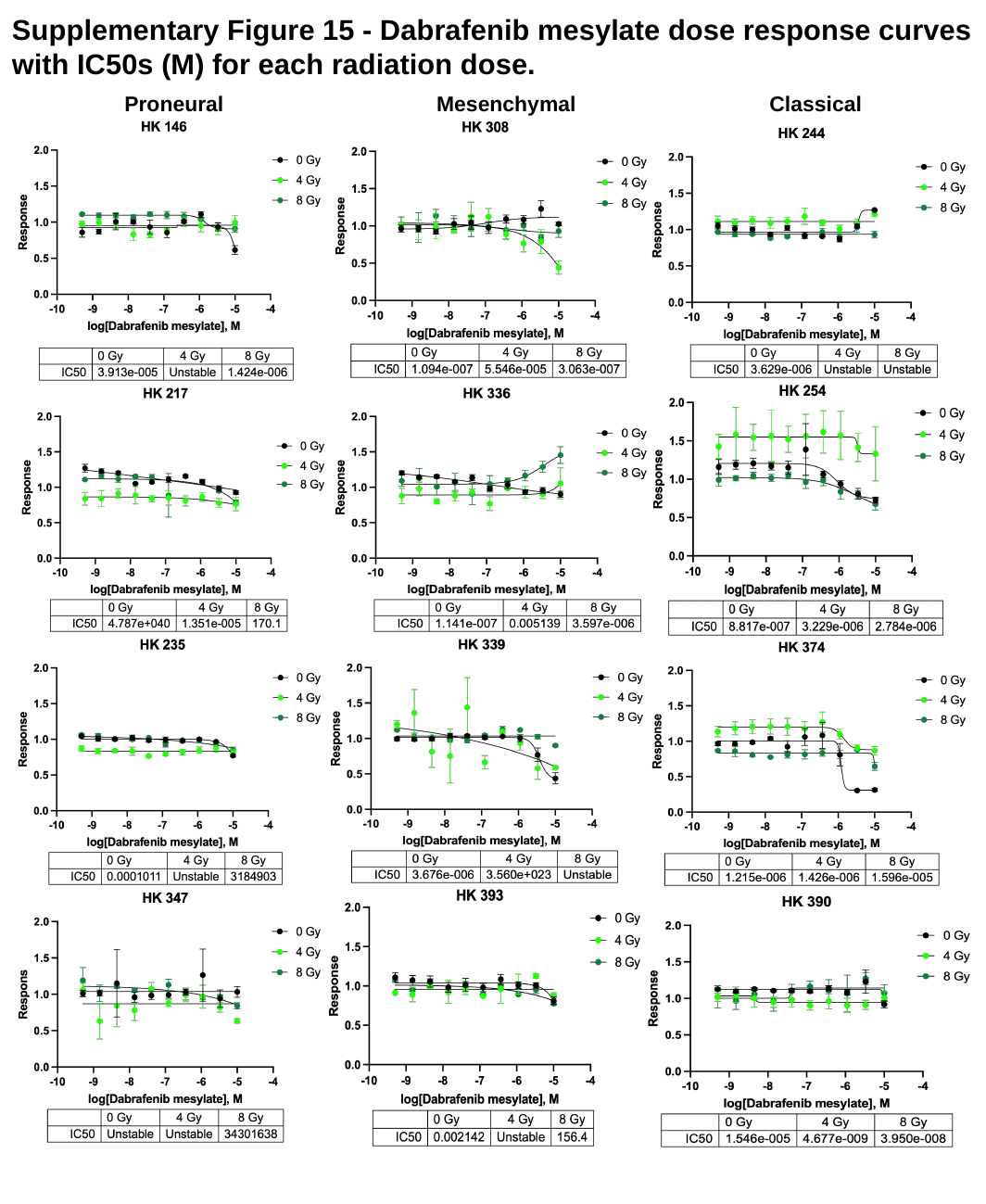

#### Supplementary Figure 15 - Dabrafenib mesylate dose response curves with IC50s (M) for each radiation dose.
Mesenchymal
Proneural
Classical

### Slide 16
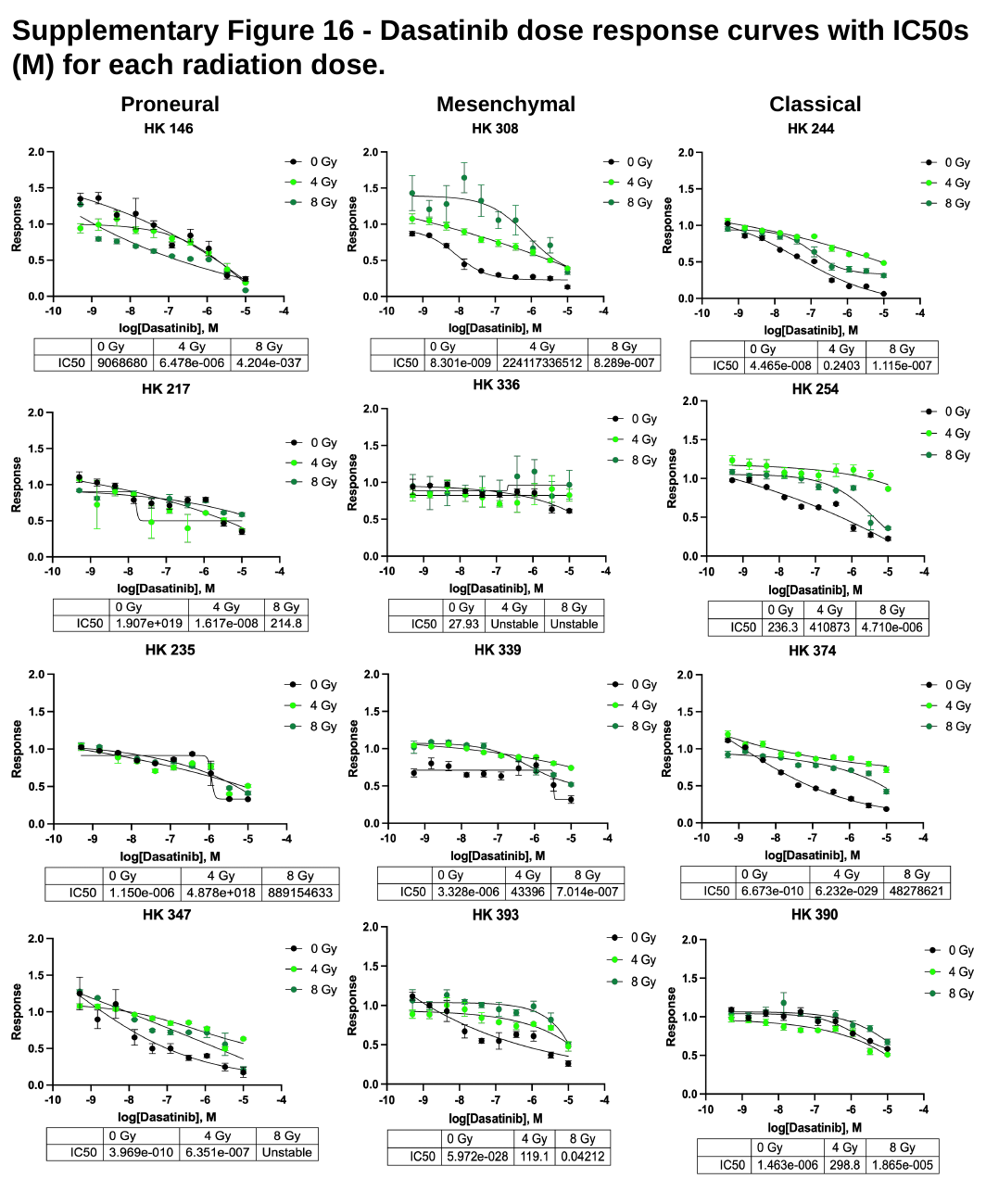

#### Supplementary Figure 16 - Dasatinib dose response curves with IC50s (M) for each radiation dose.
Proneural
Mesenchymal
Classical

### Slide 17
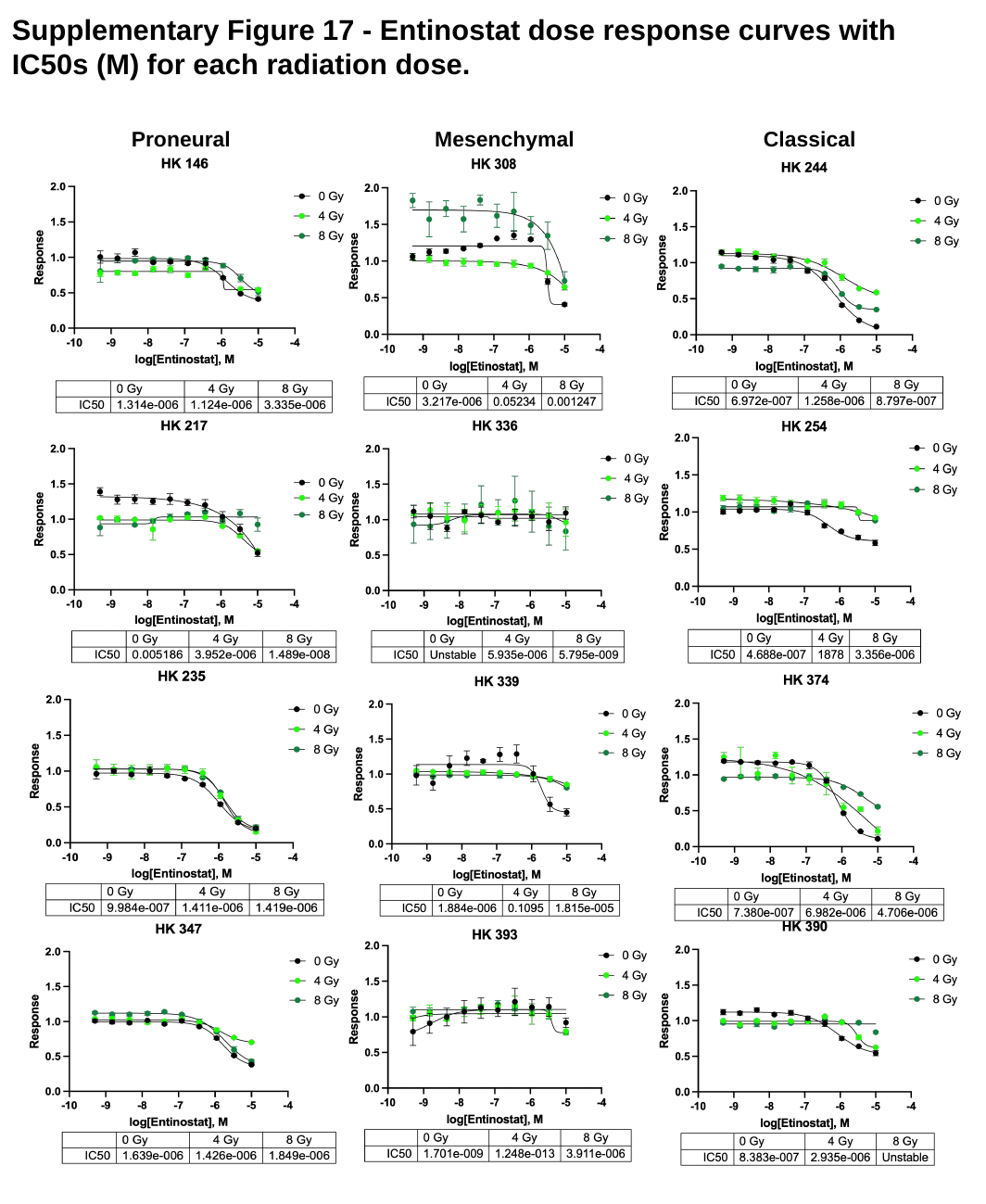

#### Supplementary Figure 17 - Entinostat dose response curves with IC50s (M) for each radiation dose.
Mesenchymal
Proneural
Classical

### Slide 18
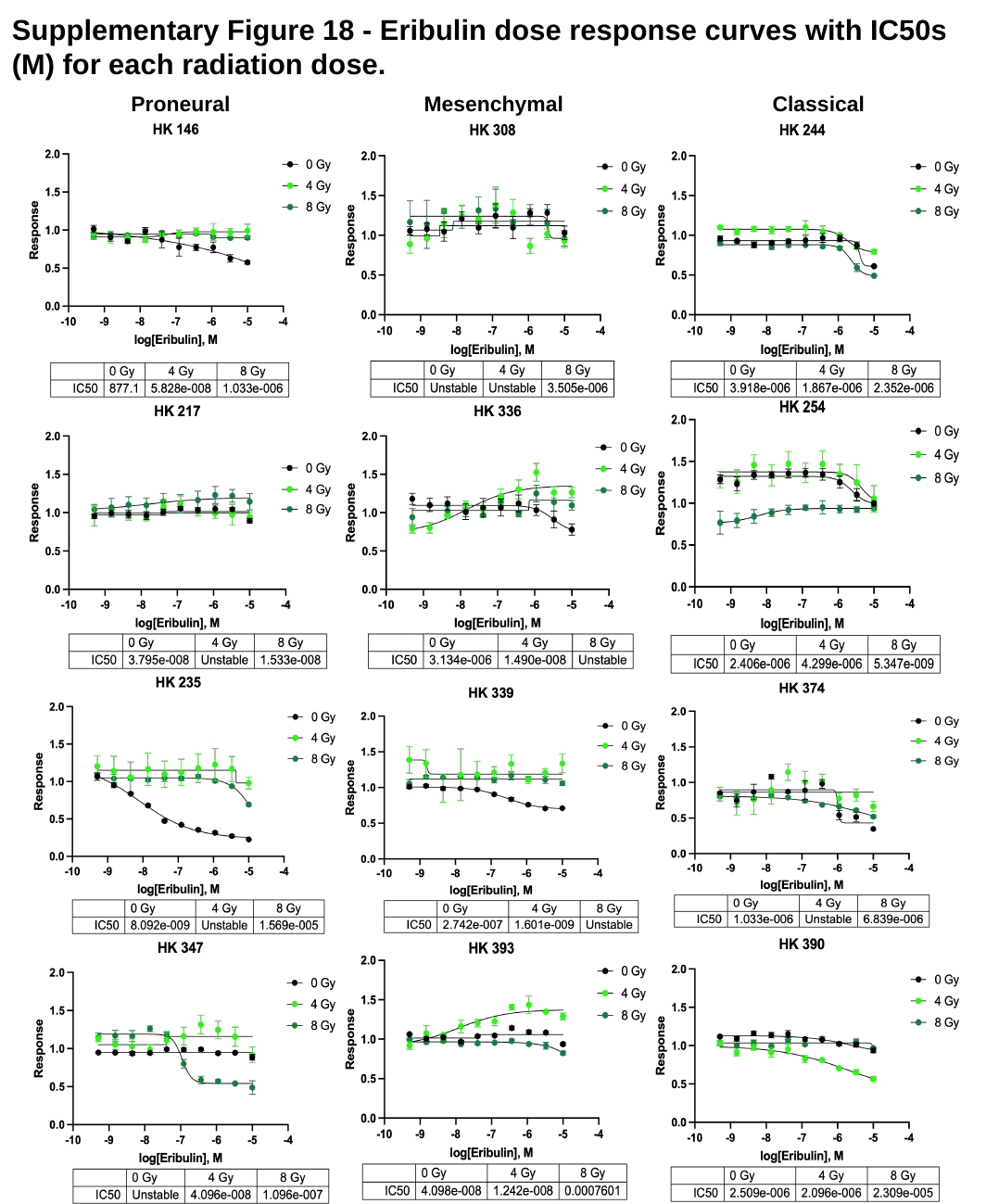

#### Supplementary Figure 18 - Eribulin dose response curves with IC50s (M) for each radiation dose.
Mesenchymal
Proneural
Classical

### Slide 19
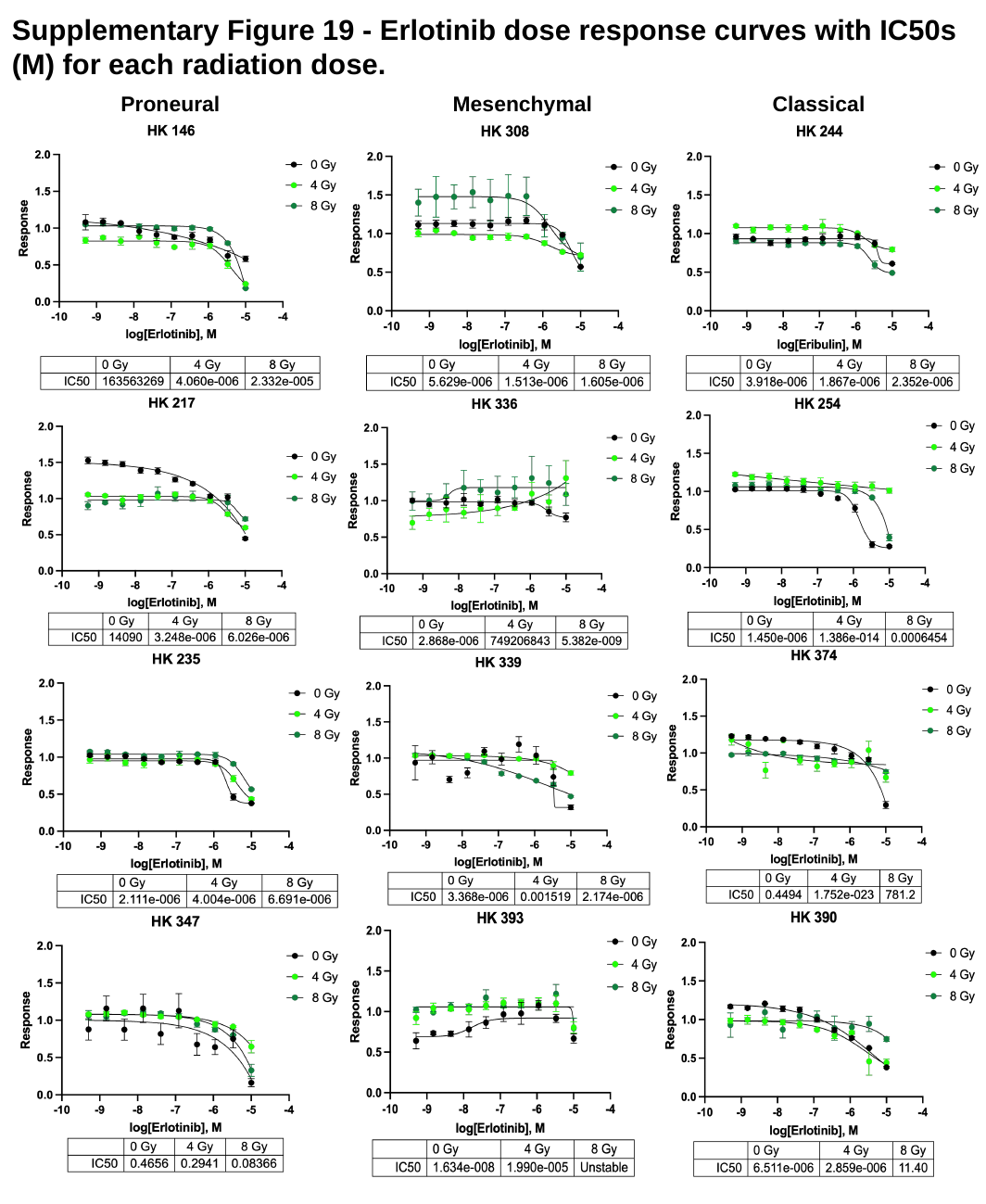

#### Supplementary Figure 19 - Erlotinib dose response curves with IC50s (M) for each radiation dose.
Proneural
Mesenchymal
Classical

### Slide 20
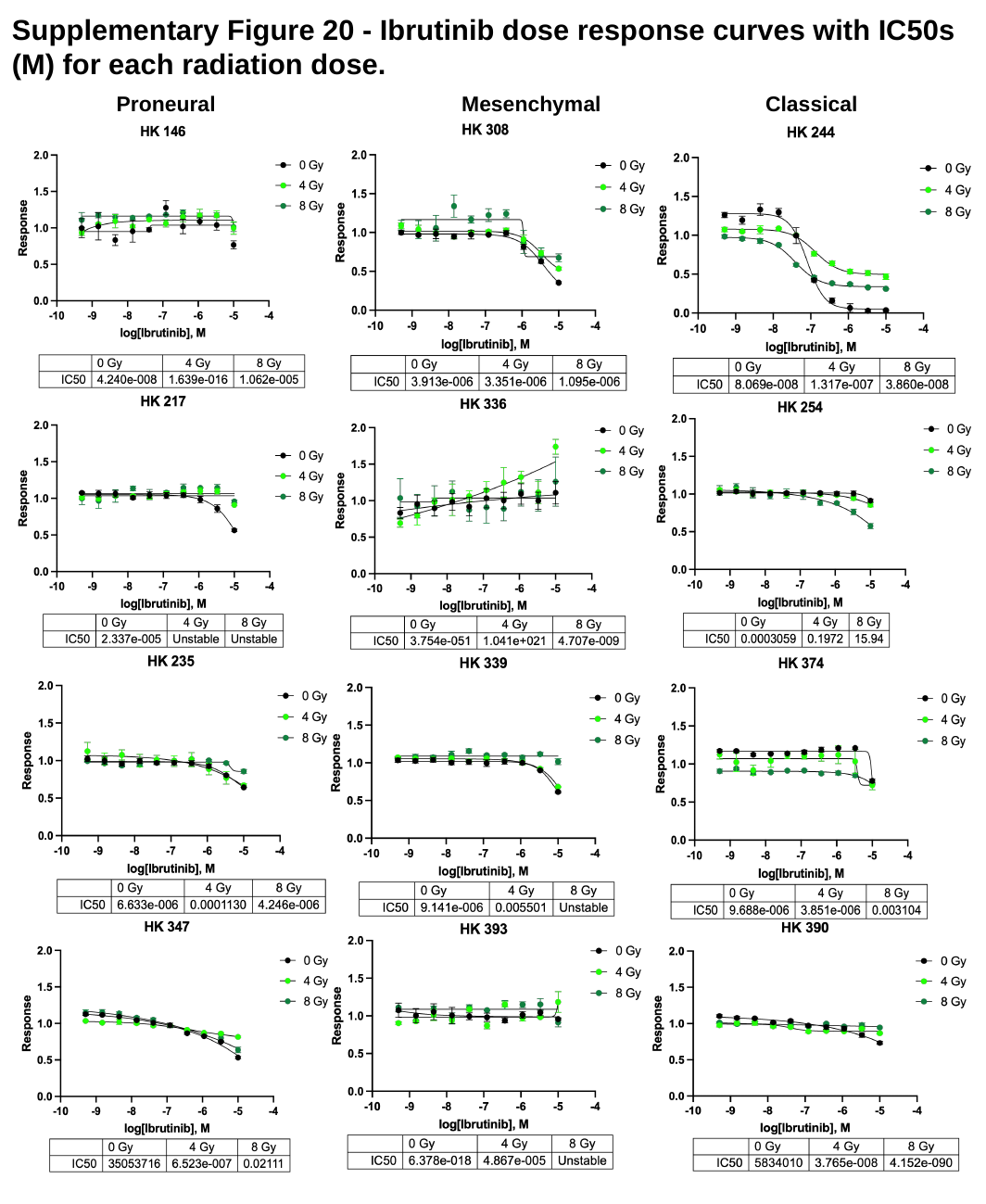

#### Supplementary Figure 20 - Ibrutinib dose response curves with IC50s (M) for each radiation dose.
Proneural
Mesenchymal
Classical

### Slide 21
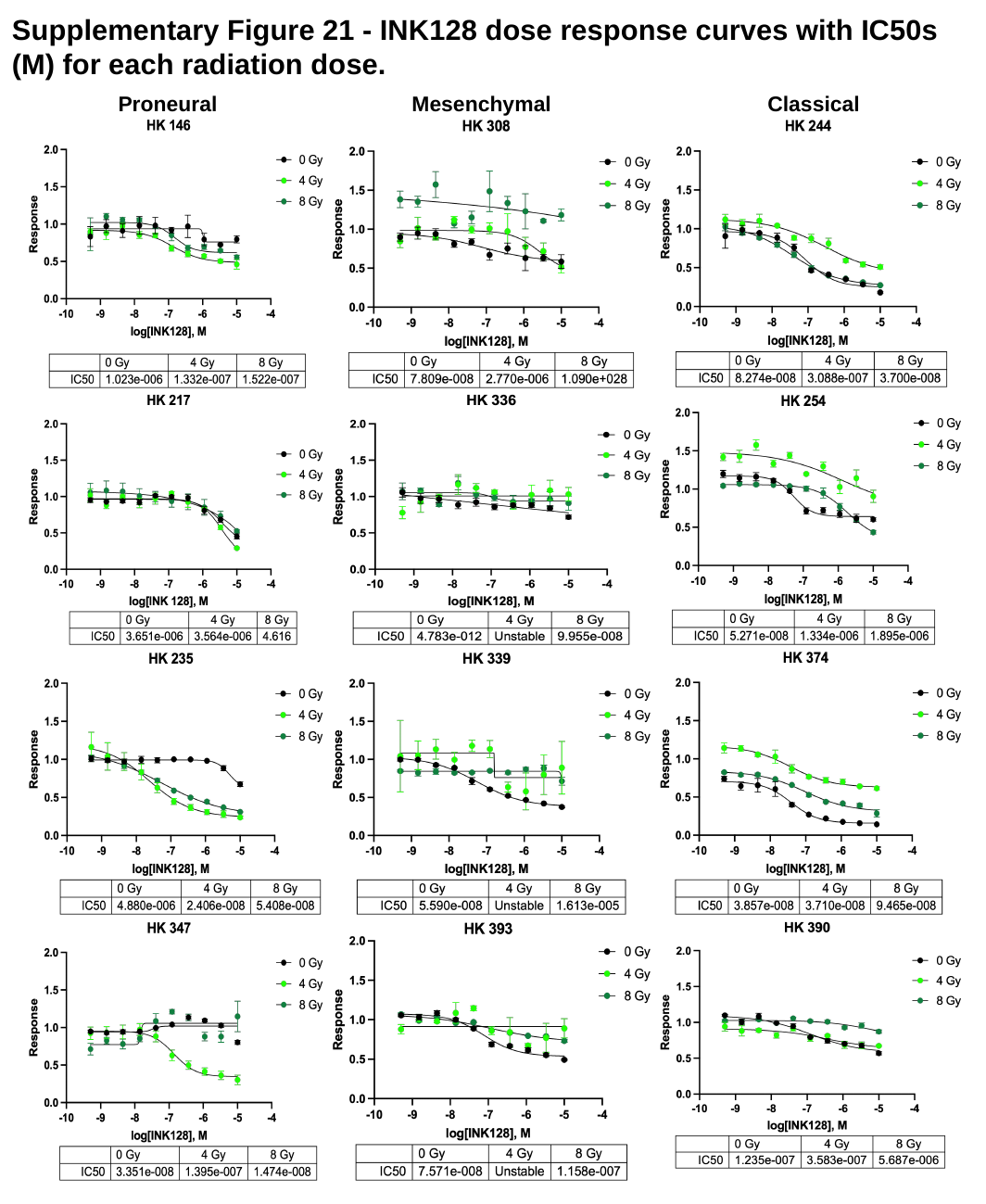

#### Supplementary Figure 21 - INK128 dose response curves with IC50s (M) for each radiation dose.
Mesenchymal
Proneural
Classical

### Slide 22
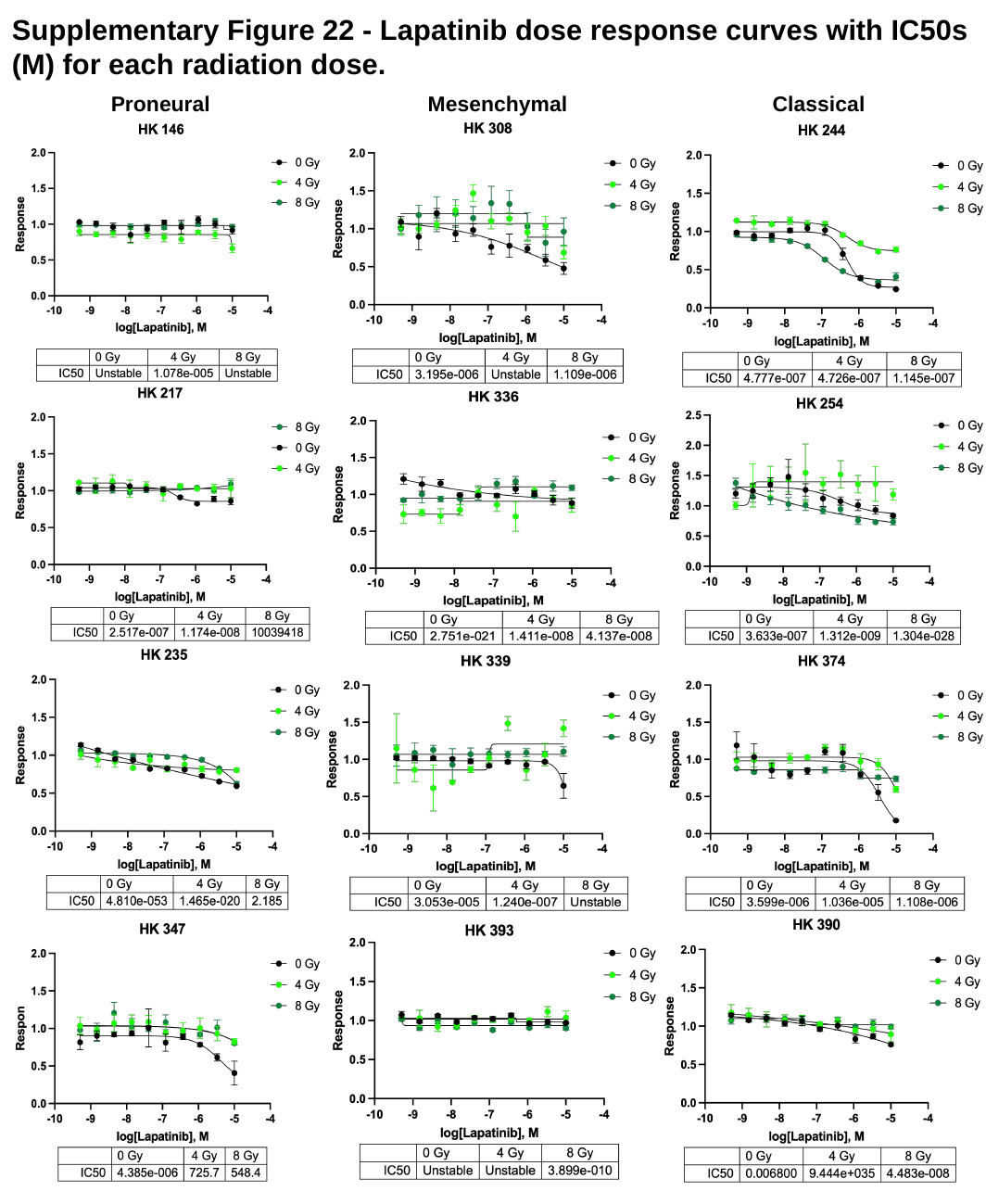

#### Supplementary Figure 22 - Lapatinib dose response curves with IC50s (M) for each radiation dose.
Mesenchymal
Proneural
Classical

### Slide 23
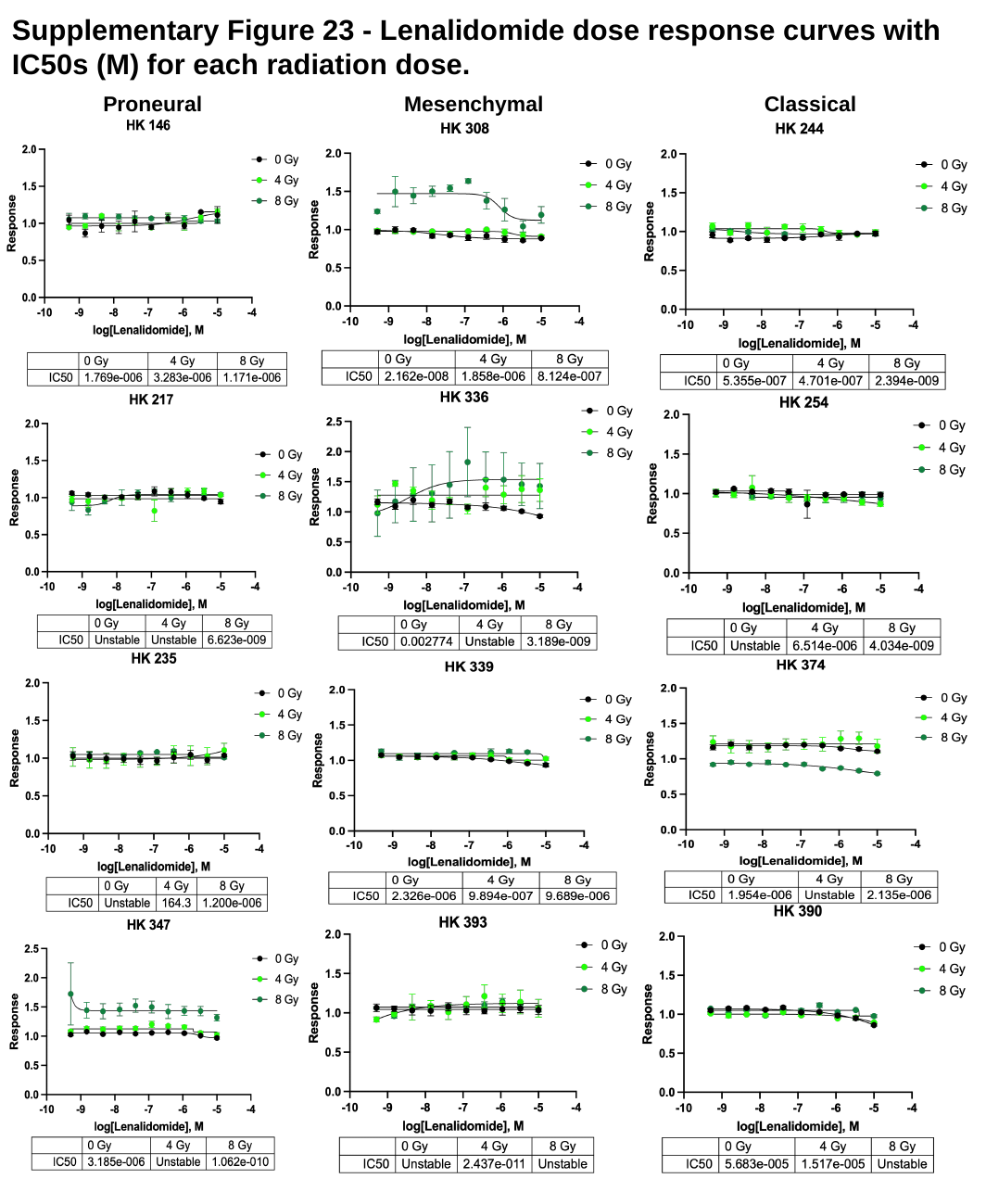

#### Supplementary Figure 23 - Lenalidomide dose response curves with IC50s (M) for each radiation dose.
Mesenchymal
Proneural
Classical

### Slide 24
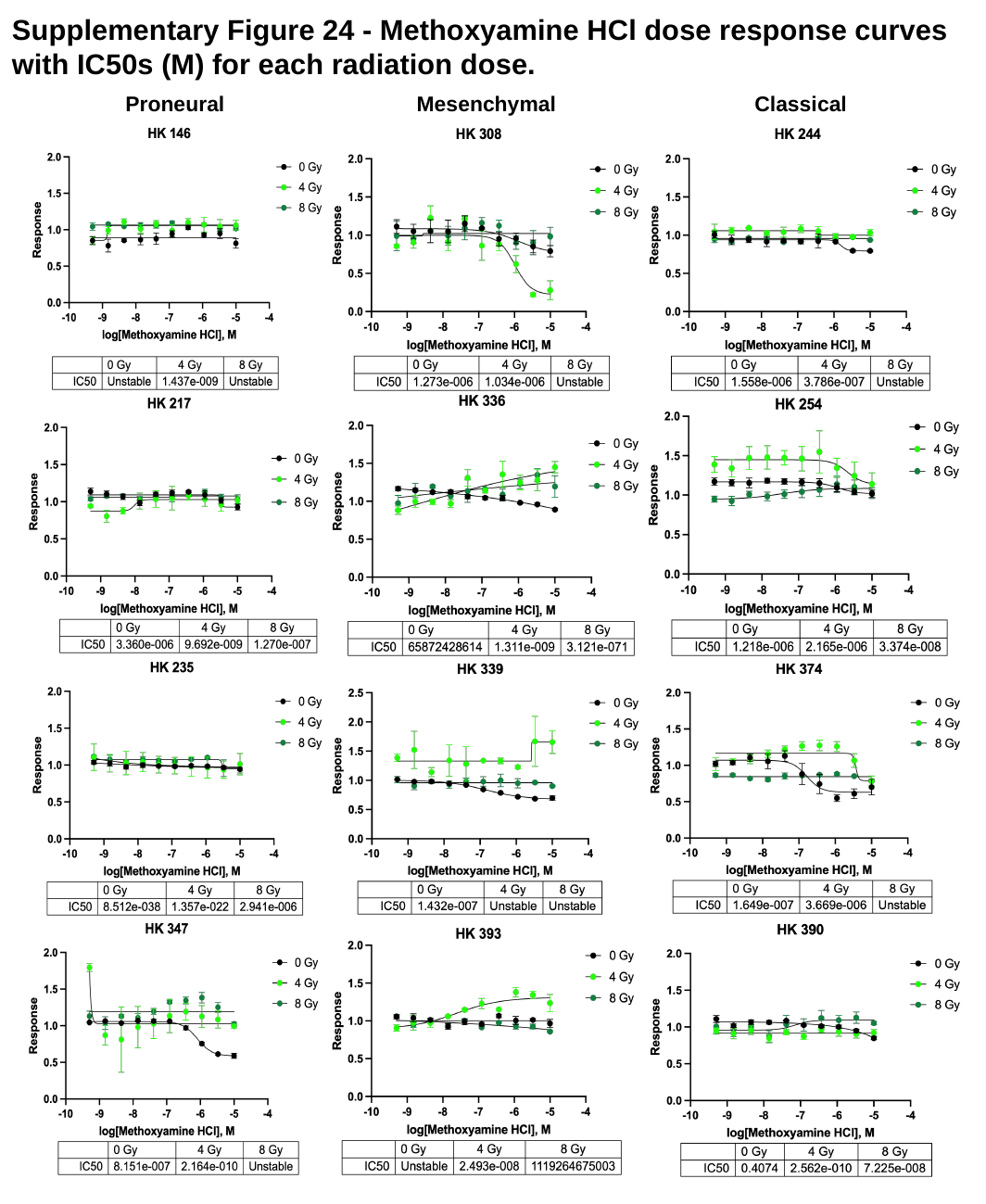

#### Supplementary Figure 24 - Methoxyamine HCl dose response curves with IC50s (M) for each radiation dose.
Mesenchymal
Classical
Proneural

### Slide 25
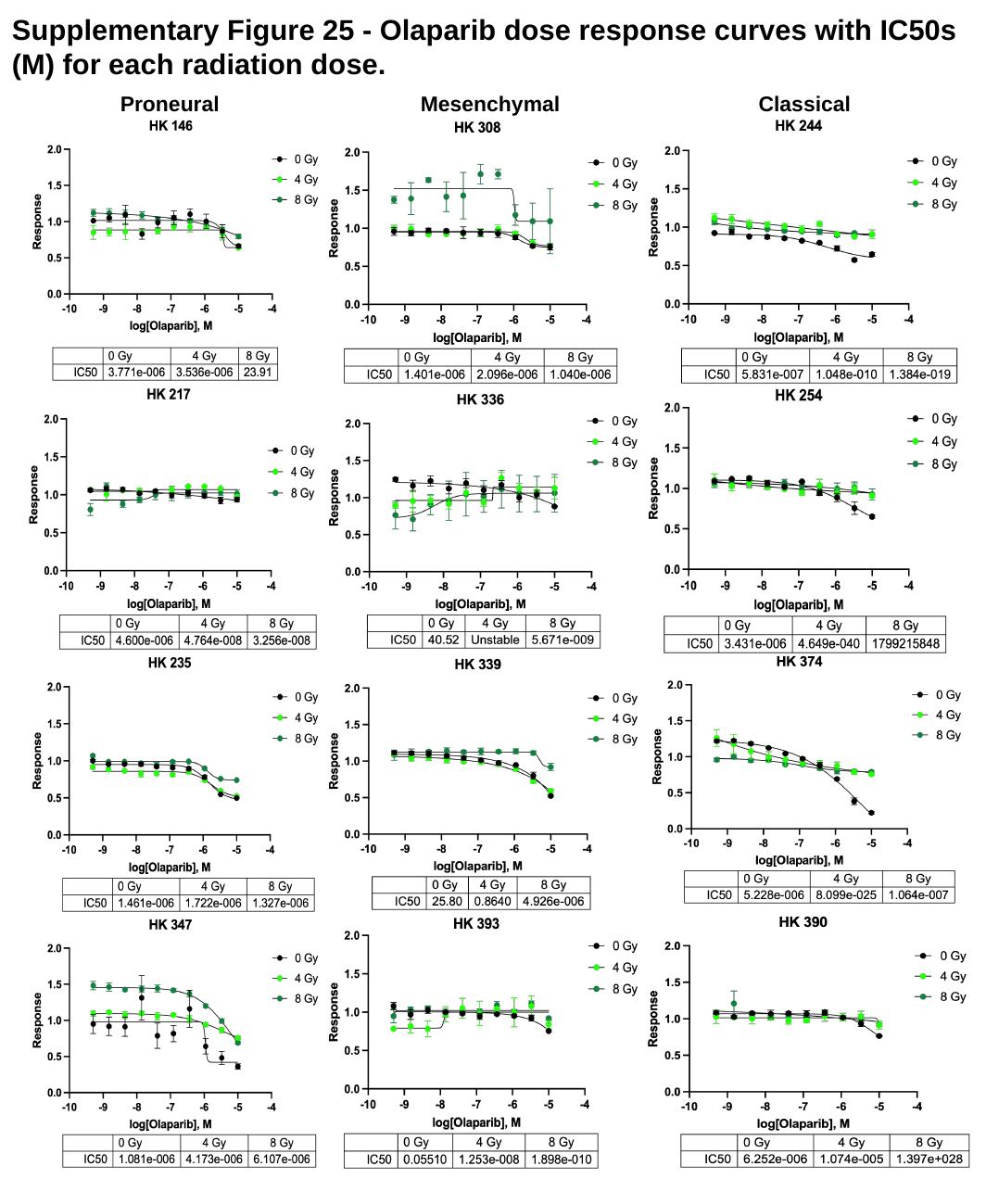

#### Supplementary Figure 25 - Olaparib dose response curves with IC50s (M) for each radiation dose.
Mesenchymal
Classical
Proneural

### Slide 26
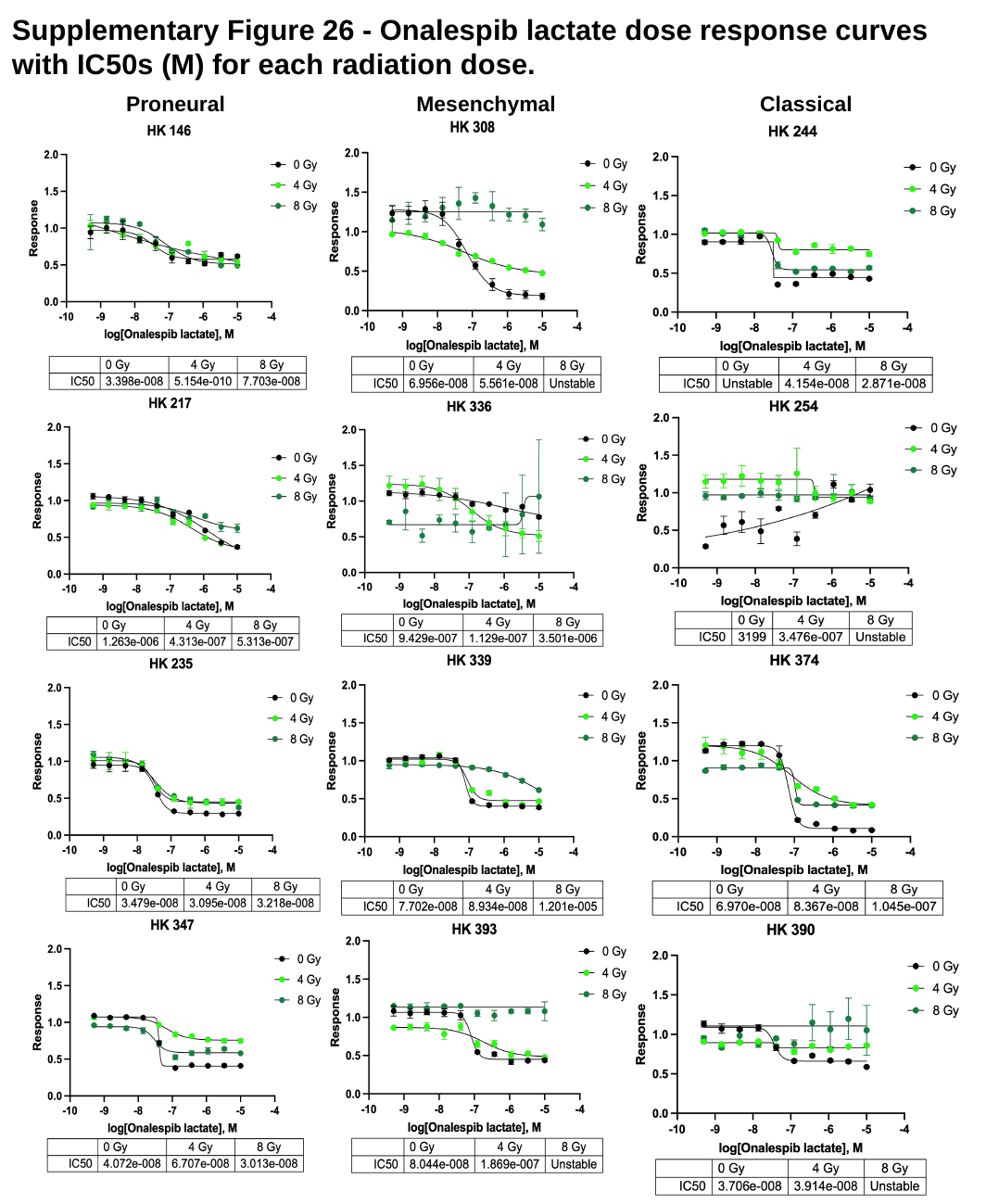

#### Supplementary Figure 26 - Onalespib lactate dose response curves with IC50s (M) for each radiation dose.
Mesenchymal
Classical
Proneural

### Slide 27
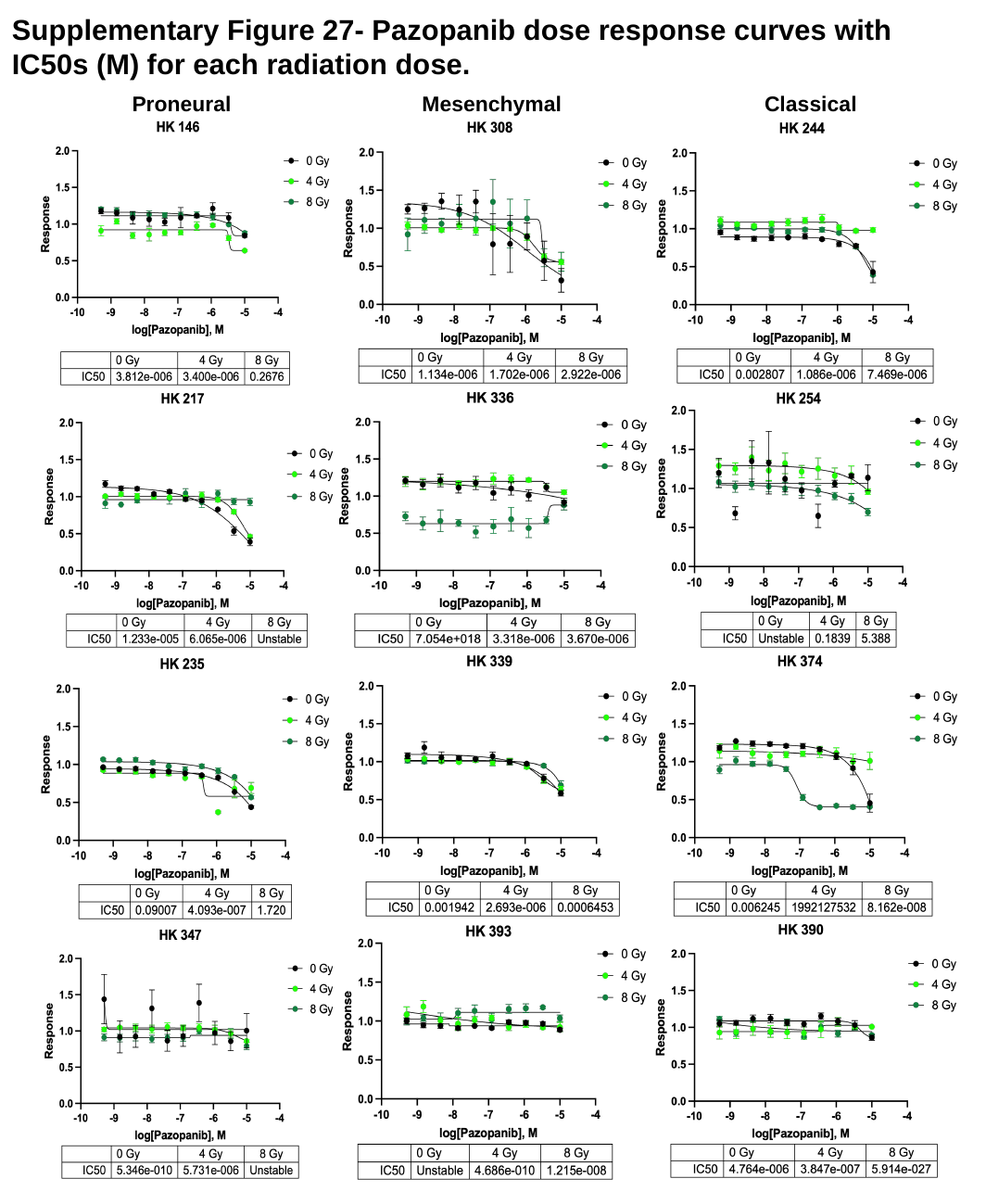

#### Supplementary Figure 27- Pazopanib dose response curves with IC50s (M) for each radiation dose.
Mesenchymal
Classical
Proneural

### Slide 28
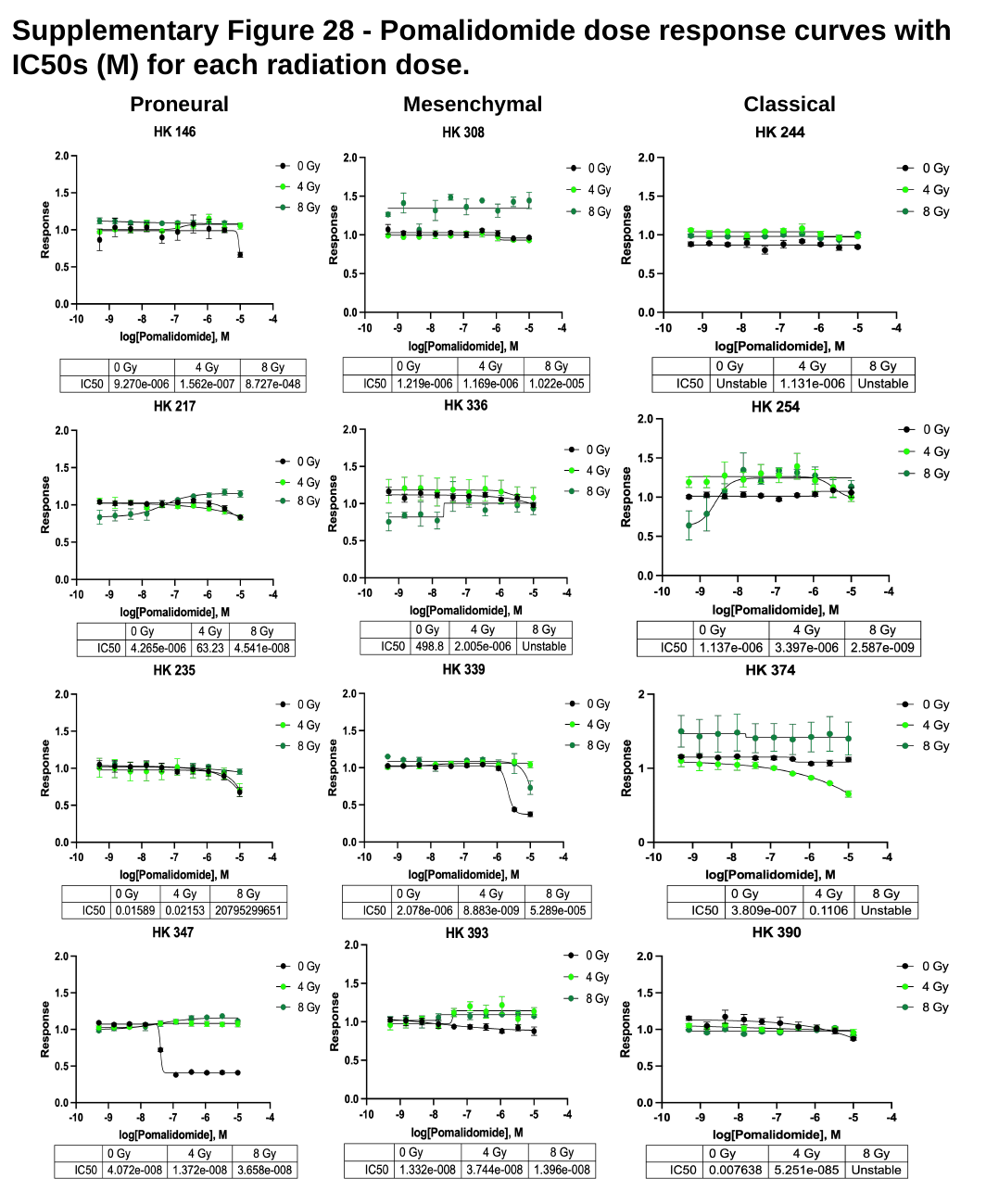

#### Supplementary Figure 28 - Pomalidomide dose response curves with IC50s (M) for each radiation dose.
Mesenchymal
Classical
Proneural

### Slide 29
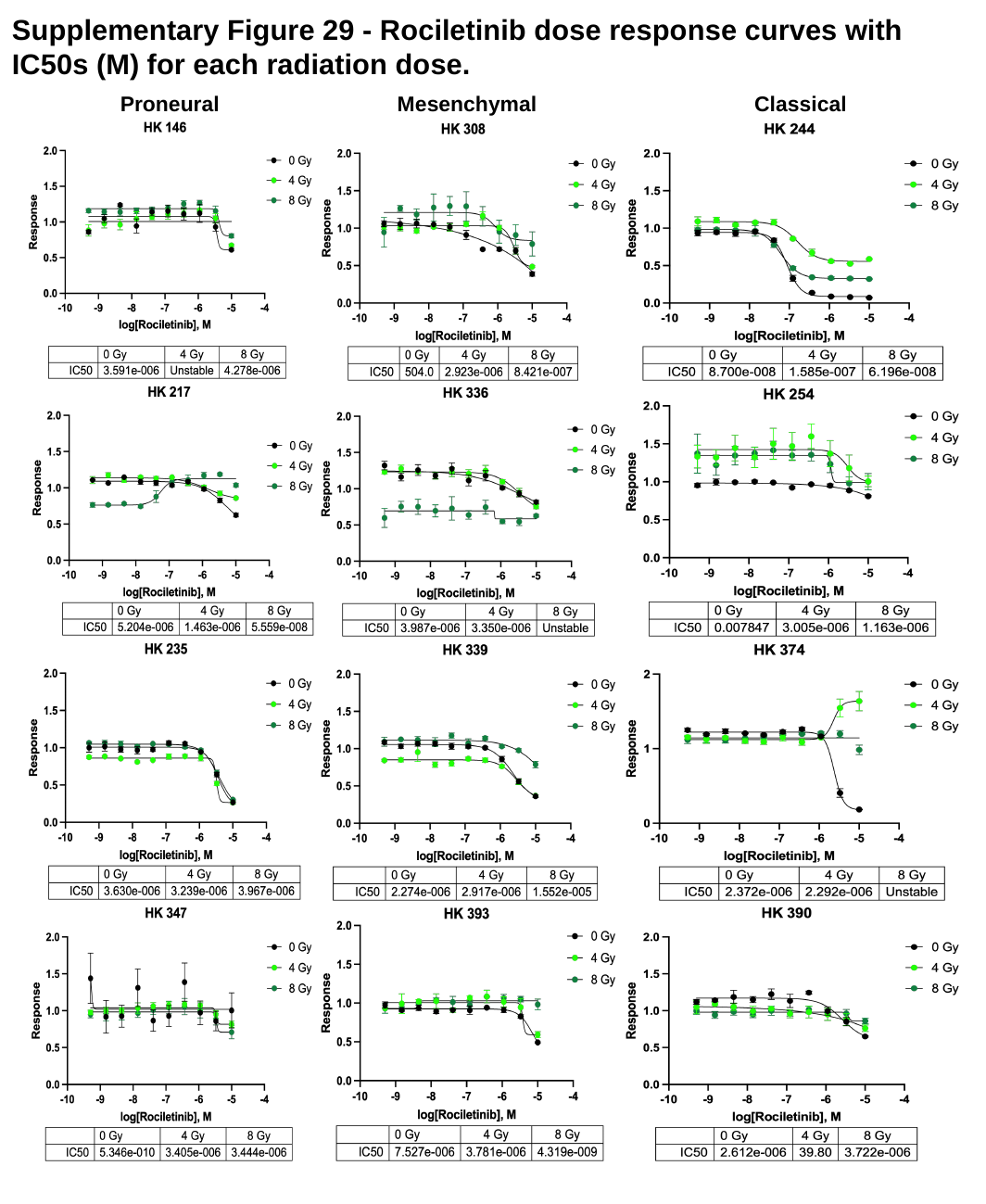

#### Supplementary Figure 29 - Rociletinib dose response curves with IC50s (M) for each radiation dose.
Classical
Proneural
Mesenchymal

### Slide 30
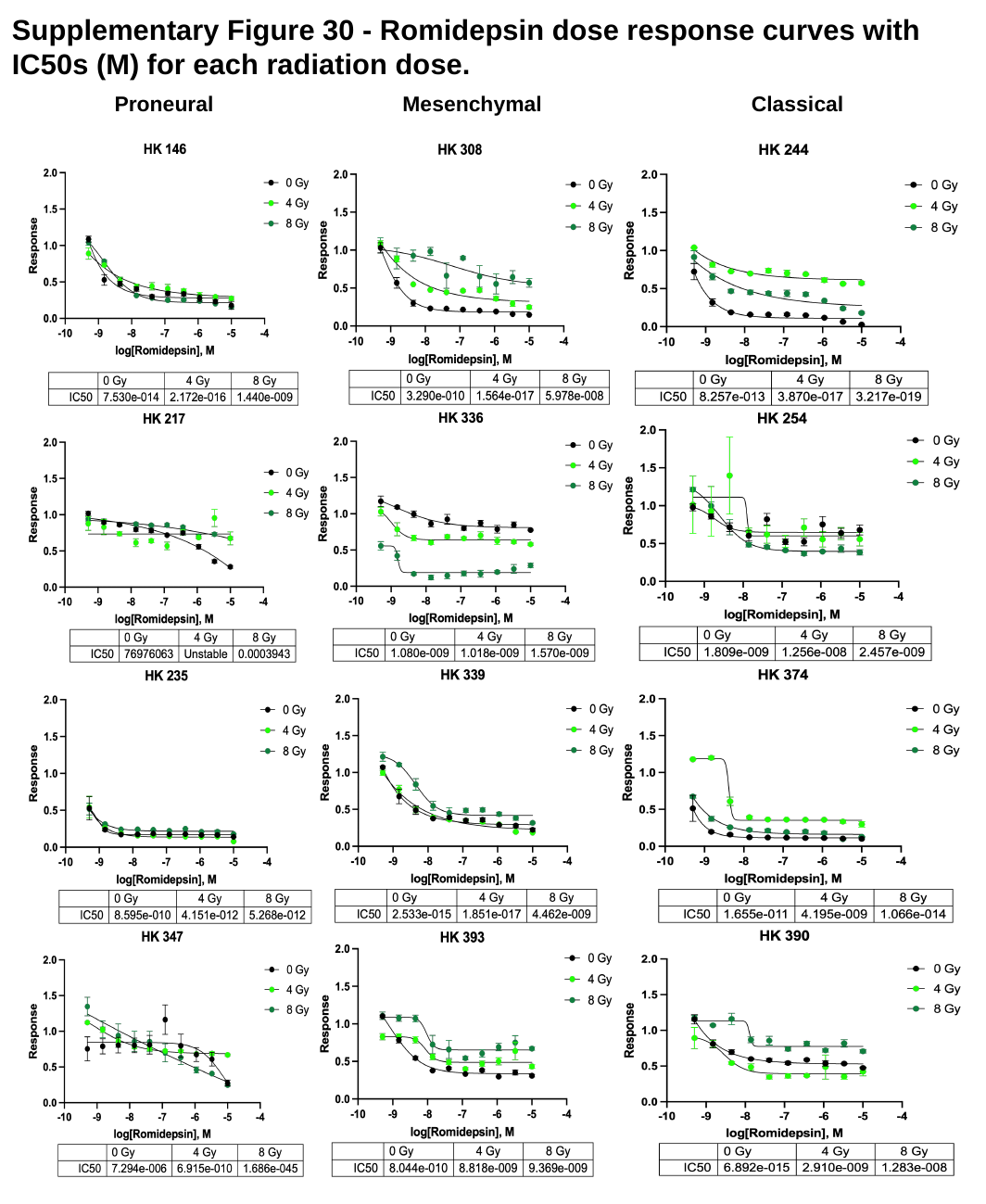

#### Supplementary Figure 30 - Romidepsin dose response curves with IC50s (M) for each radiation dose.
Mesenchymal
Classical
Proneural

### Slide 31
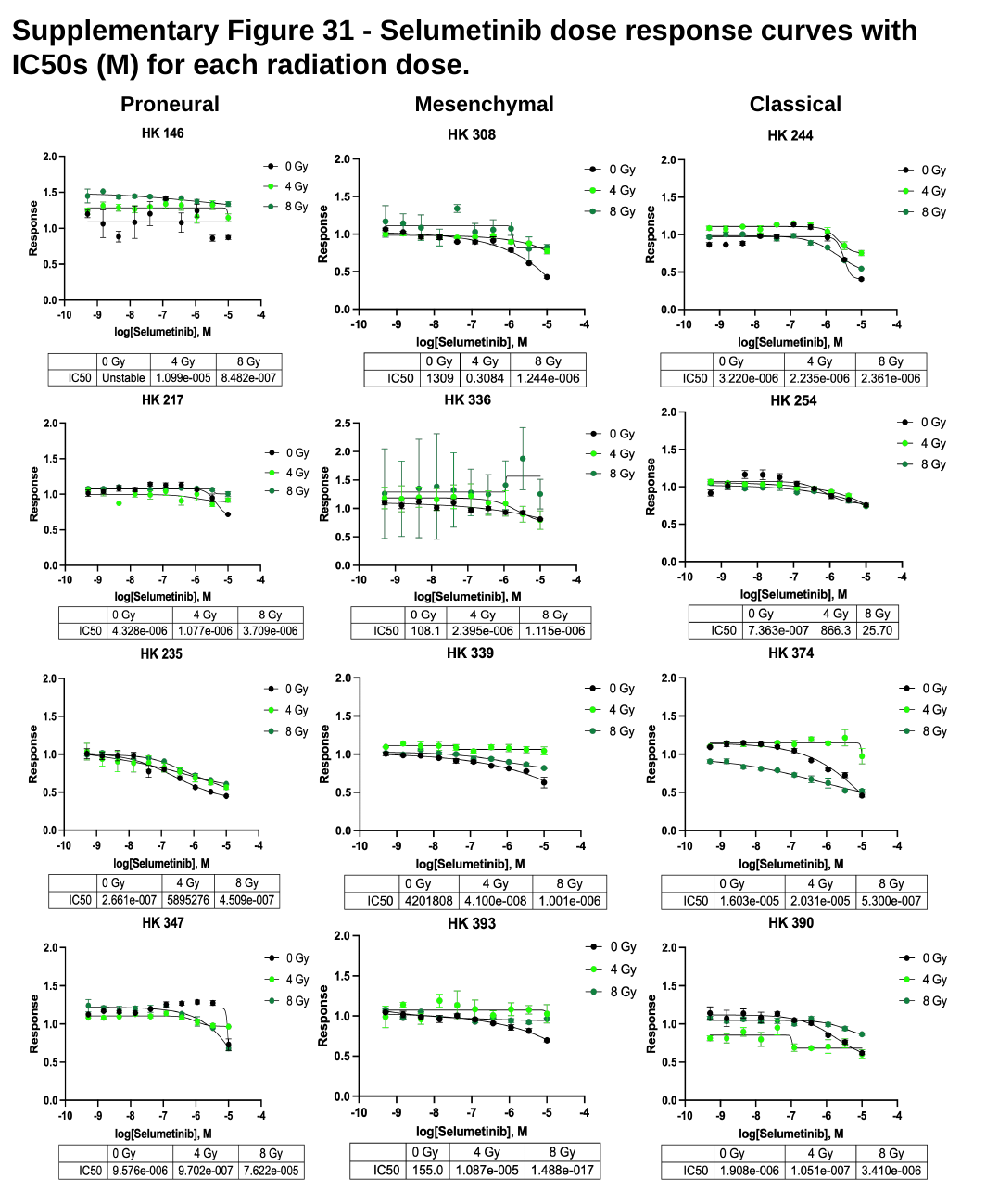

#### Supplementary Figure 31 - Selumetinib dose response curves with IC50s (M) for each radiation dose.
Mesenchymal
Classical
Proneural

### Slide 32
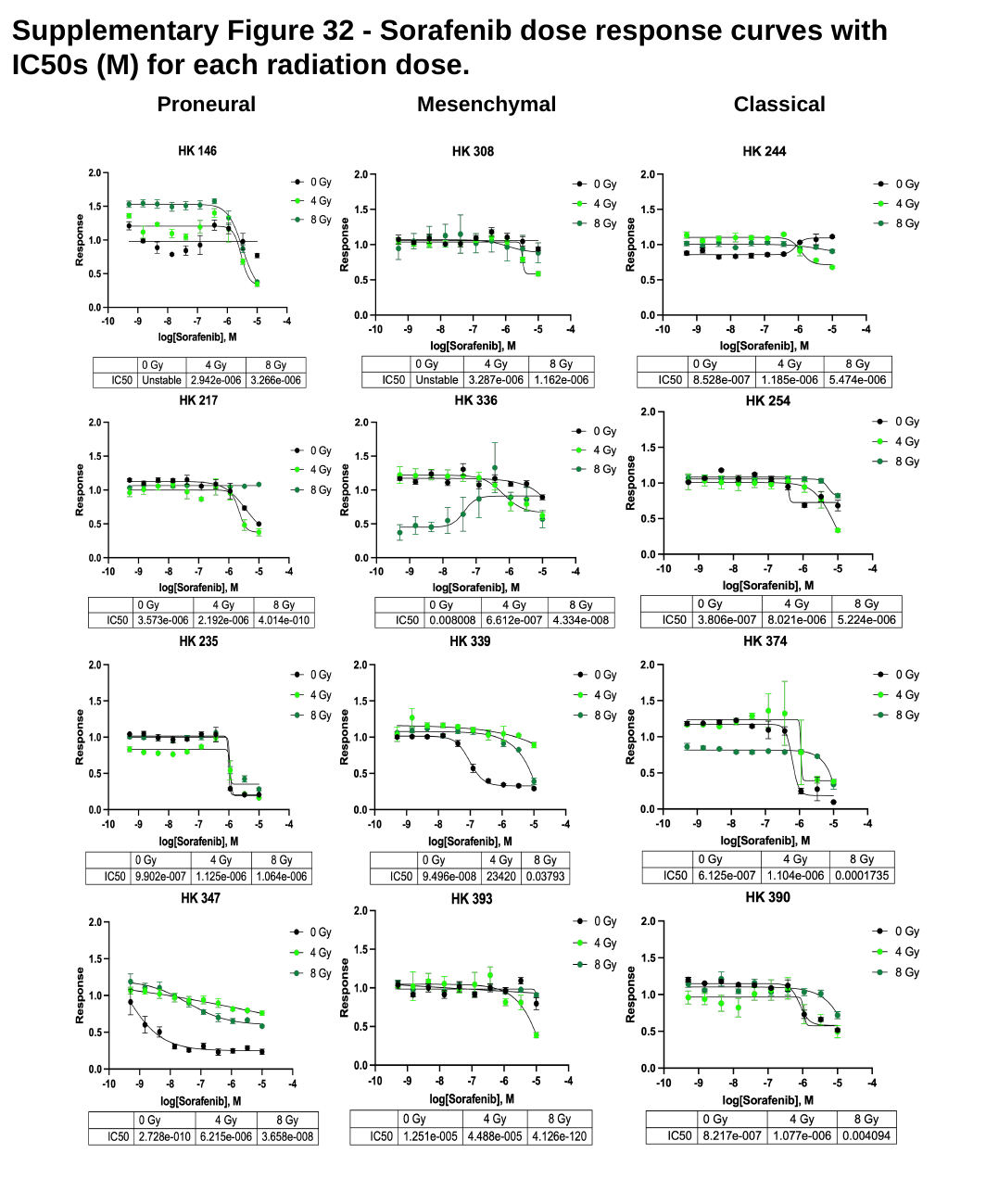

#### Supplementary Figure 32 - Sorafenib dose response curves with IC50s (M) for each radiation dose.
Mesenchymal
Classical
Proneural

### Slide 33
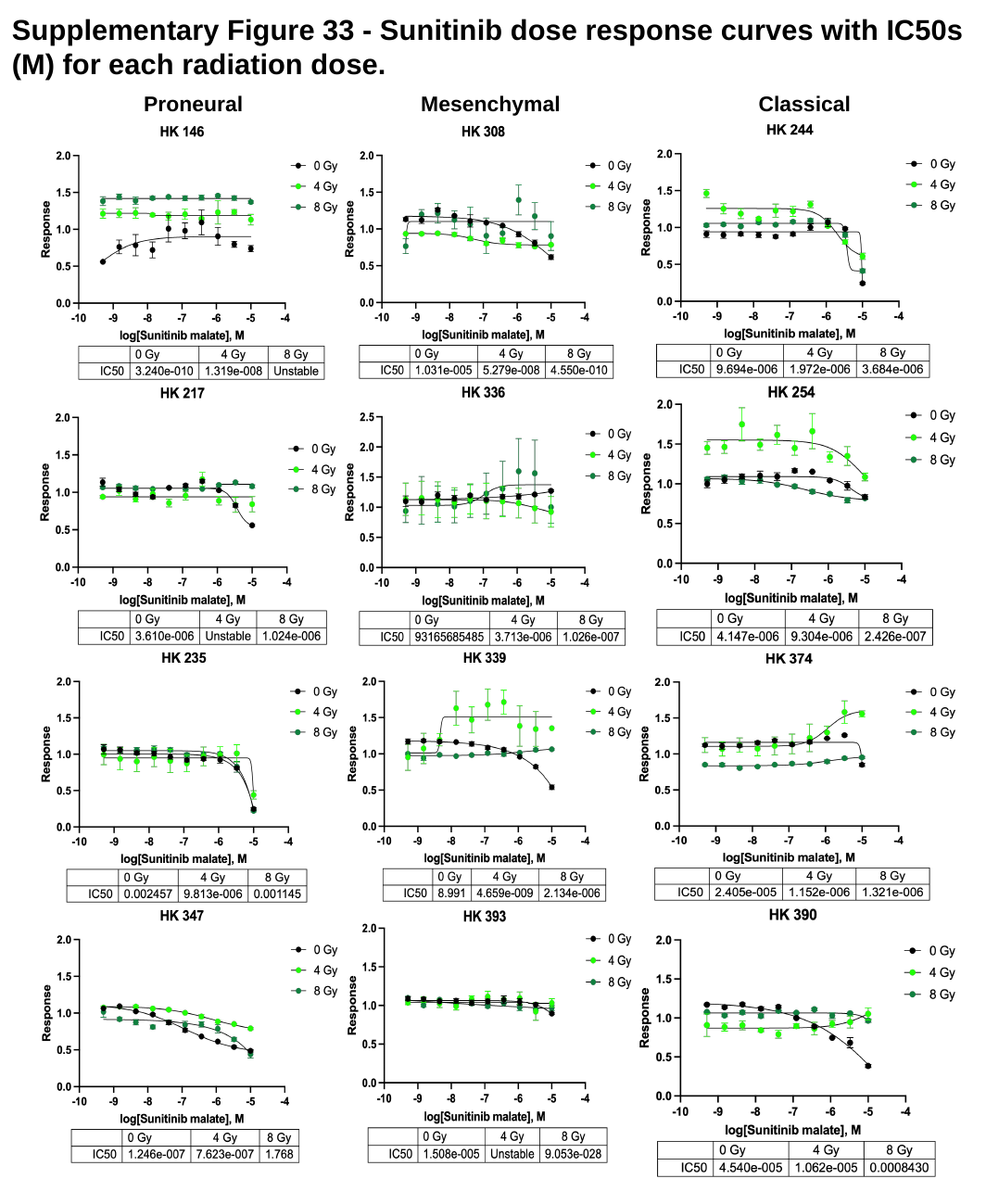

#### Supplementary Figure 33 - Sunitinib dose response curves with IC50s (M) for each radiation dose.
Classical
Mesenchymal
Proneural

### Slide 34
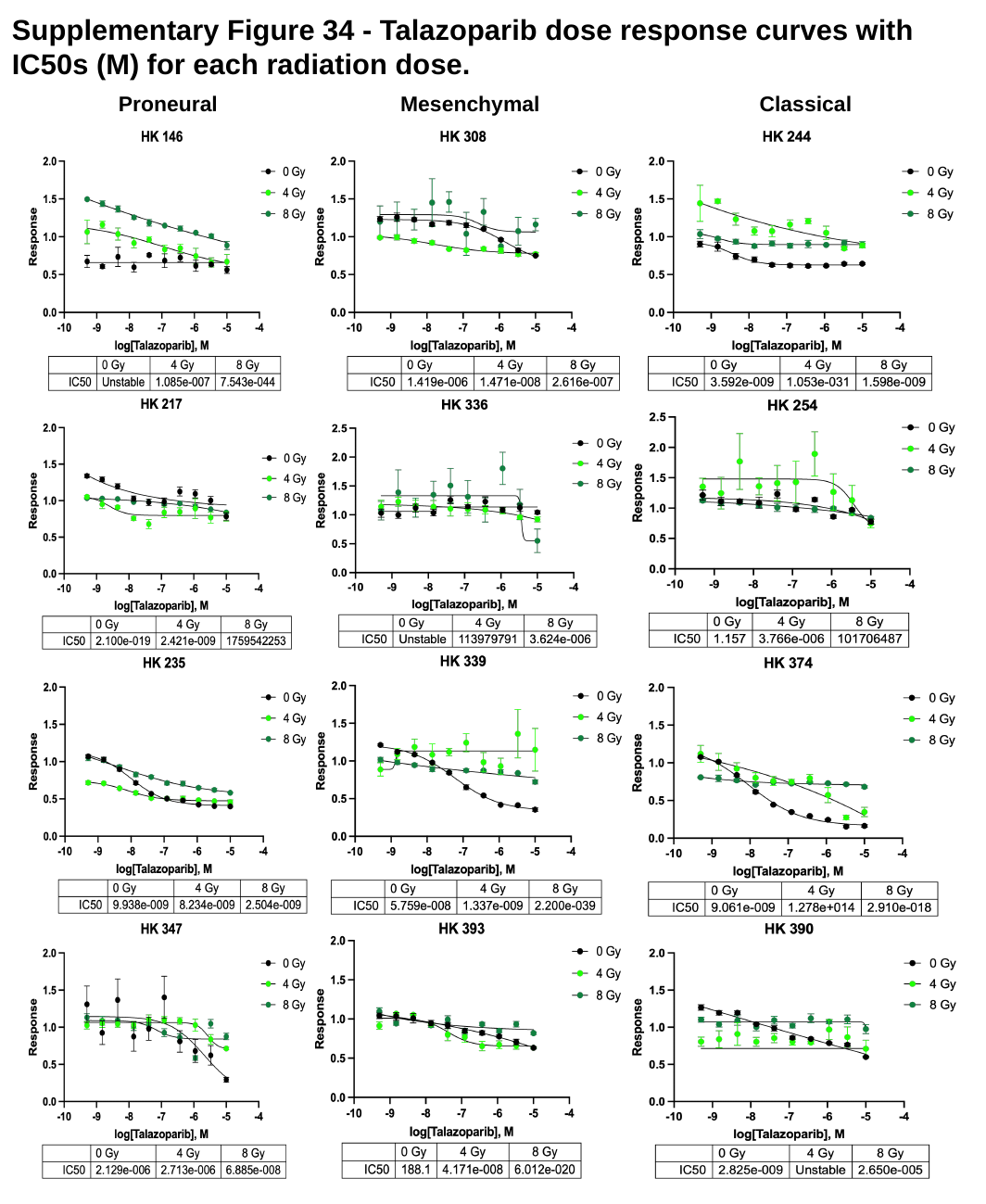

#### Supplementary Figure 34 - Talazoparib dose response curves with IC50s (M) for each radiation dose.
Mesenchymal
Classical
Proneural

### Slide 35
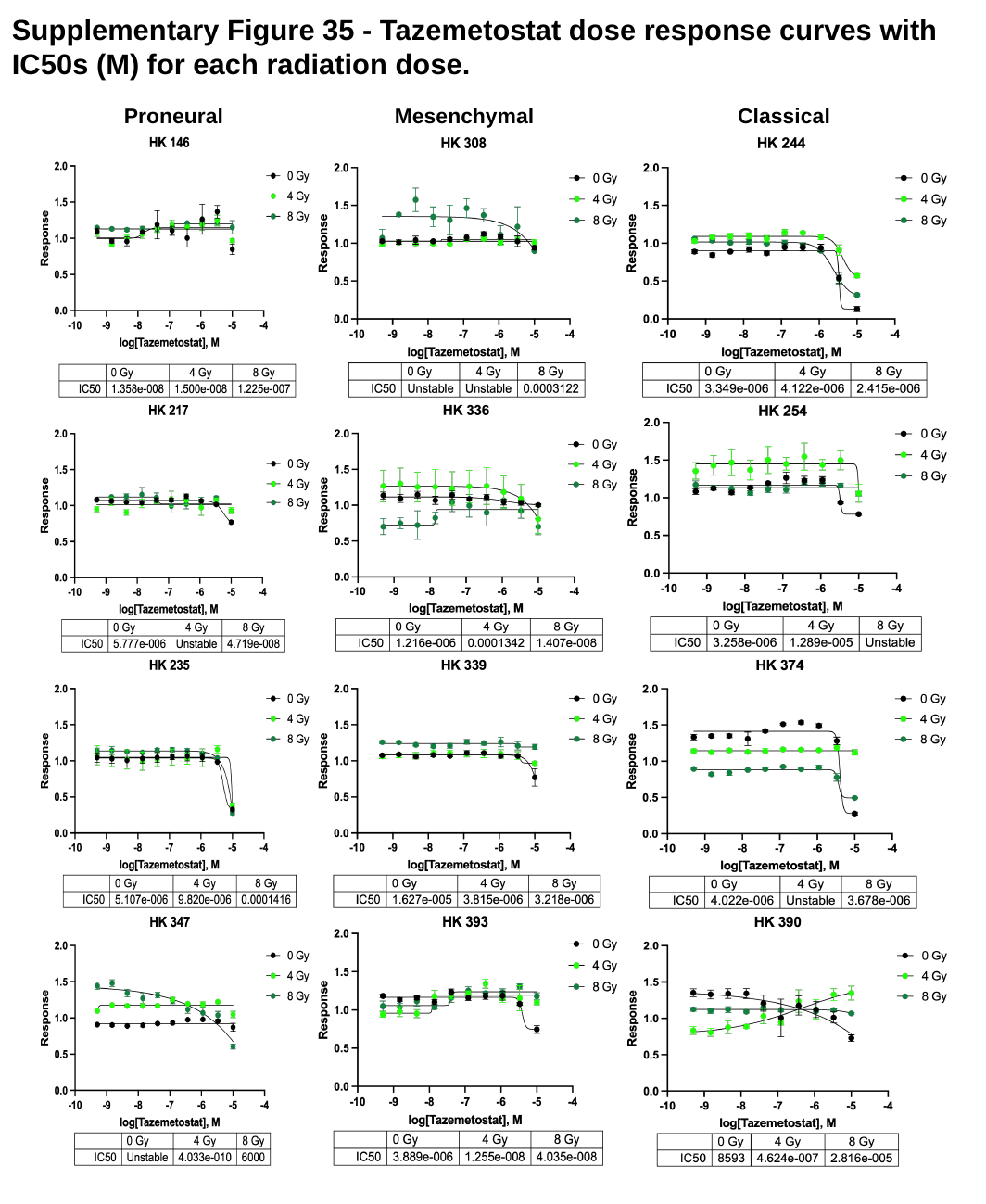

#### Supplementary Figure 35 - Tazemetostat dose response curves with IC50s (M) for each radiation dose.
Mesenchymal
Proneural
Classical

### Slide 36
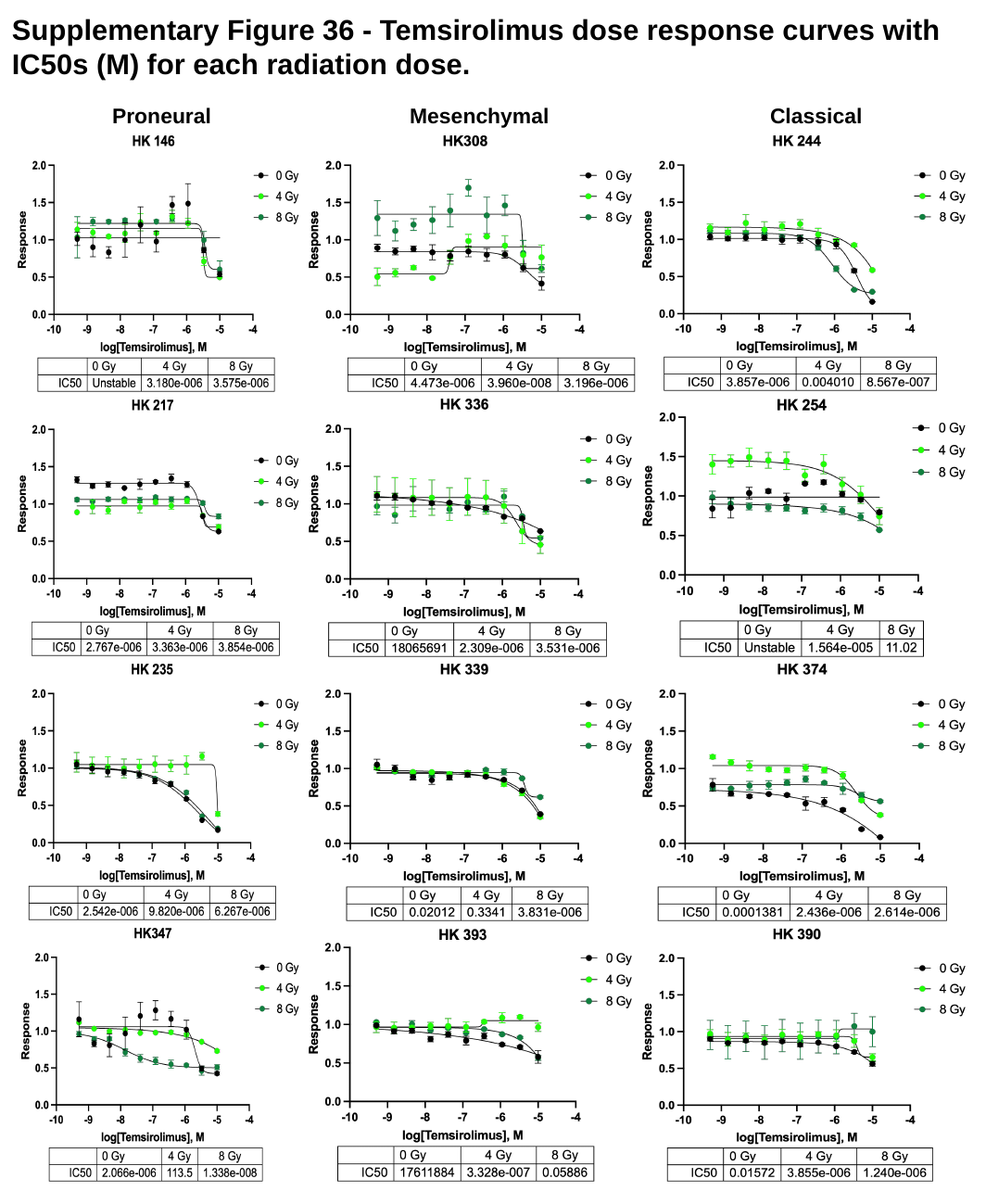

#### Supplementary Figure 36 - Temsirolimus dose response curves with IC50s (M) for each radiation dose.
Mesenchymal
Proneural
Classical

### Slide 37
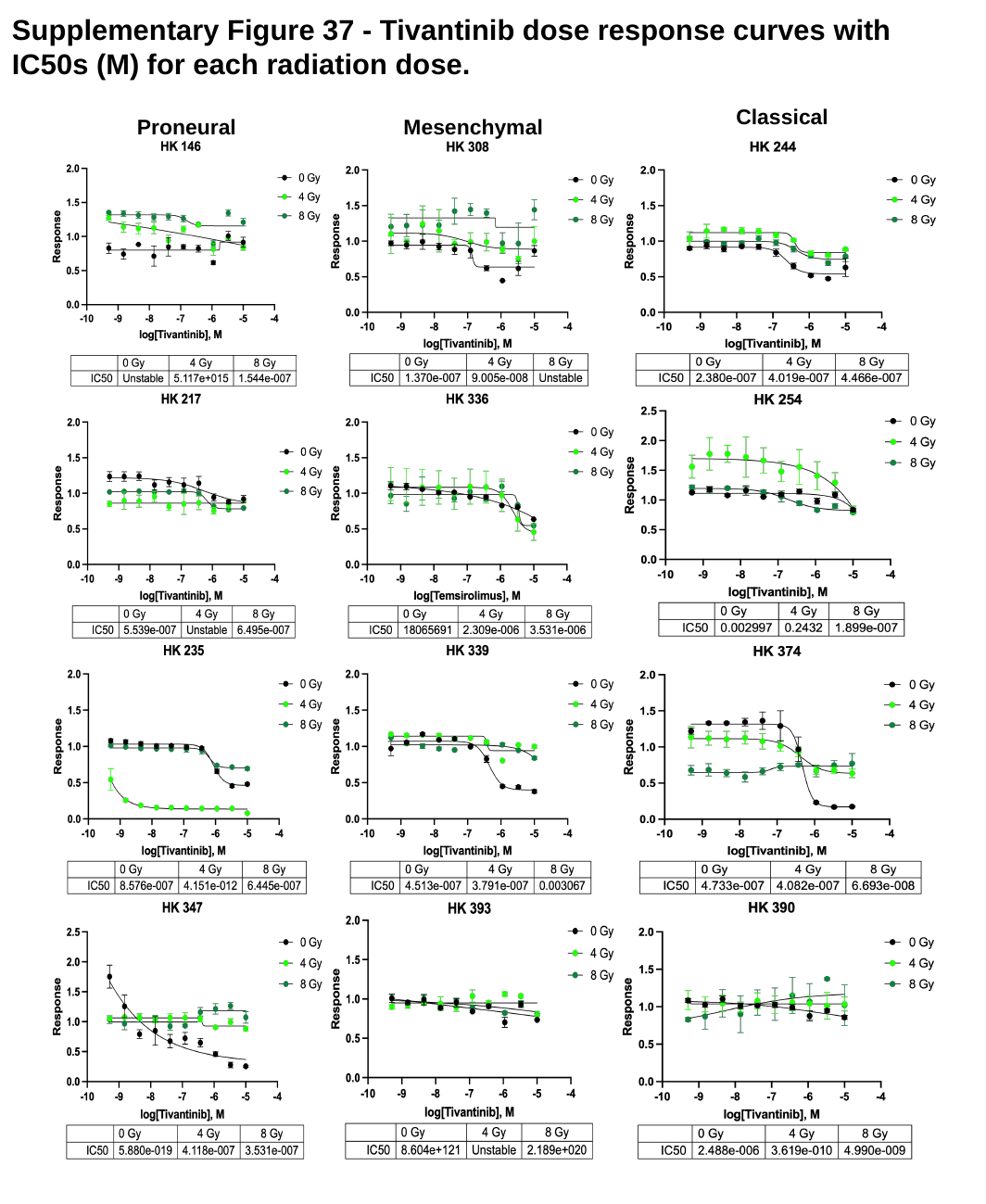

#### Supplementary Figure 37 - Tivantinib dose response curves with IC50s (M) for each radiation dose.
Classical
Mesenchymal
Proneural

### Slide 38
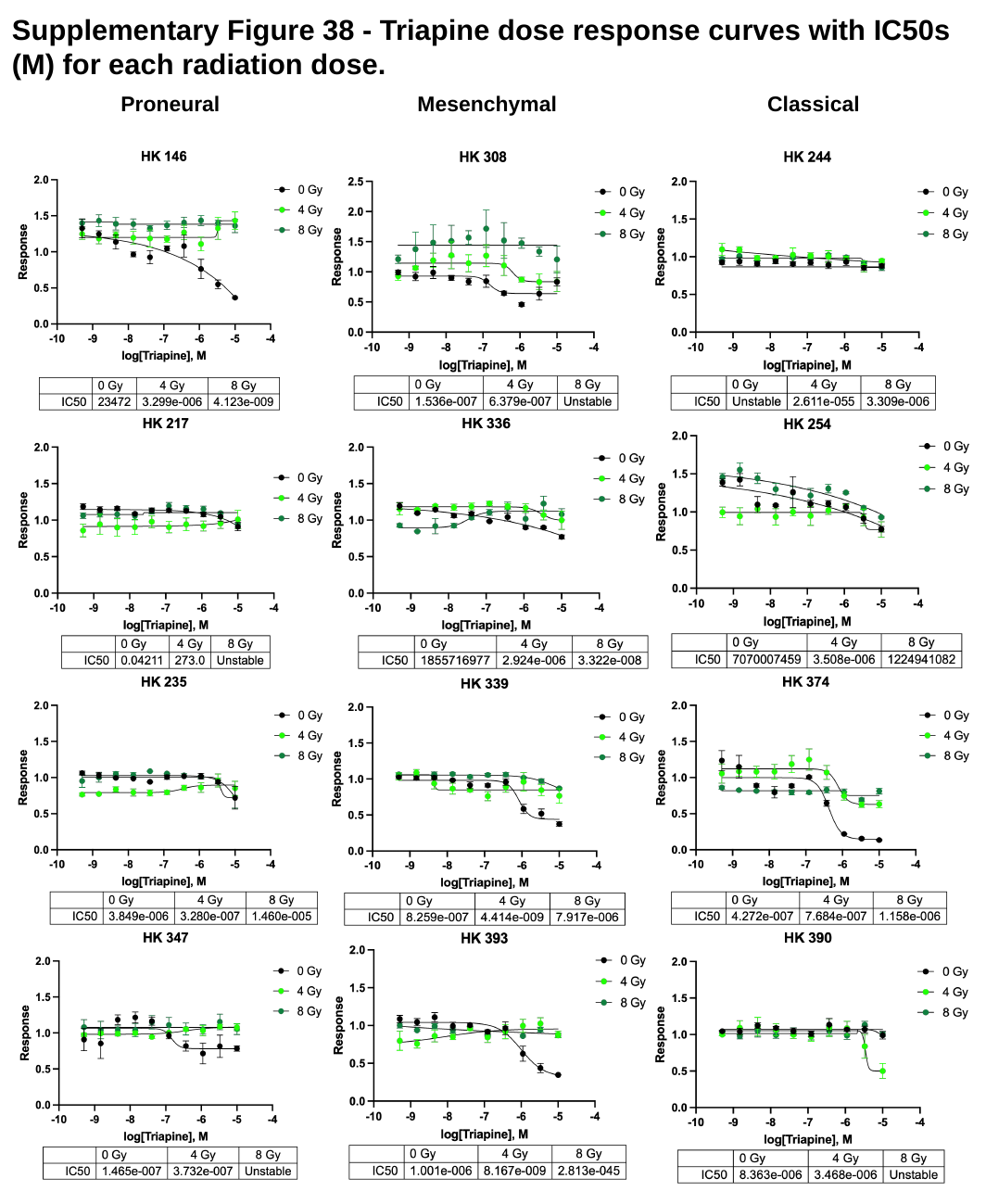

#### Supplementary Figure 38 - Triapine dose response curves with IC50s (M) for each radiation dose.
Mesenchymal
Classical
Proneural

### Slide 39
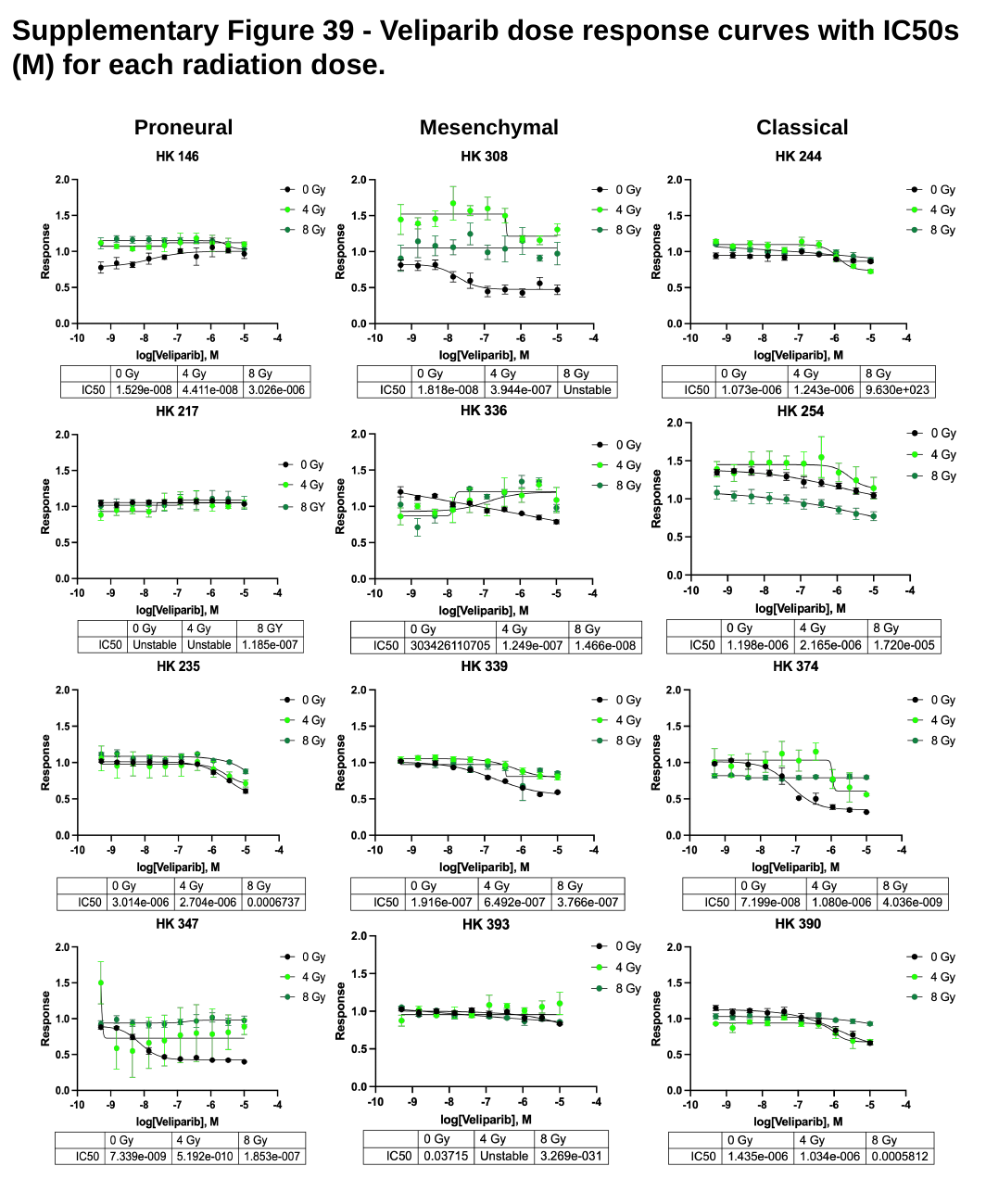

#### Supplementary Figure 39 - Veliparib dose response curves with IC50s (M) for each radiation dose.
Mesenchymal
Classical
Proneural

### Slide 40
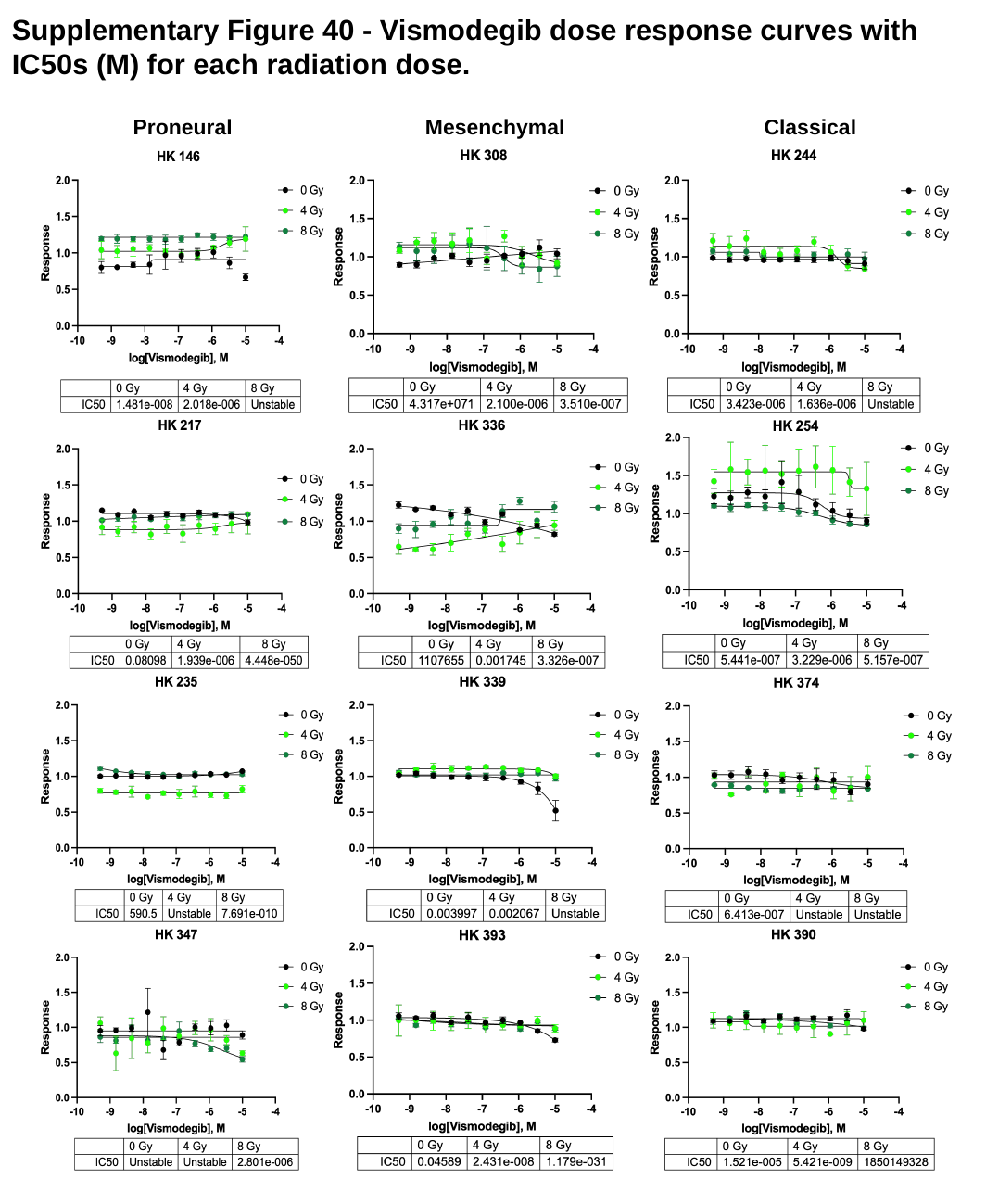

#### Supplementary Figure 40 - Vismodegib dose response curves with IC50s (M) for each radiation dose.
Mesenchymal
Classical
Proneural

### Slide 41
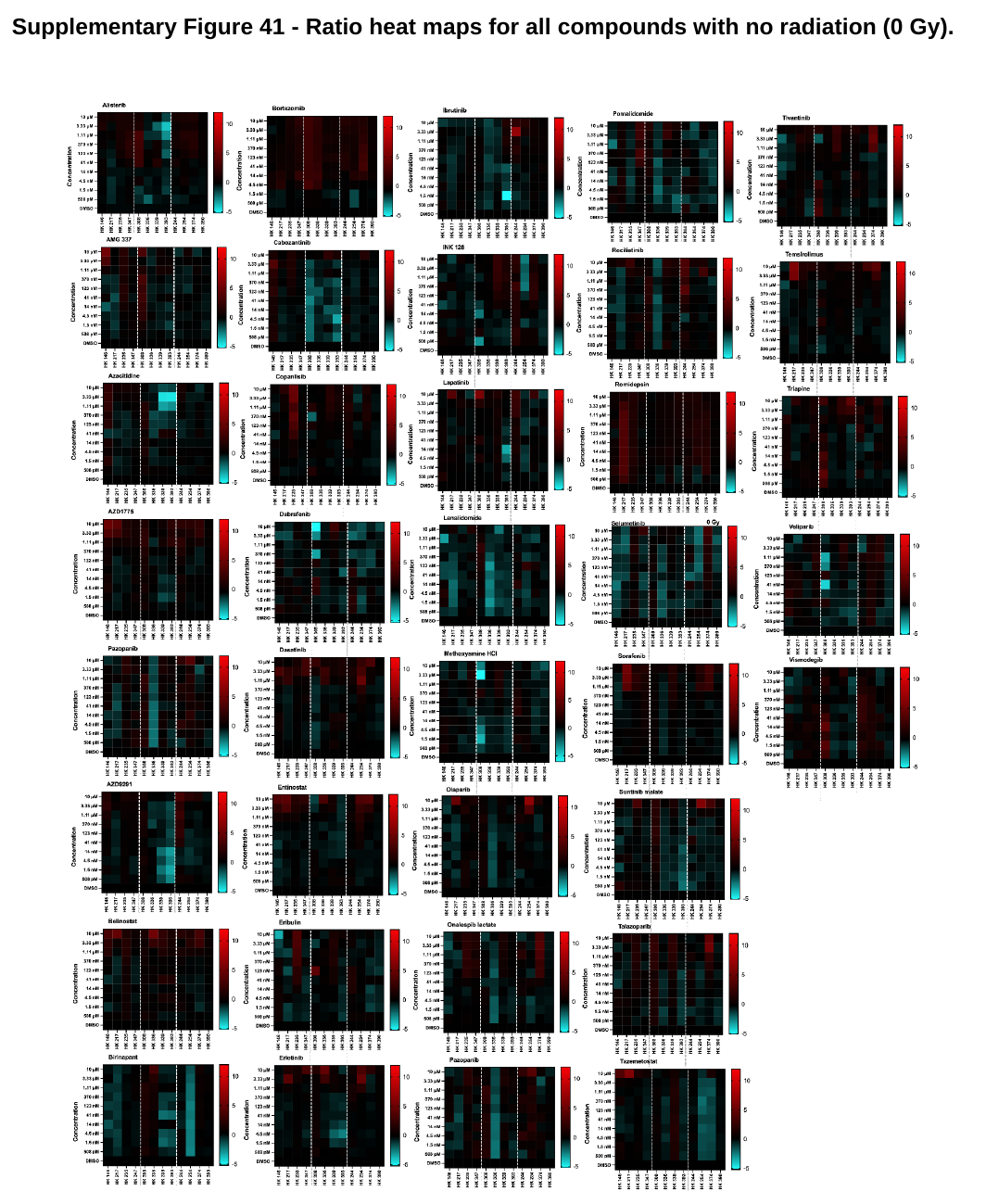

#### Supplementary Figure 41 - Ratio heat maps for all compounds with no radiation (0 Gy).

### Slide 42
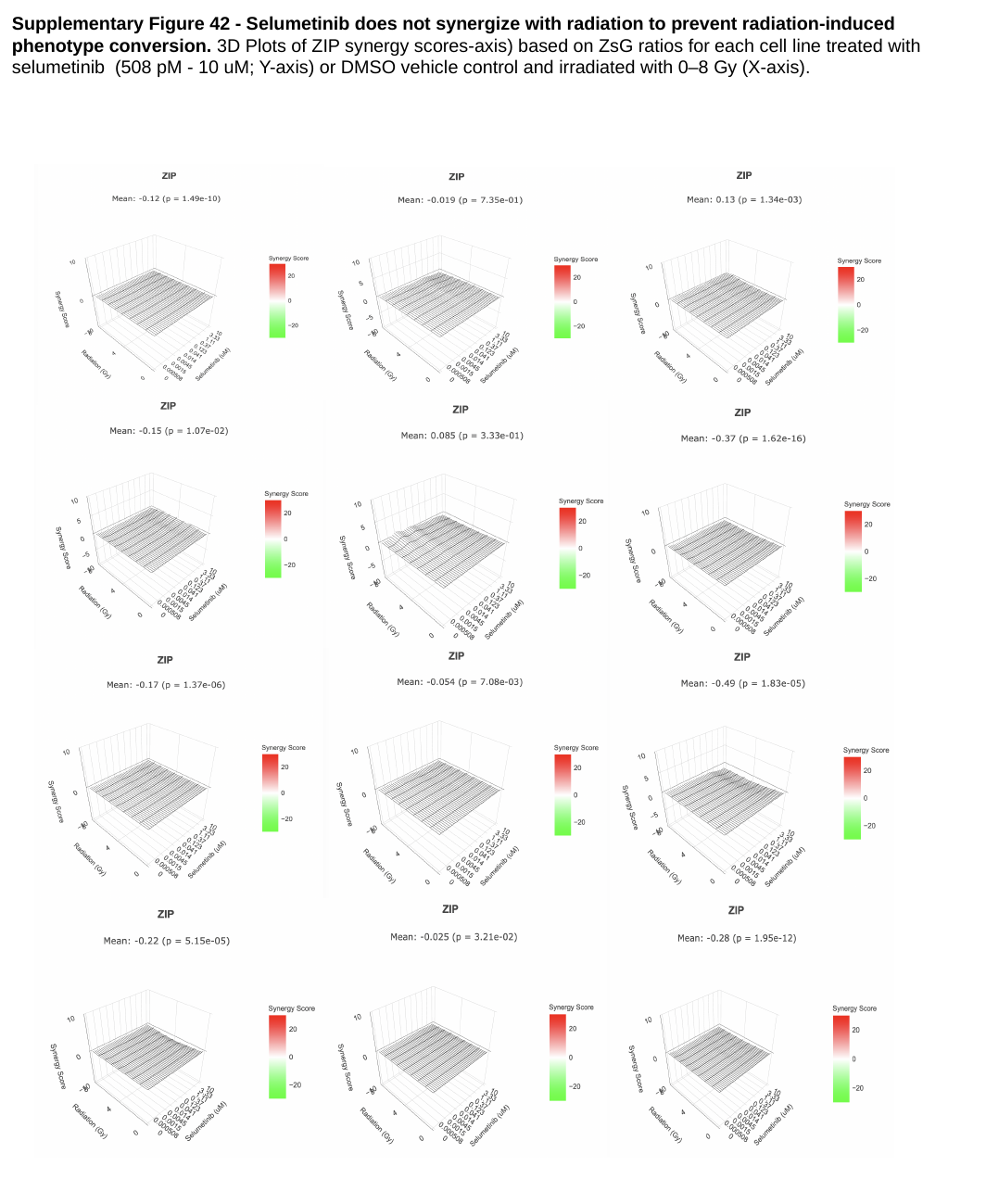

#### Supplementary Figure 42 - Selumetinib does not synergize with radiation to prevent radiation-induced phenotype conversion. 3D Plots of ZIP synergy scores-axis) based on ZsG ratios for each cell line treated with selumetinib (508 pM - 10 uM; Y-axis) or DMSO vehicle control and irradiated with 0–8 Gy (X-axis).
