## Supplementary Table 1 for "Targeting Radiation-Induced Glioma-Initiating Cells in Patient-Derived Glioblastoma"

**Supplementary Table 1.** CTEP portfolio agents used in study.

| Compound | Target | Vendor | Location |
|---|---|---|---|
| Alisertib (MLN8237) | Aurora Kinase A | AdooQ BioScience | Irvine, CA |
| AMG337 | MET kinase inhibitor | ChemieTek | Indianapolis, IN |
| Azacitidine | DNA methyltransferase (DNMTs) | AdooQ BioScience | Irvine, CA |
| AZD 1775 | WEE1, polo-like kinase1 (PLK1) | MedChem Express | Monmouth Junction, NJ |
| AZD 8186 | phosphatidylinositol 3-kinase (PI3K)β and PI3Kδ | MedChem Express | Monmouth Junction, NJ |
| AZD 9291 (Osimertinib) | Epidermal growth factor receptor (EGFR) | AdooQ Bioscience | Irvine, CA |
| Belinostat (PXD 101) | Histone deacetylases (HDAC) | AdooQ BioScience | Irvine, CA |
| Birinapant (TL32711) | SMAC mimetic; Inhibitor of Apoptosis Protein (IAP) | MedChem Express | Monmouth Junction, NJ |
| Bortezomib (PS-341; Velclade) | 26S proteasome | AdooQ BioScience | Irvine, CA |
| Cabozantinib (XL-184) | c-Met, VEGFR2, AXL, RET, platelet-derived growth factor receptor (PDGFR), c-KIT, Bcr-Abl/Src | AdooQ BioScience | Irvine, CA |
| Copanlisib | PI3K | Selleck Chemicals | Houston, Texas |
| Dabrafenib mesylate (GSK2118436B) | BRAF | AdooQ BioScience | Irvine, CA |
| Dasatinib (BMS-354825; Sprycel) | BCR-ABL, SRC family (SRC, LCK, YES, FYN), c-KIT, EPHA2, and PDGFRβ | AdooQ BioScience | Irvine, CA |
| Entinostat (MS-275, SNDX-275) | HDAC1 | AdooQ BioScience | Irvine, CA |
| Eribulin | Tubulin beta-1 chain | AdooQ BioScience | Irvine, CA |
| Erlotinib (OSL-774; Tarceva) | EGFR | MedChem Express | Monmouth Junction, NJ |

| | | | |
|---|---|---|---|
| Ibrutinib (PCL-32765) | Bruton's tyrosine kinase (BTK) | MedChem Express | Monmouth Junction, NJ |
| INK128 (MLN0128) | mTOR1/2 | Selleck Chemicals | Houston, Texas |
| Lapatinib (GW6572016) | HER2 and EGFR | AdooQ BioScience | Irvine, CA |
| Lenalidomide (CC-5013, Revlimid) | Cereblon (CRBN) | AdooQ BioScience | Irvine, CA |
| Methoxyamine HCl (TRC-102) | apyrimidinic (AP) | Sigma Aldrich | St. Louis, MO |
| Olaparib (AZD-2281) | PARP inhibitor | AdooQ BioScience | Irvine, CA |
| Onalespib lactate (AT13387) | heat shock protein 90 (Hsp90) | Biosynth International, Inc | Gardner, MA |
| Pazopanib (GW786034) | VEGF | MedChem Express | Monmouth Junction, NJ |
| Pomalidomide (CC-4047) | Cereblon, COX2 | AdooQ BioScience | Irvine, CA |
| Rociletinib (CO-1686) | EGFR | MedChem Express | Monmouth Junction, NJ |
| Romidepsin (Depsipeptide; FK228) | HDAC | AdooQ BioScience | Irvine, CA |
| Selumetinib (AZD6244) | MEK1 and MEK2 | MedChem Express | Monmouth Junction, NJ |
| Sorafenib tosylate (Bay 43-9006 Tosylate; BAY 54-9085) | RAF, VEGFR, PDGFR, FGFR, RET, c-KIT, Fms-related tyrosine kinase 3 (FLT3) | MedChem Express | Monmouth Junction, NJ |
| Sunitinib malate (SU011248 L-malate; Sutent) | VEGFR, PDGFR, c-KIT, FLT3 | AdooQ BioScience | Irvine, CA |
| Talazoparib (BMN 673) | poly ADP ribose polymerase inhibitor (PARP) | MedChem Express | Monmouth Junction, NJ |
| Tazemetostat (EPZ-6438; EPZ011989) | EZH2 methyltransferase | AdooQ BioScience | Irvine, CA |
| Temsirolimus (CCI-779, Torisel) | mTOR | AdooQ BioScience | Irvine, CA |

| | | | |
|---|---|---|---|
| Tivantinib (ARQ-197) | MET | MedChem Express | Monmouth Junction, NJ |
| Triapine (3-AP, Pan-811) | ribonucleotide reductase (RNR) | MedChem Express | Monmouth Junction, NJ |
| Veliparib (ABT-888) | PARP | AdooQ BioScience | Irvine, CA |
| Vismodegib (GDC-0449) | SMO | AdooQ BioScience | Irvine, CA |

**Supplementary Table 2.** Two-way ANOVA

| Radiation Dose | Comparison | F-ratio | P-value |
|---|---|---|---|
| 0 Gy | TCGA subtype | F (1.978, 225.5) = 1.385 | 0.2525 |
| | CTEP compound | F (37, 114) = 4.054 | **< 0.0001** |
| | TCGA x CTEP | F (73.18, 225.5) = 0.8140 | 0.8480 |
| 4 Gy | TCGA subtype | F (1.786, 203.6) = 1.437 | 0.2404 |
| | CTEP compound | F (37, 114) = 2.561 | **< 0.0001** |
| | TCGA x CTEP | F (66.08, 203.6) = 0.7943 | 0.8623 |
| 8 Gy | TCGA subtype | F (1.925, 219.5) = 8.798 | **0.0003** |
| | CTEP compound | F (37, 114) = 2.294 | **0.0004** |
| | TCGA x CTEP | F (71.23, 219.5) = 0.9163 | 0.6613 |

**Supplementary Table 3.** Mixed-effects model for 0 Gy

| CTEP Compound | Comparisons | F-ratio | P-value |
|---|---|---|---|
| Alisertib | Concentration | F (1.887, 5.662) = 4.447 | 0.0702 |
| | TCGA | F (1.382, 4.147) = 0.7039 | 0.4945 |
| | Concentration x TCGA | F (1.518, 4.553) = 1.573 | 0.2913 |
| AMG337 | Concentration | F (1.502, 4.506) = 1.903 | 0.2453 |
| | TCGA | F (1.637, 4.910) = 0.7058 | 0.5105 |
| | Concentration x TCGA | F (1.489, 4.467) = 1.190 | 0.3637 |
| Azacitidine | Concentration | F (1.299, 3.896) = 0.6344 | 0.5129 |
| | TCGA | F (1.187, 3.562) = 2.435 | 0.2066 |
| | Concentration x TCGA | F (1.763, 5.290) = 4.677 | 0.0700 |
| AZD 1775 | Concentration | F (2.230, 6.689) = 15.64 | **0.0027** |
| | TCGA | F (1.197, 3.590) = 1.871 | 0.2589 |
| | Concentration x TCGA | F (1.720, 5.160) = 1.587 | 0.2853 |
| AZD 8186 | Concentration | F (2.089, 6.266) = 1.690 | 0.2596 |
| | TCGA | F (1.787, 5.362) = 0.2946 | 0.7339 |
| | Concentration x TCGA | F (1.924, 5.772) = 0.6917 | 0.5326 |
| AZD 9291 | Concentration | F (1.890, 5.670) = 5.620 | **0.0461** |
| | TCGA | F (1.225, 3.674) = 3.259 | 0.1530 |
| | Concentration x TCGA | F (1.524, 4.573) = 1.082 | 0.3895 |
| Belinostat | Concentration | F (1.642, 4.927) = 9.642 | **0.0217** |
| | TCGA | F (1.124, 3.372) = 3.629 | 0.1434 |
| | Concentration x TCGA | F (2.023, 6.069) = 0.5778 | 0.5911 |
| Birinapant | Concentration | F (1.916, 5.747) = 2.406 | 0.1744 |
| | TCGA | F (1.188, 3.565) = 0.3224 | 0.6408 |
| | Concentration x TCGA | F (2.090, 6.269) = 1.311 | 0.3365 |

| | | | |
|---|---|---|---|
| Bortezomib | Concentration | F (2.047, 6.140) = 15.39 | **0.0040** |
| | TCGA | F (1.638, 4.914) = 0.7484 | 0.4946 |
| | Concentration x TCGA | F (2.445, 7.335) = 1.226 | 0.3574 |
| Cabozantinib | Concentration | F (2.040, 6.119) = 7.397 | **0.0230** |
| | TCGA | F (1.034, 3.101) = 5.518 | 0.0975 |
| | Concentration x TCGA | F (2.589, 7.768) = 3.575 | 0.0722 |
| Copanlisib | Concentration | F (1.288, 3.864) = 1.877 | 0.2553 |
| | TCGA | F (1.333, 3.998) = 0.5976 | 0.5298 |
| | Concentration x TCGA | F (2.023, 6.069) = 1.165 | 0.3738 |
| Dabrafenib mesylate | Concentration | F (1.280, 3.841) = 1.012 | 0.4001 |
| | TCGA | F (1.071, 3.213) = 0.8034 | 0.4415 |
| | Concentration x TCGA | F (1.534, 4.603) = 0.4888 | 0.5961 |
| Dasatinib | Concentration | F (1.189, 3.568) = 7.953 | 0.0530 |
| | TCGA | F (1.483, 4.450) = 0.2398 | 0.7360 |
| | Concentration x TCGA | F (2.508, 7.524) = 0.8787 | 0.4768 |
| Entinostat | Concentration | F (1.156, 3.467) = 7.367 | 0.0611 |
| | TCGA | F (1.488, 4.464) = 9.401 | **0.0280** |
| | Concentration x TCGA | F (1.777, 5.330) = 3.342 | 0.1166 |
| Eribulin | Concentration | F (1.512, 4.535) = 0.4903 | 0.5932 |
| | TCGA | F (1.025, 3.075) = 0.1472 | 0.7321 |
| | Concentration x TCGA | F (1.640, 4.921) = 0.8758 | 0.4511 |
| Erlotinib | Concentration | F (1.151, 3.452) = 3.645 | 0.1408 |
| | TCGA | F (1.110, 3.329) = 0.5744 | 0.5162 |
| | Concentration x TCGA | F (1.635, 4.904) = 0.3443 | 0.6855 |
| Ibrutinib | Concentration | F (2.073, 6.220) = 1.978 | 0.2164 |
| | TCGA | F (1.257, 3.770) = 11.30 | **0.0290** |

| | | | |
|---|---|---|---|
| | Concentration x TCGA | F (1.639, 4.917) = 1.376 | 0.3250 |
| INK128 | Concentration | F (1.886, 5.659) = 0.7887 | 0.4917 |
| | TCGA | F (1.192, 3.577) = 0.3994 | 0.6006 |
| | Concentration x TCGA | F (2.287, 6.862) = 2.724 | 0.1318 |
| Lapatinib | Concentration | F (1.410, 4.231) = 2.843 | 0.1652 |
| | TCGA | F (1.765, 5.294) = 0.03409 | 0.9542 |
| | Concentration x TCGA | F (2.141, 6.423) = 0.5940 | 0.5903 |
| Lenalidomide | Concentration | F (1.537, 4.612) = 0.7570 | 0.4859 |
| | TCGA | F (1.301, 3.903) = 0.8826 | 0.4341 |
| | Concentration x TCGA | F (1.916, 5.749) = 1.537 | 0.2906 |
| Methoxyamine HCl | Concentration | F (1.248, 3.743) = 1.079 | 0.3836 |
| | TCGA | F (1.118, 3.353) = 0.9646 | 0.4044 |
| | Concentration x TCGA | F (1.580, 4.739) = 1.277 | 0.3453 |
| Olaparib | Concentration | F (1.130, 3.391) = 1.729 | 0.2773 |
| | TCGA | F (1.532, 4.595) = 0.6753 | 0.5154 |
| | Concentration x TCGA | F (1.731, 5.193) = 1.139 | 0.3784 |
| Onalespib lactate | Concentration | F (1.495, 4.485) = 1.197 | 0.3621 |
| | TCGA | F (1.663, 4.989) = 1.375 | 0.3251 |
| | Concentration x TCGA | F (2.370, 7.111) = 1.037 | 0.4155 |
| Pazopanib | Concentration | F (2.370, 7.110) = 1.279 | 0.3432 |
| | TCGA | F (1.410, 4.230) = 0.3036 | 0.6835 |
| | Concentration x TCGA | F (2.745, 8.234) = 1.127 | 0.3881 |
| Pomalidomide | Concentration | F (2.149, 6.448) = 1.825 | 0.2361 |
| | TCGA | F (1.204, 3.612) = 0.6917 | 0.4850 |
| | Concentration x TCGA | F (2.805, 8.414) = 1.678 | 0.2448 |
| Rociletinib | Concentration | F (2.506, 7.518) = 2.295 | 0.1628 |

| | | | |
|---|---|---|---|
| | TCGA | F (1.680, 5.040) = 0.7549 | 0.4945 |
| | Concentration x TCGA | F (2.055, 6.164) = 1.077 | 0.3992 |
| Romidepsin | Concentration | F (1.658, 4.974) = 4.162 | 0.0898 |
| | TCGA | F (1.168, 3.504) = 1.516 | 0.3048 |
| | Concentration x TCGA | F (1.885, 5.656) = 1.132 | 0.3817 |
| Selumetinib | Concentration | F (2.130, 6.391) = 0.8282 | 0.4859 |
| | TCGA | F (1.535, 4.606) = 0.1725 | 0.7947 |
| | Concentration x TCGA | F (2.490, 7.469) = 0.8253 | 0.4985 |
| Sorafenib tosylate | Concentration | F (1.128, 3.384) = 4.570 | 0.1117 |
| | TCGA | F (1.108, 3.325) = 0.8101 | 0.4424 |
| | Concentration x TCGA | F (1.449, 4.347) = 1.028 | 0.4012 |
| Sunitinib malate | Concentration | F (1.585, 4.755) = 6.419 | 0.0479 |
| | TCGA | F (1.120, 3.359) = 0.1315 | 0.7653 |
| | Concentration x TCGA | F (1.957, 5.870) = 1.780 | 0.2484 |
| Talazoparib | Concentration | F (1.339, 4.018) = 2.343 | 0.2065 |
| | TCGA | F (1.401, 4.203) = 0.1131 | 0.8298 |
| | Concentration x TCGA | F (1.837, 5.511) = 0.8678 | 0.4611 |
| Tazemetostat | Concentration | F (1.400, 4.201) = 6.441 | 0.0570 |
| | TCGA | F (1.679, 5.038) = 2.039 | 0.2227 |
| | Concentration x TCGA | F (1.840, 5.521) = 3.413 | 0.1099 |
| Temsirolimus | Concentration | F (1.220, 3.661) = 6.029 | 0.0742 |
| | TCGA | F (1.812, 5.437) = 0.02329 | 0.9694 |
| | Concentration x TCGA | F (1.755, 5.265) = 0.9432 | 0.4338 |
| Tivantinib | Concentration | F (1.372, 4.115) = 1.325 | 0.3356 |
| | TCGA | F (1.088, 3.263) = 2.395 | 0.2148 |
| | Concentration x TCGA | F (2.152, 6.456) = 0.9235 | 0.4509 |

| | | | |
|---|---|---|---|
| Triapine | Concentration | F (1.358, 4.074) = 2.322 | 0.2074 |
| | TCGA | F (1.403, 4.209) = 0.9753 | 0.4134 |
| | Concentration x TCGA | F (2.349, 7.047) = 0.7726 | 0.5167 |
| Veliparib | Concentration | F (1.422, 4.265) = 0.4310 | 0.6131 |
| | TCGA | F (1.531, 4.592) = 2.038 | 0.2284 |
| | Concentration x TCGA | F (1.897, 5.690) = 1.092 | 0.3929 |
| Vismodegib | Concentration | F (1.552, 4.655) = 1.636 | 0.2808 |
| | TCGA | F (1.484, 4.451) = 0.3816 | 0.6466 |
| | Concentration x TCGA | F (1.797, 5.390) = 0.7142 | 0.5164 |

**Supplementary Table 4.** Mixed-effects model for 4 Gy

| CTEP Compound | Comparisons | F-ratio | P-value |
|---|---|---|---|
| Alisertib | Concentration | F (2.257, 6.770) = 2.924 | 0.1188 |
| | TCGA | F (1.116, 3.349) = 0.3249 | 0.6282 |
| | Concentration x TCGA | F (1.622, 4.865) = 1.021 | 0.4075 |
| AMG337 | Concentration | F (1.015, 3.044) = 1.060 | 0.3796 |
| | TCGA | F (1.022, 3.065) = 0.8851 | 0.4176 |
| | Concentration x TCGA | F (1.013, 3.040) = 0.9730 | 0.3974 |
| Azacitidine | Concentration | F (1.739, 5.218) = 0.6161 | 0.5541 |
| | TCGA | F (1.072, 3.215) = 0.4585 | 0.5570 |
| | Concentration x TCGA | F (1.682, 5.046) = 0.6047 | 0.5551 |
| AZD 1775 | Concentration | F (2.619, 7.858) = 2.067 | 0.1867 |
| | TCGA | F (1.342, 4.027) = 0.6774 | 0.5009 |
| | Concentration x TCGA | F (2.417, 7.252) = 1.979 | 0.2051 |
| AZD 8186 | Concentration | F (1.129, 3.387) = 1.474 | 0.3112 |
| | TCGA | F (1.008, 3.024) = 1.005 | 0.3904 |
| | Concentration x TCGA | F (1.155, 3.465) = 1.086 | 0.3794 |
| AZD 9291 | Concentration | F (2.003, 6.010) = 0.9218 | 0.4477 |
| | TCGA | F (1.181, 3.544) = 0.2985 | 0.6536 |
| | Concentration x TCGA | F (2.050, 6.149) = 1.169 | 0.3729 |
| Belinostat | Concentration | F (1.045, 3.136) = 3.024 | 0.1773 |
| | TCGA | F (1.609, 4.826) = 0.3267 | 0.6934 |
| | Concentration x TCGA | F (1.305, 3.915) = 0.6467 | 0.5089 |
| Birinapant | Concentration | F (1.923, 5.770) = 2.456 | 0.1695 |
| | TCGA | F (1.008, 3.024) = 0.4991 | 0.5319 |

| | | | |
|---|---|---|---|
| | Concentration x TCGA | F (1.540, 4.620) = 0.9137 | 0.4356 |
| Bortezomib | Concentration | F (1.047, 3.141) = 1.789 | 0.2723 |
| | TCGA | F (1.155, 3.465) = 0.5648 | 0.5248 |
| | Concentration x TCGA | F (1.130, 3.390) = 0.6898 | 0.4793 |
| Cabozantinib | Concentration | F (2.200, 6.599) = 1.209 | 0.3625 |
| | TCGA | F (1.198, 3.594) = 1.609 | 0.2913 |
| | Concentration x TCGA | F (2.696, 8.088) = 1.332 | 0.3267 |
| Copanlisib | Concentration | F (1.073, 3.220) = 1.042 | 0.3856 |
| | TCGA | F (1.099, 3.297) = 0.6551 | 0.4875 |
| | Concentration x TCGA | F (1.243, 3.730) = 1.326 | 0.3349 |
| Dabrafenib mesylate | Concentration | F (1.032, 3.097) = 1.040 | 0.3843 |
| | TCGA | F (1.080, 3.239) = 1.211 | 0.3533 |
| | Concentration x TCGA | F (1.030, 3.090) = 0.9755 | 0.3977 |
| Dasatinib | Concentration | F (1.517, 4.550) = 7.088 | **0.0436** |
| | TCGA | F (1.001, 3.004) = 1.690 | 0.2845 |
| | Concentration x TCGA | F (1.535, 4.605) = 3.407 | 0.1265 |
| Entinostat | Concentration | F (1.007, 3.020) = 1.819 | 0.2700 |
| | TCGA | F (1.006, 3.017) = 1.054 | 0.3803 |
| | Concentration x TCGA | F (1.059, 3.178) = 1.116 | 0.3703 |
| Eribulin | Concentration | F (1.407, 4.222) = 1.271 | 0.3459 |
| | TCGA | F (1.034, 3.103) = 0.1639 | 0.7198 |
| | Concentration x TCGA | F (1.368, 4.104) = 0.8931 | 0.4343 |
| Erlotinib | Concentration | F (1.947, 5.841) = 6.270 | **0.0356** |
| | TCGA | F (1.194, 3.581) = 1.699 | 0.2796 |
| | Concentration x TCGA | F (2.028, 6.085) = 2.436 | 0.1669 |
| Ibrutinib | Concentration | F (2.521, 7.562) = 0.9576 | 0.4461 |

| | | | |
|---|---|---|---|
| | TCGA | F (1.024, 3.071) = 0.4215 | 0.5661 |
| | Concentration x TCGA | F (2.663, 7.989) = 1.197 | 0.3656 |
| INK128 | Concentration | F (1.981, 5.944) = 0.8007 | 0.4911 |
| | TCGA | F (1.050, 3.150) = 0.3150 | 0.6226 |
| | Concentration x TCGA | F (1.679, 5.036) = 0.8902 | 0.4479 |
| Lapatinib | Concentration | F (1.292, 3.876) = 1.614 | 0.2891 |
| | TCGA | F (1.124, 3.373) = 1.252 | 0.3469 |
| | Concentration x TCGA | F (1.283, 3.848) = 0.7418 | 0.4747 |
| Lenalidomide | Concentration | F (1.447, 4.342) = 4.225 | 0.0999 |
| | TCGA | F (1.037, 3.110) = 0.9125 | 0.4121 |
| | Concentration x TCGA | F (2.383, 7.148) = 2.794 | 0.1228 |
| Methoxyamine HCl | Concentration | F (1.021, 3.063) = 0.9516 | 0.4024 |
| | TCGA | F (1.018, 3.054) = 0.7016 | 0.4654 |
| | Concentration x TCGA | F (1.021, 3.064) = 0.8224 | 0.4329 |
| Olaparib | Concentration | F (2.681, 8.042) = 0.8224 | 0.5053 |
| | TCGA | F (1.170, 3.511) = 1.808 | 0.2669 |
| | Concentration x TCGA | F (2.224, 6.673) = 1.014 | 0.4213 |
| Onalespib lactate | Concentration | F (2.589, 7.766) = 2.931 | 0.1057 |
| | TCGA | F (1.433, 4.299) = 0.002285 | 0.9894 |
| | Concentration x TCGA | F (1.914, 5.741) = 0.6466 | 0.5518 |
| Pazopanib | Concentration | F (1.995, 5.984) = 0.6241 | 0.5669 |
| | TCGA | F (1.246, 3.739) = 1.106 | 0.3777 |
| | Concentration x TCGA | F (2.682, 8.047) = 1.217 | 0.3597 |
| Pomalidomide | Concentration | F (1.914, 5.741) = 0.8221 | 0.4803 |
| | TCGA | F (1.419, 4.258) = 1.907 | 0.2476 |
| | Concentration x TCGA | F (1.717, 5.152) = 1.363 | 0.3270 |

| | | | |
|---|---|---|---|
| Rociletinib | Concentration | F (2.019, 6.058) = 3.704 | 0.0888 |
| | TCGA | F (1.148, 3.445) = 0.8524 | 0.4338 |
| | Concentration x TCGA | F (2.036, 6.108) = 4.591 | 0.0603 |
| Romidepsin | Concentration | F (2.262, 6.785) = 2.623 | 0.1409 |
| | TCGA | F (1.248, 3.743) = 1.274 | 0.3444 |
| | Concentration x TCGA | F (1.533, 4.600) = 1.424 | 0.3167 |
| Selumetinib | Concentration | F (2.291, 6.874) = 0.7494 | 0.5246 |
| | TCGA | F (1.007, 3.022) = 5.554 | 0.0991 |
| | Concentration x TCGA | F (2.626, 7.879) = 5.699 | **0.0245** |
| Sorafenib tosylate | Concentration | F (1.071, 3.214) = 3.868 | 0.1382 |
| | TCGA | F (1.014, 3.042) = 0.9341 | 0.4059 |
| | Concentration x TCGA | F (1.974, 5.921) = 1.705 | 0.2600 |
| Sunitinib malate | Concentration | F (1.123, 3.370) = 4.596 | 0.1113 |
| | TCGA | F (1.383, 4.150) = 4.766 | 0.0886 |
| | Concentration x TCGA | F (1.453, 4.360) = 0.9044 | 0.4348 |
| Talazoparib | Concentration | F (2.032, 6.095) = 2.334 | 0.1766 |
| | TCGA | F (1.592, 4.775) = 0.001975 | 0.9939 |
| | Concentration x TCGA | F (1.475, 4.425) = 1.078 | 0.3895 |
| Tazemetostat | Concentration | F (2.262, 6.785) = 2.623 | 0.1409 |
| | TCGA | F (1.248, 3.743) = 1.274 | 0.3444 |
| | Concentration x TCGA | F (1.533, 4.600) = 1.424 | 0.3167 |
| Temsirolimus | Concentration | F (1.051, 3.153) = 1.254 | 0.3454 |
| | TCGA | F (1.230, 3.689) = 1.216 | 0.3550 |
| | Concentration x TCGA | F (1.269, 3.807) = 1.093 | 0.3811 |
| Tivantinib | Concentration | F (1.113, 3.339) = 1.389 | 0.3240 |
| | TCGA | F (1.118, 3.353) = 0.8424 | 0.4345 |

| | | | |
|---|---|---|---|
| | Concentration x TCGA | F (1.214, 3.642) = 0.8072 | 0.4503 |
| Triapine | Concentration | F (1.116, 3.347) = 0.7897 | 0.4485 |
| | TCGA | F (1.311, 3.932) = 0.4042 | 0.6137 |
| | Concentration x TCGA | F (1.184, 3.553) = 1.138 | 0.3695 |
| Veliparib | Concentration | F (1.292, 3.876) = 1.353 | 0.3303 |
| | TCGA | F (1.142, 3.426) = 8.033 | 0.0554 |
| | Concentration x TCGA | F (1.358, 4.074) = 1.585 | 0.2922 |
| Vismodegib | Concentration | F (1.187, 3.561) = 2.567 | 0.1968 |
| | TCGA | F (1.466, 4.397) = 4.374 | 0.0942 |
| | Concentration x TCGA | F (1.280, 3.839) = 1.307 | 0.3385 |

**Supplementary Table 5.** Mixed-effects model for 8 Gy.

| CTEP Compound | Comparisons | F-ratio | P-value |
|---|---|---|---|
| Alisertib | Concentration | F (1.477, 4.430) = 1.492 | 0.3053 |
| | TCGA | F (1.665, 4.995) = 1.901 | 0.2402 |
| | Concentration x TCGA | F (2.469, 7.406) = 0.9553 | 0.4460 |
| AMG337 | Concentration | F (2.170, 6.509) = 1.172 | 0.3726 |
| | TCGA | F (1.940, 5.820) = 2.474 | 0.1672 |
| | Concentration x TCGA | F (2.609, 7.828) = 0.5531 | 0.6389 |
| Azacitidine | Concentration | F (1.436, 4.308) = 0.9142 | 0.4313 |
| | TCGA | F (1.253, 3.759) = 2.529 | 0.1959 |
| | Concentration x TCGA | F (2.944, 8.832) = 1.774 | 0.2233 |
| AZD 1775 | Concentration | F (1.229, 3.687) = 0.2619 | 0.6839 |
| | TCGA | F (1.989, 5.967) = 0.7275 | 0.5208 |
| | Concentration x TCGA | F (1.686, 5.059) = 0.7389 | 0.5008 |
| AZD 8186 | Concentration | F (1.104, 3.313) = 1.116 | 0.3716 |
| | TCGA | F (1.731, 5.193) = 1.101 | 0.3881 |
| | Concentration x TCGA | F (1.181, 3.543) = 0.8178 | 0.4452 |
| AZD 9291 | Concentration | F (2.273, 6.818) = 0.7629 | 0.5180 |
| | TCGA | F (1.301, 3.902) = 1.775 | 0.2673 |
| | Concentration x TCGA | F (2.113, 6.338) = 0.9250 | 0.4494 |
| Belinostat | Concentration | F (1.330, 3.990) = 4.294 | 0.1052 |
| | TCGA | F (1.031, 3.094) = 0.01788 | 0.9075 |
| | Concentration x TCGA | F (1.559, 4.677) = 0.6391 | 0.5316 |
| Birinapant | Concentration | F (2.160, 6.480) = 1.164 | 0.3748 |
| | TCGA | F (1.486, 4.457) = 1.267 | 0.3472 |
| | Concentration x TCGA | F (2.103, 6.309) = 1.080 | 0.3988 |

| | | | |
|---|---|---|---|
| Bortezomib | Concentration | F (2.281, 6.842) = 6.697 | **0.0226** |
| | TCGA | F (1.771, 5.313) = 0.9005 | 0.4479 |
| | Concentration x TCGA | F (1.781, 5.342) = 0.6091 | 0.5601 |
| Cabozantinib | Concentration | F (1.682, 5.047) = 0.6335 | 0.5427 |
| | TCGA | F (1.614, 4.843) = 1.095 | 0.3878 |
| | Concentration x TCGA | F (2.513, 7.540) = 0.5024 | 0.6629 |
| Copanlisib | Concentration | F (1.077, 3.231) = 0.5236 | 0.5314 |
| | TCGA | F (1.705, 5.115) = 2.676 | 0.1614 |
| | Concentration x TCGA | F (1.676, 5.029) = 0.9006 | 0.4445 |
| Dabrafenib mesylate | Concentration | F (1.602, 4.807) = 0.7134 | 0.5055 |
| | TCGA | F (1.599, 4.796) = 1.387 | 0.3231 |
| | Concentration x TCGA | F (1.902, 5.705) = 0.3584 | 0.7038 |
| Dasatinib | Concentration | F (2.174, 6.523) = 5.092 | **0.0448** |
| | TCGA | F (1.007, 3.022) = 0.9417 | 0.4038 |
| | Concentration x TCGA | F (2.079, 6.236) = 0.7320 | 0.5231 |
| Entinostat | Concentration | F (1.379, 4.136) = 4.774 | 0.0887 |
| | TCGA | F (1.014, 3.041) = 1.094 | 0.3729 |
| | Concentration x TCGA | F (1.665, 4.994) = 0.9286 | 0.4355 |
| Eribulin | Concentration | F (1.888, 5.665) = 0.7766 | 0.4964 |
| | TCGA | F (1.401, 4.203) = 1.276 | 0.3450 |
| | Concentration x TCGA | F (2.794, 8.381) = 1.299 | 0.3349 |
| Erlotinib | Concentration | F (1.382, 4.145) = 4.572 | 0.0938 |
| | TCGA | F (1.812, 5.436) = 0.5319 | 0.5989 |
| | Concentration x TCGA | F (1.759, 5.277) = 0.5745 | 0.5744 |
| Ibrutinib | Concentration | F (2.075, 6.224) = 1.918 | 0.2246 |
| | TCGA | F (1.233, 3.699) = 0.8083 | 0.4512 |

| | | | |
|---|---|---|---|
| | Concentration x TCGA | F (1.694, 5.082) = 0.6524 | 0.5355 |
| INK128 | Concentration | F (1.247, 3.741) = 0.9388 | 0.4167 |
| | TCGA | F (1.419, 4.257) = 0.2389 | 0.7279 |
| | Concentration x TCGA | F (1.512, 4.537) = 0.9943 | 0.4119 |
| Lapatinib | Concentration | F (2.092, 6.276) = 0.1188 | 0.8973 |
| | TCGA | F (1.402, 4.206) = 0.7124 | 0.4929 |
| | Concentration x TCGA | F (2.286, 6.857) = 1.206 | 0.3633 |
| Lenalidomide | Concentration | F (1.739, 5.216) = 0.3843 | 0.6723 |
| | TCGA | F (1.277, 3.832) = 1.393 | 0.3233 |
| | Concentration x TCGA | F (1.781, 5.342) = 0.5419 | 0.5915 |
| Methoxyamine HCl | Concentration | F (2.597, 7.790) = 0.9584 | 0.4471 |
| | TCGA | F (1.430, 4.291) = 1.084 | 0.3869 |
| | Concentration x TCGA | F (1.963, 5.888) = 1.190 | 0.3669 |
| Olaparib | Concentration | F (1.809, 5.428) = 0.8882 | 0.4532 |
| | TCGA | F (1.317, 3.950) = 0.6649 | 0.5033 |
| | Concentration x TCGA | F (1.883, 5.649) = 0.6155 | 0.5640 |
| Onalespib lactate | Concentration | F (2.377, 7.131) = 0.3515 | 0.7481 |
| | TCGA | F (1.218, 3.655) = 1.636 | 0.2873 |
| | Concentration x TCGA | F (1.821, 5.464) = 0.5161 | 0.6076 |
| Pazopanib | Concentration | F (2.193, 6.580) = 0.6654 | 0.5585 |
| | TCGA | F (1.165, 3.496) = 1.017 | 0.3945 |
| | Concentration x TCGA | F (2.389, 7.166) = 0.8978 | 0.4663 |
| Pomalidomide | Concentration | F (2.506, 7.517) = 1.381 | 0.3160 |
| | TCGA | F (1.447, 4.342) = 0.2560 | 0.7198 |
| | Concentration x TCGA | F (2.372, 7.115) = 1.096 | 0.3962 |
| Rociletinib | Concentration | F (1.597, 4.790) = 1.804 | 0.2553 |

| | | | |
|---|---|---|---|
| | TCGA | F (1.146, 3.439) = 0.3530 | 0.6177 |
| | Concentration x TCGA | F (2.003, 6.010) = 0.5495 | 0.6040 |
| Romidepsin | Concentration | F (1.327, 3.981) = 2.104 | 0.2293 |
| | TCGA | F (1.551, 4.653) = 6.886 | **0.0441** |
| | Concentration x TCGA | F (1.590, 4.771) = 1.418 | 0.3173 |
| Selumetinib | Concentration | F (2.225, 6.675) = 0.9306 | 0.4501 |
| | TCGA | F (1.340, 4.019) = 0.02521 | 0.9324 |
| | Concentration x TCGA | F (1.961, 5.882) = 1.153 | 0.3766 |
| Sorafenib tosylate | Concentration | F (1.851, 5.554) = 2.270 | 0.1905 |
| | TCGA | F (1.288, 3.865) = 0.1873 | 0.7470 |
| | Concentration x TCGA | F (1.864, 5.591) = 2.225 | 0.1947 |
| Sunitinib malate | Concentration | F (1.785, 5.355) = 7.021 | **0.0335** |
| | TCGA | F (1.394, 4.183) = 3.631 | 0.1249 |
| | Concentration x TCGA | F (1.633, 4.898) = 1.633 | 0.2795 |
| Talazoparib | Concentration | F (1.046, 3.139) = 1.072 | 0.3784 |
| | TCGA | F (1.009, 3.026) = 3.483 | 0.1582 |
| | Concentration x TCGA | F (1.192, 3.577) = 2.126 | 0.2328 |
| Tazemetostat | Concentration | F (1.422, 4.267) = 2.920 | 0.1597 |
| | TCGA | F (1.179, 3.537) = 0.01089 | 0.9467 |
| | Concentration x TCGA | F (1.995, 5.984) = 2.683 | 0.1474 |
| Temsirolimus | Concentration | F (1.149, 3.448) = 1.993 | 0.2475 |
| | TCGA | F (1.013, 3.040) = 0.8526 | 0.4249 |
| | Concentration x TCGA | F (1.618, 4.854) = 1.525 | 0.2976 |
| Tivantinib | Concentration | F (1.232, 3.695) = 0.3906 | 0.6103 |
| | TCGA | F (1.028, 3.084) = 0.8000 | 0.4392 |
| | Concentration x TCGA | F (1.311, 3.932) = 1.158 | 0.3681 |

| | | | |
|---|---|---|---|
| Triapine | Concentration | F (1.575, 4.725) = 0.6210 | 0.5402 |
| | TCGA | F (1.020, 3.060) = 0.007156 | 0.9406 |
| | Concentration x TCGA | F (2.079, 6.237) = 0.8903 | 0.4609 |
| Veliparib | Concentration | F (1.739, 5.216) = 2.277 | 0.1945 |
| | TCGA | F (1.285, 3.854) = 1.270 | 0.3453 |
| | Concentration x TCGA | F (1.857, 5.571) = 2.484 | 0.1699 |
| Vismodegib | Concentration | F (1.447, 4.340) = 0.7402 | 0.4862 |
| | TCGA | F (1.498, 4.495) = 1.634 | 0.2821 |
| | Concentration x TCGA | F (1.739, 5.216) = 1.210 | 0.3609 |

**Supplementary Table 6.** Post-hoc analyses for selumetinib

| TCGA Subtype | Dunnett's multiple comparisons test | Adjusted P-value |
|---|---|---|
| Proneural | DMSO vs. 10 μM | 0.8325 |
| | DMSO vs. 3.33 μM | 0.9943 |
| | DMSO vs. 1.11 μM | 0.9884 |
| | DMSO vs. 370 nM | 0.9994 |
| | DMSO vs. 123 nM | 0.5918 |
| | DMSO vs. 41 nM | 0.4940 |
| | DMSO vs. 14 nM | 0.4607 |
| | DMSO vs. 4.5 nM | 0.6752 |
| | DMSO vs. 1.5 nM | 0.4613 |
| | DMSO vs. 508 pM | 0.7928 |
| Mesenchymal | DMSO vs. 10 μM | >0.9999 |
| | DMSO vs. 3.33 μM | 0.1036 |
| | DMSO vs. 1.11 μM | 0.7279 |
| | DMSO vs. 370 nM | 0.7294 |
| | DMSO vs. 123 nM | 0.7133 |
| | DMSO vs. 41 nM | 0.5797 |
| | DMSO vs. 14 nM | 0.7439 |
| | DMSO vs. 4.5 nM | 0.8420 |
| | DMSO vs. 1.5 nM | 0.8078 |
| | DMSO vs. 508 pM | 0.9797 |
| Classical | DMSO vs. 10 μM | **0.0423** |
| | DMSO vs. 3.33 μM | 0.0551 |

| | | |
|---|---|---|
| | DMSO vs. 1.11 μM | **0.0403** |
| | DMSO vs. 370 nM | 0.4321 |
| | DMSO vs. 123 nM | 0.7768 |
| | DMSO vs. 41 nM | 0.7817 |
| | DMSO vs. 14 nM | 0.4388 |
| | DMSO vs. 4.5 nM | >0.9999 |
| | DMSO vs. 1.5 nM | >0.9999 |
| | DMSO vs. 508 pM | 0.5397 |

**Supplementary Table 7.** Overall test for differences in stem cell frequencies between any of the groups, n=3-8/group.

| TCGA Subtype | Cell Line | Chisq | DF | P-value |
|---|---|---|---|---|
| Proneural | HK 217 | Inf | 8 | **0** |
| Mesenchymal | HK 336 | 707 | 8 | **1.91e-147** |
| Classical | HK 244 | 220 | 8 | **4.04e-43** |

**Supplementary Table 8.** Pairwise tests for differences in stem cell frequencies in Proneural cell line HK 217.

| Group 1 | Group 2 | Chisq | DF | Pr(>Chisq) |
|---|---|---|---|---|
| 10uM_Selum_0Gy | 10uM_Selum_4Gy | 27.6 | 1 | **1.47e-07** |
| 10uM_Selum_0Gy | 10uM_Selum_8Gy | 66.9 | 1 | **2.92e-16** |
| 10uM_Selum_0Gy | 1uM_Selum_0Gy | 9.63 | 1 | **0.00191** |
| 10uM_Selum_0Gy | 1uM_Selum_4Gy | 6.46 | 1 | **0.011** |
| 10uM_Selum_0Gy | 1uM_Selum_8Gy | 101 | 1 | **1.18e-23** |
| 10uM_Selum_0Gy | DMSO_0Gy | 6.53 | 1 | **0.0106** |
| 10uM_Selum_0Gy | DMSO_4Gy | 2.23 | 1 | 0.136 |
| 10uM_Selum_0Gy | DMSO_8Gy | 153 | 1 | **4.03e-35** |
| 10uM_Selum_4Gy | 10uM_Selum_8Gy | 10.2 | 1 | **0.00139** |
| 10uM_Selum_4Gy | 1uM_Selum_0Gy | 70.9 | 1 | **3.73e-17** |
| 10uM_Selum_4Gy | 1uM_Selum_4Gy | 8.29 | 1 | **0.00398** |
| 10uM_Selum_4Gy | 1uM_Selum_8Gy | 24.8 | 1 | **6.37e-07** |
| 10uM_Selum_4Gy | DMSO_0Gy | 80.6 | 1 | **2.82e-19** |
| 10uM_Selum_4Gy | DMSO_4Gy | 56.3 | 1 | **6.33e-14** |
| 10uM_Selum_4Gy | DMSO_8Gy | 49 | 1 | **2.53e-12** |
| 10uM_Selum_8Gy | 1uM_Selum_0Gy | 127 | 1 | **2.31e-29** |
| 10uM_Selum_8Gy | 1uM_Selum_4Gy | 36.1 | 1 | **1.86e-09** |
| 10uM_Selum_8Gy | 1uM_Selum_8Gy | 2.59 | 1 | 0.107 |
| 10uM_Selum_8Gy | DMSO_0Gy | 153 | 1 | **4.23e-35** |
| 10uM_Selum_8Gy | DMSO_4Gy | 116 | 1 | **5.06e-27** |
| 10uM_Selum_8Gy | DMSO_8Gy | 9.99 | 1 | **0.00157** |
| 1uM_Selum_0Gy | 1uM_Selum_4Gy | 33 | 1 | **9.01e-09** |
| 1uM_Selum_0Gy | 1uM_Selum_8Gy | 173 | 1 | **1.49e-39** |
| 1uM_Selum_0Gy | DMSO_0Gy | 1.02 | 1 | 0.312 |
| 1uM_Selum_0Gy | DMSO_4Gy | 3.63 | 1 | 0.0569 |
| 1uM_Selum_0Gy | DMSO_8Gy | 246 | 1 | **1.55e-55** |
| 1uM_Selum_4Gy | 1uM_Selum_8Gy | 62.4 | 1 | **2.86e-15** |
| 1uM_Selum_4Gy | DMSO_0Gy | 32.8 | 1 | **1.03e-08** |
| 1uM_Selum_4Gy | DMSO_4Gy | 19.6 | 1 | **9.68e-06** |
| 1uM_Selum_4Gy | DMSO_8Gy | 104 | 1 | **1.73e-24** |
| 1uM_Selum_8Gy | DMSO_0Gy | 219 | 1 | **1.25e-49** |
| 1uM_Selum_8Gy | DMSO_4Gy | 170 | 1 | **7.31e-39** |
| 1uM_Selum_8Gy | DMSO_8Gy | 1.97 | 1 | 0.161 |
| DMSO_0Gy | DMSO_4Gy | 1.23 | 1 | 0.267 |
| DMSO_0Gy | DMSO_8Gy | Inf | 1 | **0** |
| DMSO_4Gy | DMSO_8Gy | Inf | 1 | **0** |

**Supplementary Table 9.** Pairwise tests for differences in stem cell frequencies in Mesenchymal cell line HK 336.

| Group 1 | Group 2 | Chisq | DF | Pr(>Chisq) |
|---|---|---|---|---|
| 10uM_Selum_0Gy | 10uM_Selum_4Gy | 62.1 | 1 | **3.24e-15** |
| 10uM_Selum_0Gy | 10uM_Selum_8Gy | 109 | 1 | **2.01e-25** |
| 10uM_Selum_0Gy | 1uM_Selum_0Gy | 0.0499 | 1 | 0.823 |
| 10uM_Selum_0Gy | 1uM_Selum_4Gy | 5.33 | 1 | 0.021 |
| 10uM_Selum_0Gy | 1uM_Selum_8Gy | 132 | 1 | 1.22e-30 |
| 10uM_Selum_0Gy | DMSO_0Gy | 21.3 | 1 | 3.96e-06 |
| 10uM_Selum_0Gy | DMSO_4Gy | 43.4 | 1 | 4.39e-11 |
| 10uM_Selum_0Gy | DMSO_8Gy | 130 | 1 | 4.71e-30 |
| 10uM_Selum_4Gy | 10uM_Selum_8Gy | 6.63 | 1 | 0.01 |
| 10uM_Selum_4Gy | 1uM_Selum_0Gy | 77.9 | 1 | 1.11e-18 |
| 10uM_Selum_4Gy | 1uM_Selum_4Gy | 30 | 1 | 4.36e-08 |
| 10uM_Selum_4Gy | 1uM_Selum_8Gy | 12.5 | 1 | 0.000402 |
| 10uM_Selum_4Gy | DMSO_0Gy | 208 | 1 | 3.48e-47 |
| 10uM_Selum_4Gy | DMSO_4Gy | 6.75 | 1 | 0.0094 |
| 10uM_Selum_4Gy | DMSO_8Gy | 9.21 | 1 | 0.00241 |
| 10uM_Selum_8Gy | 1uM_Selum_0Gy | 134 | 1 | 5.9e-31 |
| 10uM_Selum_8Gy | 1uM_Selum_4Gy | 63.8 | 1 | 1.36e-15 |
| 10uM_Selum_8Gy | 1uM_Selum_8Gy | 0.802 | 1 | 0.37 |
| 10uM_Selum_8Gy | DMSO_0Gy | 317 | 1 | **5.55e-71** |
| 10uM_Selum_8Gy | DMSO_4Gy | 35.3 | 1 | **2.85e-09** |
| 10uM_Selum_8Gy | DMSO_8Gy | 0.0221 | 1 | 0.882 |
| 1uM_Selum_0Gy | 1uM_Selum_4Gy | 7.4 | 1 | **0.00653** |
| 1uM_Selum_0Gy | 1uM_Selum_8Gy | 163 | 1 | **2.73e-37** |
| 1uM_Selum_0Gy | DMSO_0Gy | 22.8 | 1 | **1.8e-06** |
| 1uM_Selum_0Gy | DMSO_4Gy | 57.5 | 1 | **3.42e-14** |
| 1uM_Selum_0Gy | DMSO_8Gy | 161 | 1 | **5.59e-37** |
| 1uM_Selum_4Gy | 1uM_Selum_8Gy | 81.5 | 1 | **1.72e-19** |
| 1uM_Selum_4Gy | DMSO_0Gy | 53.2 | 1 | **3.01e-13** |
| 1uM_Selum_4Gy | DMSO_4Gy | 15.4 | 1 | **8.76e-05** |
| 1uM_Selum_4Gy | DMSO_8Gy | 77.8 | 1 | **1.12e-18** |
| 1uM_Selum_8Gy | DMSO_0Gy | 379 | 1 | **2.36e-84** |
| 1uM_Selum_8Gy | DMSO_4Gy | 55 | 1 | **1.23e-13** |
| 1uM_Selum_8Gy | DMSO_8Gy | 0.803 | 1 | 0.37 |
| DMSO_0Gy | DMSO_4Gy | 188 | 1 | **7.49e-43** |
| DMSO_0Gy | DMSO_8Gy | 390 | 1 | **1.04e-86** |
| DMSO_4Gy | DMSO_8Gy | 51.9 | 1 | **5.96e-1** |

**Supplementary Table 8.** Pairwise tests for differences in stem cell frequencies in Classical cell line HK 244.

| Group 1 | Group 2 | Chisq | DF | Pr(>Chisq) |
|---|---|---|---|---|
| 10uM_Selum_0Gy | 10uM_Selum_4Gy | 29.1 | 1 | **6.87e-08** |
| 10uM_Selum_0Gy | 10uM_Selum_8Gy | 5.74 | 1 | **0.0166** |
| 10uM_Selum_0Gy | 1uM_Selum_0Gy | 10.6 | 1 | **0.00111** |
| 10uM_Selum_0Gy | 1uM_Selum_4Gy | 9.85 | 1 | **0.0017** |
| 10uM_Selum_0Gy | 1uM_Selum_8Gy | 32.2 | 1 | **1.41e-08** |
| 10uM_Selum_0Gy | DMSO_0Gy | 22 | 1 | **2.72e-06** |
| 10uM_Selum_0Gy | DMSO_4Gy | 1.3 | 1 | 0.254 |
| 10uM_Selum_0Gy | DMSO_8Gy | 0.14 | 1 | 0.708 |
| 10uM_Selum_4Gy | 10uM_Selum_8Gy | 8.31 | 1 | **0.00395** |
| 10uM_Selum_4Gy | 1uM_Selum_0Gy | 55.5 | 1 | **9.22e-14** |
| 10uM_Selum_4Gy | 1uM_Selum_4Gy | 50.2 | 1 | **1.4e-12** |
| 10uM_Selum_4Gy | 1uM_Selum_8Gy | 1.04 | 1 | 0.307 |
| 10uM_Selum_4Gy | DMSO_0Gy | 120 | 1 | **5.63e-28** |
| 10uM_Selum_4Gy | DMSO_4Gy | 45.2 | 1 | **1.77e-11** |
| 10uM_Selum_4Gy | DMSO_8Gy | 37.7 | 1 | **8.15e-10** |
| 10uM_Selum_8Gy | 1uM_Selum_0Gy | 26.2 | 1 | **3.13e-07** |
| 10uM_Selum_8Gy | 1uM_Selum_4Gy | 23.8 | 1 | **1.07e-06** |
| 10uM_Selum_8Gy | 1uM_Selum_8Gy | 12.1 | 1 | **0.000508** |
| 10uM_Selum_8Gy | DMSO_0Gy | 53.9 | 1 | **2.08e-13** |
| 10uM_Selum_8Gy | DMSO_4Gy | 12.8 | 1 | **0.00034** |
| 10uM_Selum_8Gy | DMSO_8Gy | 8.47 | 1 | **0.00361** |
| 1uM_Selum_0Gy | 1uM_Selum_4Gy | 0.0121 | 1 | 0.912 |
| 1uM_Selum_0Gy | 1uM_Selum_8Gy | 58.2 | 1 | **2.32e-14** |
| 1uM_Selum_0Gy | DMSO_0Gy | 0.00554 | 1 | 0.941 |
| 1uM_Selum_0Gy | DMSO_4Gy | 5.77 | 1 | **0.0163** |
| 1uM_Selum_0Gy | DMSO_8Gy | 9.49 | 1 | **0.00207** |
| 1uM_Selum_4Gy | 1uM_Selum_8Gy | 53.4 | 1 | **2.76e-13** |
| 1uM_Selum_4Gy | DMSO_0Gy | 0.00418 | 1 | 0.948 |
| 1uM_Selum_4Gy | DMSO_4Gy | 5.47 | 1 | **0.0194** |
| 1uM_Selum_4Gy | DMSO_8Gy | 8.77 | 1 | **0.00307** |
| 1uM_Selum_8Gy | DMSO_0Gy | 110 | 1 | **1.02e-25** |
| 1uM_Selum_8Gy | DMSO_4Gy | 46.6 | 1 | **8.74e-12** |
| 1uM_Selum_8Gy | DMSO_8Gy | 39.6 | 1 | **3.06e-10** |
| DMSO_0Gy | DMSO_4Gy | 12.1 | 1 | **0.000503** |
| DMSO_0Gy | DMSO_8Gy | 20.9 | 1 | **4.83e-06** |
| DMSO_4Gy | DMSO_8Gy | 0.682 | 1 | 0.409 |

**Supplementary Table 9.** Goodness of fit tests.

| Cell Line | Estimated Slope | Test | Chisq | DF | P-Value |
|---|---|---|---|---|---|
| HK 217 | 0.619 | Likelihood ratio test of single-hit model | 119 | 1 | **1.24e-27** |
| | | Score test of heterogeneity | 7.56e-05 | 1 | 0.993 |
| HK 336 | 0.627 | Likelihood ratio test of single-hit model | 93 | 1 | **5.23e-22** |
| | | Score test of heterogeneity | 2.43e-12 | 1 | 1 |
| HK 244 | 0.728 | Likelihood ratio test of single-hit model | 41.7 | 1 | **1.08e-10** |
| | | Score test of heterogeneity | 1.58e-27 | 1 | 1 |
